# Pathway Analysis within Multiple Human Ancestries Reveals Novel Signals for Epistasis in Complex Traits

**DOI:** 10.1101/2020.09.24.312421

**Authors:** Michael C. Turchin, Gregory Darnell, Lorin Crawford, Sohini Ramachandran

## Abstract

Genome-wide association (GWA) studies have identified thousands of significant genetic associations in humans across a number of complex traits. However, the majority of these studies focus on linear additive relationships between genotypic and phenotypic variation. Epistasis, or non-additive genetic interactions, has been identified as a major driver of both complex trait architecture and evolution in multiple model organisms; yet, this same phenomenon is not considered to be a significant factor underlying human complex traits. There are two possible reasons for this assumption. First, most large GWA studies are conducted solely with European cohorts; therefore, our understanding of broad-sense heritability for many complex traits is limited to just one ancestry group. Second, current epistasis mapping methods commonly identify significant genetic interactions by exhaustively searching across all possible pairs of SNPs. In these frameworks, estimated epistatic effects size are often small and power can be low due to the multiple testing burden. Here, we present a case study that uses a novel region-based mapping approach to analyze sets of variants for the presence of epistatic effects across six diverse subgroups within the UK Biobank. We refer to this method as the “MArginal ePIstasis Test for Regions” or MAPIT-R. Even with limited sample sizes, we find a total of 245 pathways within the KEGG and REACTOME databases that are significantly enriched for epistatic effects in height and body mass index (BMI), with 67% of these pathways being detected within individuals of African ancestry. As a secondary analysis, we introduce a novel region-based “leave-one-out” approach to localize pathway-level epistatic signals to specific interacting genes in BMI. Overall, our results indicate that non-European ancestry populations may be better suited for the discovery of non-additive genetic variation in human complex traits — further underscoring the need for publicly available, biobank-sized datasets of diverse groups of individuals.

## Introduction

Genome-wide association (GWA) studies are a powerful tool for understanding the genetic architecture of complex traits and phenotypes [1–8]. The most common approach for conducting GWA studies is to use a linear mixed model to test for statistical associations between individual genetic variants and a phenotype of interest; here, the estimated regression coefficients represent an additive relationship between number of copies of a single-nucleotide polymorphism (SNP) and the phenotypic state. While this approach has produced many statistically significant additive associations, it is less amenable to detecting nonlinear genetic associations that also contribute to a trait’s genetic architecture. Epistasis, commonly defined as the nonlinear, or non-additive, interaction between multiple genetic variants, is a well-established phenomenon in a number of model organisms [9–18]. Epistasis has also been suggested as a major driver of both phenotypic variation and evolution [19–26]. Still, there remains skepticism and controversy regarding the importance of epistasis in human complex traits and diseases [27–34]. For example, multiple studies have suggested that phenotypic variation can be mainly explained with additive effects [27, 28, 32]; although, this hypothesis has been been challenged recently [35]. In initial work to locate the “missing heritability” in the human genome — the discrepancy between larger pedigree-based trait heritability estimates and smaller SNP-based trait heritability estimates using the first wave of human GWA study results [36–38] — it was suggested that epistasis may account for a significant portion of this observed discrepancy [24, 39, 40]. However, other studies have posited that, for at least some human phenotypes, genetic interactions are unlikely to be a major contributor to total heritability [34, 41, 42].

Algorithmically, detecting statistically significant epistatic signals via genome-wide scans is much more computationally expensive than the the traditional hypothesis-generating GWA framework. GWA tests for additive effects are linear in the number of SNPs, while epistasis scans usually consider, at a minimum, all pairwise combinations of SNPs (e.g., a total of *J* (*J* − 1)/2 possible pairwise combinations for *J* variants in a study). Methods that fall within the MArginal ePIstasis Test (MAPIT) framework [43–46] attempt to address these challenges by alternatively testing for *marginal* epistasis. Specifically, instead of directly identifying individual pairwise or higher-order interactions, these approaches focus on identifying variants that have a non-zero interaction effect with any other variant in the data. Indeed, analyzing epistasis among pairs of SNPs can be underpowered in GWA studies, particularly when applied to polygenic traits or traits which are generated by many mutations of small effect [4, 47–49].

To overcome this limitation, more recent computational approaches have expanded the additive GWA framework to aggregate across multiple SNP-level association signals and test for the enrichment of genes and pathways [50–61]. In Nakka et al. [62] we showed that enrichment analyses applied to multiple ancestries can identify genes and gene networks contributing to disease risk that ancestry-specific enrichment analyses fail to find. Recent multiethnic GWA studies have also found that using non-European populations offer new insights into additive genetic architecture [63–70]. However, despite this growing body of work and increasing efforts to promote conducting GWA studies in diverse ancestries [68, 71–75], few studies have investigated the role of epistasis in shaping multiethnic human genetic variation (but see [76–79]). Expanding epistasis studies to include non-European ancestries, as well as to aggregate over multiple SNP-level signals, may reveal a new understanding of non-additive genetic architecture in human complex traits.

In this study, our objective is to expand the marginal epistasis framework from individual SNPs to user-specified sets of variants (e.g., genes, signaling pathways) and apply the framework to multiple, diverse human ancestries. We aim to detect novel interactions between biologically relevant disease mechanisms underlying complex traits and to analyze multiple human ancestries, all while reducing the multiple testing burden that traditionally hinders exhaustive epistatic scans. We implement our new approach in “MArginal ePIstasis Test for Regions”, which we refer to as MAPIT-R. We apply MAPIT-R using pathway annotations from the “Kyoto Encyclopedia of Genes and Genomes” (KEGG) and REACTOME databases [80] to standing height and body mass index (BMI) assayed in individuals from multiple human ancestry “subgroups” (British, African, Caribbean, Chinese, Indian, and Pakistani) in the UK Biobank [81]. Spanning across all these subgroups, we find more than 200 pathways that have significant marginal epistatic effects on standing height and BMI. We then investigate the distribution of these significant non-additive signals across ancestries, traits, and pathways, finding future directions to prioritize for studies of epistasis in human complex traits.

## Materials and Methods

### Overview of the MAPIT-R Model

We describe the intuition behind the “MArginal ePIstasis Test for Regions” (MAPIT-R) in detail here. Consider a genome-wide association (GWA) study with *N* individuals. Within this study, we assume that we have an *N*-dimensional vector of quantitative traits **y**, an *N × J* matrix of genotypes **X**, with *J* denoting the number of single nucleotide polymorphisms (SNPs) encoded as {0,1, 2} copies of a reference allele at each locus, and a list of *L* predefined genomic regions of interests 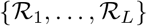. We will let each genomic region *l* represent a known collection of annotated SNPs 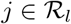 with set cardinality 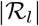. In this work, each 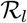 includes sets of SNPs that fall within functional regions of genes that have been annotated as being members of certain pathways or gene sets (see Supplementary Note). Recall that our objective is to test whether a set of biologically relevant variants have a nonzero interaction effect with any other region along the genome. Therefore, MAPIT-R works by examining one region at a time (indexed by l) and fits the following linear mixed model

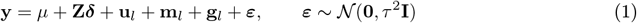

where *μ* is an intercept term; **Z** is a matrix of covariates (e.g., the top principal components from the genotype matrix) with coefficients ***δ***; 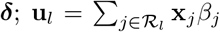 is the summation of region-specific effects with corresponding additive effect sizes *β_j_* for the *j*-th variant; **x**_*j*_ is an *N*-dimensional genotypic vector for the *j*-th variant in the *l*-th region that is the focus of the model; 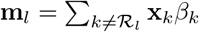 is the combined additive effects from all other 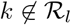 SNPs in the data that have not been annotated as being within the 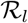 region of interest with coefficients *β_k_*; **x**_*k*_ is an *N*-dimensional genotypic vector for the *k*-th variant in the data that has not been annotated as being within the 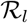 region of interest; 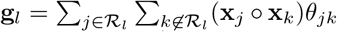 is the summation of all pairwise interaction effects (i.e., the Hadamard product **X**_*j*_ ○ **x**_*k*_) between the *j*-th variant in the *l*-th annotated region 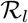 and all other *k ≠ j* variants outside of 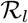 with corresponding coefficients *θ_jk_*; and ***ε*** is a normally distributed error term with mean zero and independent residual error variance scaled by the component *τ*^2^. There are a few important takeaways from this formulation of MAPIT-R. First, the term **m**_*l*_ effectively represents the polygenic background of all variants except for those that have been annotated for the *l*-th region of interest. Second, and most importantly, the term **g**_*l*_ is the main focus of the model and represents the *marginal epistatic* effect of the region **R**_*l*_ [43, 44]. It is important to note that each component of the model will change with every new region that is considered.

For convenience, we assume that both the genotype matrix (column-wise) and the trait of interest have been mean-centered and standardized to have unit variance. Next, because the model in Eq. (1) is an underdetermined linear system (*J > N*), we ensure identifiability by assuming that the individual regression coefficients follow univariate normal distributions where

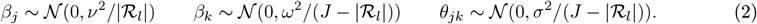

With the assumption of normally distributed effect sizes, the MAPIT-R model defined in Eq. (1) becomes a multiple variance component model where 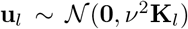 with 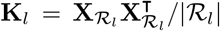 being the genetic relatedness matrix computed using genotypes from all variants within the region of interest; 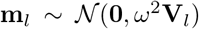 with 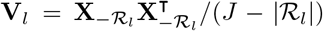 being the genetic relatedness matrix computed using genotypes outside the region of interest; and 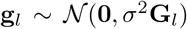 with **G**_*l*_ = **K**_*l*_ ○ **V**_*l*_ representing a second-order interaction relationship matrix which is obtained by using the Hadamard product (i.e., the squaring of each element) between the region-specific relatedness matrix and its corresponding polygenic background. Importantly, the variance component *σ*^2^ effectively captures the marginal epistatic effect for the *l*-th region. Even though we limit ourselves to the task of identifying second order (i.e., pairwise) epistatic relationships between sets of SNPs in this paper, extensions to higher-order and gene-by-environmental interactions are straightforward to implement for alternative analyses [43, 45, 82–84].

### Hypothesis Testing with the MAPIT-R Framework

In this section, we now describe how to perform joint estimation of all the variance component parameters in the MAPIT-R model. Since our goal is to identify genomic regions that have significant non-zero interaction effects on a given phenotype, we examine each annotated SNP-set *l* = 1,…, *L* in turn, and test the null hypothesis in Eq. (1) and Eq. (2) that *H*_0_: *σ*^2^ = 0. We make use of the MQS method for parameter estimation and hypothesis testing [83]. Briefly, MQS is based on the computationally efficient method of moments and produces estimates that are mathematically identical to the Haseman-Elston (HE) cross-product regression [85]. To estimate the variance components with MQS, we first regress out the additive effects of the *l*-th SNP-set, the fixed covariates, and the intercept terms. Equivalently, we multiply both sides of Eq. (1) by a projection (hat) matrix such that the model becomes orthogonal to the column space of the intercept term *μ*. Specifically, we define **H = I − B(B^⊤^B)^−1^B^⊤^** where 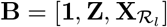 is a concatenated matrix and with **1** being an *N*-dimensional vector of ones. This yields a simplified model

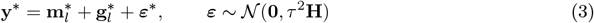

where **y* = Hy** is the projected phenotype of interest; 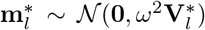 with 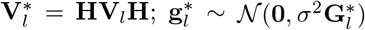 with 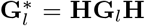; and ***ε*\* = H*ε*** is the projected residual error, respectively. Then lastly, for each annotation considered, the MQS estimate for the marginal epistatic effect is computed as

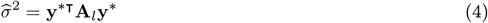

where 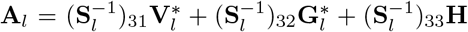 with elements (**S**_*l*_)_*jk*_ = tr(**∑**_*lj*_ **∑**_*lk*_) for the covariance matrices subscripted as 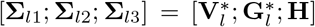. Here, tr(**●**) is used to denote the matrix trace function. It has been well established that the marginal variance component estimate 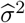 follows a mixture of chi-square distributions under the null hypothesis because of its quadratic form and the assumed normally distributed trait **y** [43, 53, 86–89]. Namely, 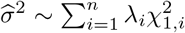, where 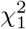 are chi-square random variables with one degree of freedom and (λ_1_,…, λ_n_) are the eigenvalues of the matrix [43, 83]

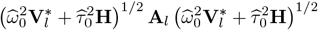

with 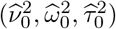 being the MQS estimates of (*ν*^2^, *ω*^2^, *τ*^2^) under the null hypothesis. Several approximation and exact methods have been suggested to obtain *p*-values under the distribution of 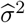. In this paper, we use the Davies exact method [87, 90].

### Software Availability

Code for implementing the “MArginal ePIstasis Test for Regions” (MAPIT-R) is freely available in R/Rcpp and is located at https://cran.r-project.org/web/packages/MAPITR/index.html. All MAPIT-R functions use the `CompQuadForm R` package to compute *p*-values with the Davies method. Note that the Davies method can sometimes yield a *p*-value that exactly equals 0. This can occur when the true *p*-value is extremely small [91]. In this case, we report *p*-values as being truncated at 1 × 10^−10^. Alternatively, one could also compute *p*-values for all MAPIT-R based functions using Kuonen’s saddlepoint method [91, 92] or Satterthwaite’s approximation equation [93].

### SNP-Set and Pathway Annotations

To create appropriate pathway annotations for MAPIT-R, we first assign SNPs to genes and then aggregate the genes together according to pathway definitions provided by the KEGG and REACTOME databases, respectively. KEGG and REACTOME pathway definitions were downloaded and extracted from the Broad Institute’s Molecular Signatures Database (MSigDB; https://www.gsea-msigdb.org/gsea/msigdb/collections.jsp#C2) under the collection “C2: Curated Gene Sets” [80]. SNPs were annotated using Annovar [94] and were then mapped to a given gene if they were exonic, intronic, in the 5’ and 3’ UTRs, or within 20kb upstream or downstream of the gene.

### UK Biobank Data

To create the UK Biobank population subgroups used in this study (UK Biobank Application Number 2241), we first extracted and grouped individuals by the self-identified ancestries of “African”, “British”, “Caribbean”, “Chinese”, “Indian”, and “Pakistani”. For the British subgroups, five sets of N = 4,000 and 10,000 non-overlapping individuals were created — with one set from each sample size being used for “primary analyses” and the remaining four being used for the “replication analyses”. Standard quality control procedures were applied to each population subgroup (see Supplementary Note for details). “Local” principal component analysis (PCA) was conducted to confirm ancestry groupings and to remove outliers. We refer to conducting PCA on each subgroup separately as “local” PCA to help distinguish from the alternative setup of conducting PCA on the entire dataset jointly, which we refer to as “global” PCA (see Supplementary Figure 1). Note that the genetic data we used in this study were the directly genotyped variant sets from the UK Biobank after running imputation of missing genotypes on the University of Michigan Imputation Server [95]. Here, imputation was conducted manually with an ancestry-diverse and sample-size balanced reference panel (1000G Phase 3 v5). For details on the final UK Biobank dataset, see Supplementary Tables 1 and 2. Lastly, both the standing height and body mass index (BMI) traits were adjusted for age, gender, and assessment center. Following previous pipelines [33, 96], each dataset was first divided into males and females. Age was then regressed out within each sex, and the resulting residuals were inverse normalized. These normalized values were then combined back together and assessment center designations were regressed out. Top 10 “local” principal components (PCs) were included as covariates during the actual MAPIT-R analyses. In total we conducted 24 different analyses (2 pathway databases, 2 phenotypes, 6 population subgroups), which we refer to as ‘database-phenotype-subgroup’ combinations. Lastly, for analyses using permuted phenotypes, permutations were conducted within-subgroup and done by randomly reassigning phenotypes to individuals.

**Figure 1.**
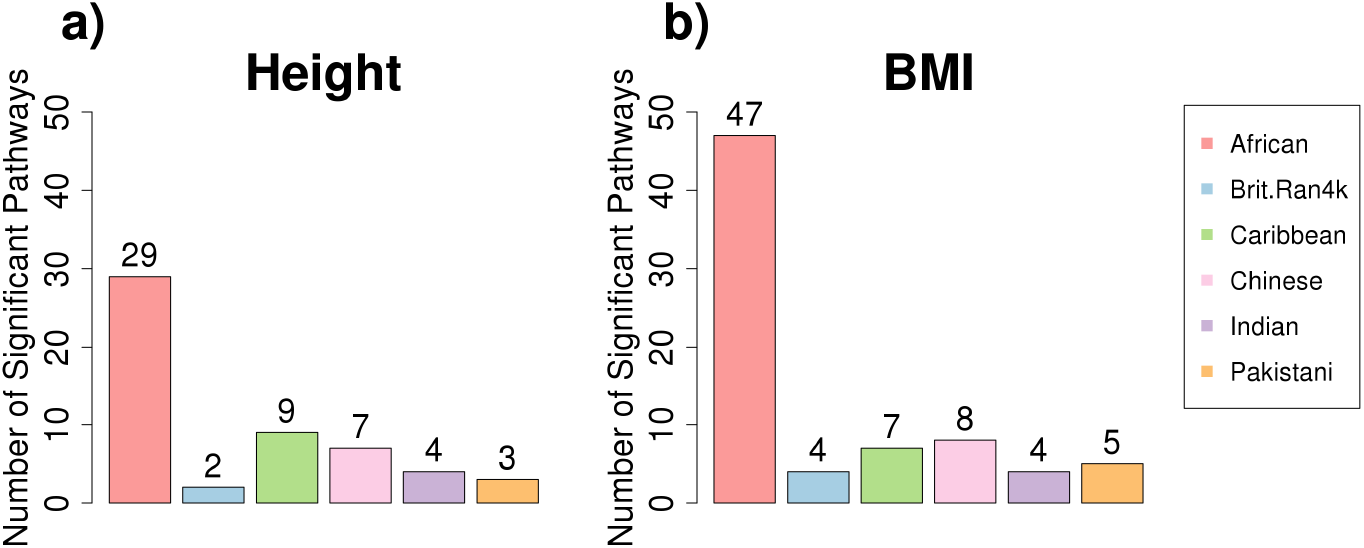
Number of KEGG pathways identified by MAPIT-R that have significant marginal epistatic effects within (a) standing height and (b) body mass index (BMI) per subgroup in the UK Biobank. Here, subgroups in the UK Biobank included individuals based on their self-identified ancestries: “African”, “British”, “Caribbean”, “Chinese”, “Indian”, and “Pakistani” (see legend to the right of panel **(b)**). Genome-wide significance was determined by using Bonferroni-corrected *p*-value thresholds based on the number of pathways tested in each database-phenotype-subgroup combination (see Supplementary Table 1). Across all database-phenotype combinations, the African subgroup has the largest numbers of significant pathways. For lists of the specific significant pathways per database-phenotype-subgroup combination, see Supplementary Table 3. Results from running MAPIT-R with REACTOME database pathways can be found in Supplementary Figure 2.

## Results

### Multiethnic Analyses Enables the Detection of Pathway-Level Interactions

We applied MAPIT-R to height and body mass index (BMI) to detect pathways from the KEGG and REACTOME databases [80] with significant epistatic interactions with other regions on the genome, using genotype data and diverse individuals from the UK Biobank. We focused on height and BMI due to the extensive work that has already been done investigating the broad-sense and narrow-sense heritabilities of these traits [29, 41, 97–100], and we used the KEGG and REACTOME databases because they cover an extensive range of both biological processes and pathway-sizes (measured in SNP counts). We analyzed six different human ancestry subgroups that we extracted from the UK Biobank: African (*N* = 3111), British (*N* = 3848, chosen randomly from the full *N* = 472,218 cohort), Caribbean (*N* = 3833), Chinese (*N* = 1448), Indian (*N* = 5077), and Pakistani (*N* = 1581) (Supplementary Figure 1 and Supplementary Tables 1-2). Subgroups were extracted based on self-identified ancestry and individuals were filtered using standard quality control procedures (see Materials and Methods and Supplementary Note for details). In total, we conducted 24 different analyses (i.e., 2 pathway databases, 2 phenotypes, 6 population subgroups), which we refer to as ‘database-phenotype-subgroup’ combinations.

Applying MAPIT-R to height and BMI within each ancestry subgroups, we find a total of 245 enriched pathways that have genome-wide significant signals for marginal epistatic interactions with the rest of the genome (Figure 1, Supplementary Figure 2, and Supplementary Table 3) Here, *p*-value significance thresholds were determined by using Bonferroni correction based on the number of pathways tested per analysis (see Supplementary Table 1). Overall, a similar number of pathways were statistically enriched between the KEGG and REACTOME databases (130 and 115, respectively); however, we find that BMI yields more non-additive genetic signal than height (155 versus 90 significant pathways, respectively). Across each ancestry-specific subgroup, our findings overlap with results from other work showing evidence for the importance of epistasis in human immunity, particularly involving the Major Histocompatibility Complex (MHC) [101–107], as well as the key roles metabolic processes and cellular signaling play in trait architecture for model systems [108–114]. Most notably, however, the majority of our results occurred within the African subgroup: 165 out of 245 significant pathways across all analyses.

Focusing on the African subgroup, the enriched pathways represent multiple biologically relevant themes in both height and BMI (Table 1 and Supplementary Table 3). When analyzing height with annotations from the KEGG database, we find that most of the statistically significant marginal epistatic interactions occur in pathways related to canonical signaling cascades, functions within the immune system, and sets of genes that affect heart conditions. Previous multiethnic GWA studies of height have found additive associations with cytokine genes [115] and WNT/beta-catenin signaling [116]. Results from MAPIT-R suggest that non-additive interactions involving cytokine receptors (*p*-value = 2.84 × 10^−8^) and genes within the WNT-signaling pathway (*p*-value = 6.54 × 10^−6^) also contribute to the complex genetic architecture of height as well. In BMI, we find similar themes, as well as multiple statistically significant signals from metabolic pathways (Table 1). Notably, MAPIT-R identified pathways related to ErbB signaling (*p*-value = 3.30 × 10^−7^) and ether lipid metabolism (*p*-value = 1.41 × 10^−4^) as having significant marginal epistatic effects — both of which have also been shown to have additive associations with BMI as well [96, 117, 118].

**Table 1.**
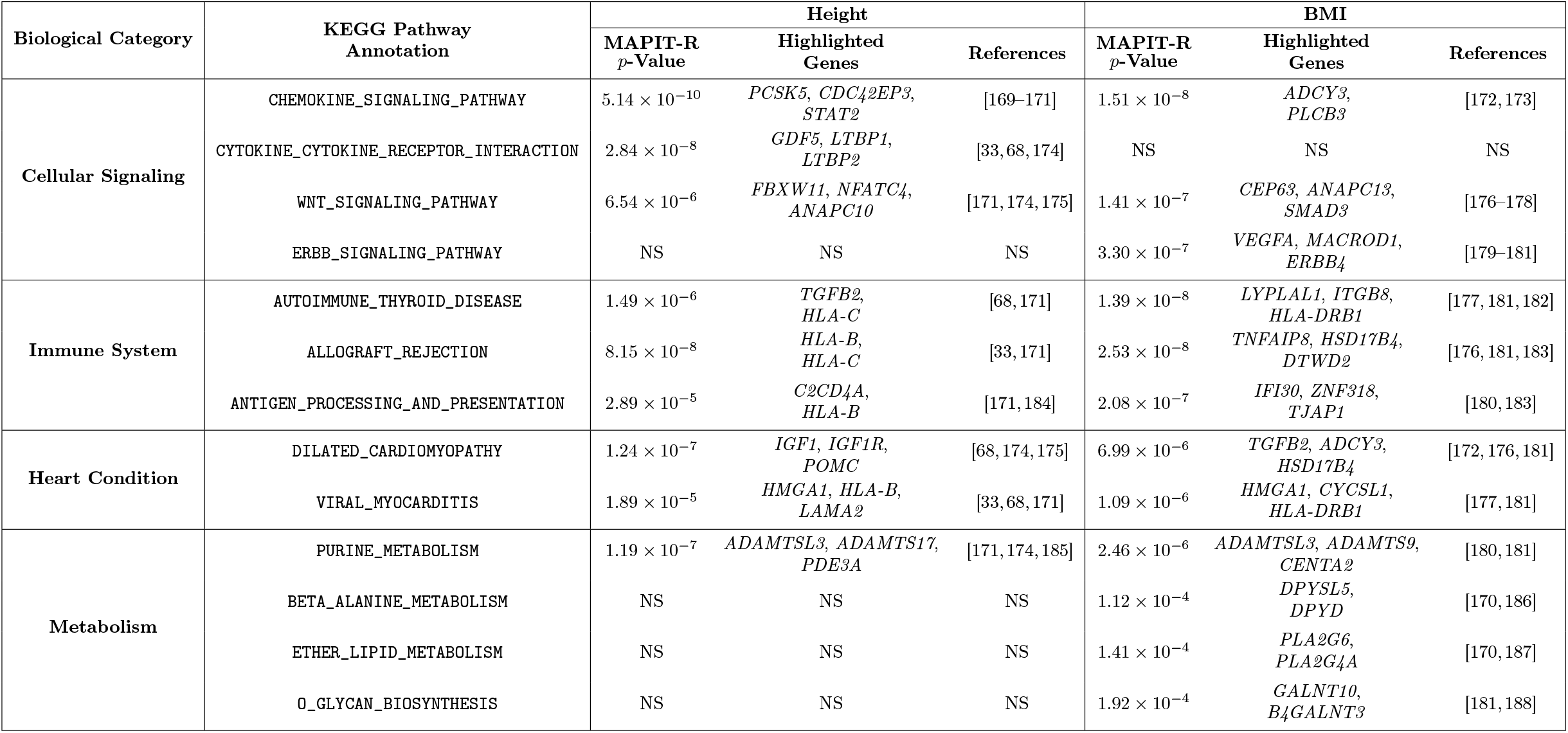
Biological themes among the MAPIT-R significant KEGG pathways for height and body mass index (BMI) within the African subgroup in the UK Biobank. The biological themes include: cellular signaling, immune system, heart condition, and metabolism. Notably enriched pathways for each biological theme are included in the second column. For each pathway, MAPIT-R *p*-values, highlighted gene associations, and references for each gene association are shown for both height (third, forth, and fifth columns) and BMI (sixth, seventh, and eighth columns). Genome-wide significance was determined by using Bonferroni-corrected *p*-value thresholds based on the number of pathways tested in each database-phenotype-subgroup combination (Supplementary Table 1). “Highlighted Genes” and “References” were determined using relevant SNP association citations from the GWAS Catalog (version 1.0.2) [7]. For a full list of MAPIT-R significant pathways in all database-phenotype-subgroup combinations, see Supplementary Table 3. NS indicates that a pathway was not genome-wide significant for a given phenotype.

It is important to note that, in our analyses, the African subgroup has neither the largest sample size nor the largest number of SNPs following quality control (Supplementary Table 1). Thus, to investigate the power of MAPIT-R and its sensitivity to underlying parameters, we conducted simulation studies under a range of genetic architectures (Supplementary Figure 3) [43]. Here, we found that MAPIT-R both controls type 1 error accurately and also has the power to effectively detect pathway level marginal epistasis, even for polygenic traits where the contribution from individual SNPs to the broad-sense heritability of a trait can be quite low. We also ran versions of MAPIT-R on the real data, but with permuted phenotypes, to ensure that the model was not identifying significant non-additive genetic relationships by chance (Supplementary Figures 4 and 5). These permutations allowed us to further investigate MAPIT-R’s false discovery rates, in which we observe values only as high as 1.5% across our different database-phenotype-subgroup combinations at multiple significance thresholds (Supplementary Table 4).

### Evidence of Epistasis within the Non-African Subgroups

In our analyses of the British, Chinese, Caribbean, Indian, and Pakistani subgroups, we identify 80 pathways in total that have significant marginal epistatic interactions. Interestingly, many of these pathways overlap with the set of significant results from the African subgroup; there is notably less overlap though in results between each of the individual non-African subgroups (Figure 2 and Supplementary Figure 6). For example, in the height analysis with KEGG annotations, 6-out-of-7 and 7-out-of-8 enriched pathways identified using the Caribbean and Chinese subgroups overlap with those detected while using the African subgroup, respectively. However, there is no overlap in results from our marginal epistasis scans at the pathway level between the Chinese and Caribbean subgroups.

**Figure 2.**
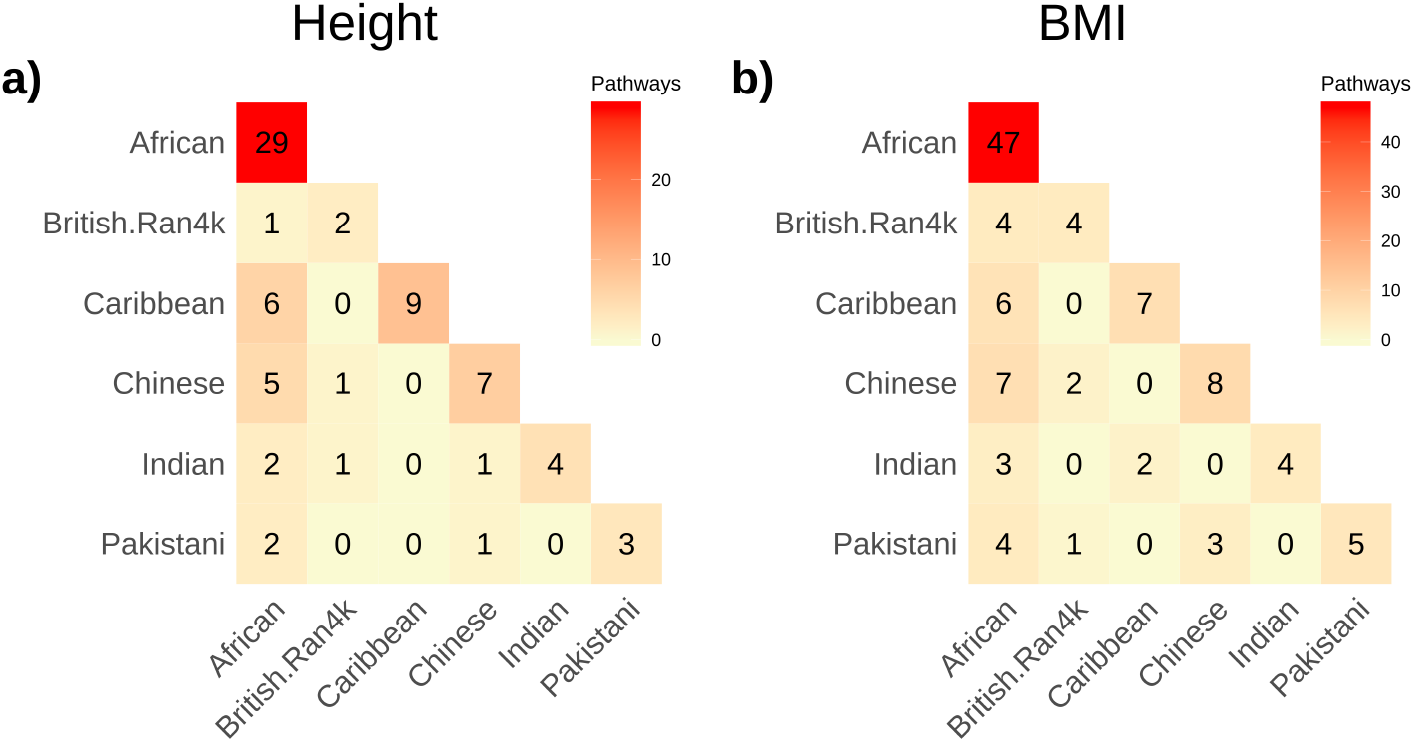
Heatmaps depicting the overlap of MAPIT-R significant KEGG pathways for (a) standing height and (b) body mass index (BMI) across the different ancestry-specific subgroups in the UK Biobank. Here, subgroups in the UK Biobank included individuals based on their self-identified ancestries: “African”, “British”, “Caribbean”, “Chinese”, “Indian”, and “Pakistani” (ordered here from top-to-bottom and left-to-right). Genome-wide significance was determined by using Bonferroni-corrected *p*-value thresholds based on the number of pathways tested in each databasephenotype-subgroup combination (see Supplementary Table 1). The diagonal shows the total number of genome-wide significant pathways per subgroup. We observe that significant pathways identified in non-African subgroups overlap more often with pathways from the African subgroup than they do with pathways from the other, remaining non-African subgroups. Results for both phenotypes in the REACTOME database can be seen in Supplementary Figure 6.

The pathways commonly identified with significant marginal epistatic signals in both the African and Caribbean subgroups contain genes related to multiple kinases (e.g., *MAPK1, ROCK1, PRKCB, PAK1*) and calcium channel proteins (e.g., *CACNA1S, CACNA1D*) (Supplementary Tables 5 and 6) — many of which are supported by associations validated in previous GWA applications [33, 119]. In contrast, the pathways with significant marginal epistatic effects identified in both the African and Chinese subgroups are pathways related to the immune system and contain multiple HLA loci (e.g., *HLA-DRA, HLA-DRB1, HLA-A, HLA-B*) (Supplementary Tables 5 and 6). These results are unsurprising since it is well known that the MHC region holds significant clinical relevance in complex traits [44, 103, 104, 120]; however, more recent work has also suggested that Han Chinese genomes may be particularly enriched for interactions involving HLA loci [121].

### Stronger Epistatic Signals underlie BMI than Height

In our analyses with the African subgroup, we detected far more significantly enriched pathways for BMI than in height while using both the KEGG and REACTOME database annotation (Figure 1 and Supplementary Figure 2). While there is considerable correlation between the MAPIT-R *p*-values in height and BMI (Pearson correlation coefficient *r* = 0.76 in KEGG and 0.72 in REACTOME, respectively), there are stronger marginal epistatic signals in BMI that remain significant after Bonferroni-correction (Figure 3). These results align with pedigree-based heritability estimates for each trait, which have indicated narrow-sense heritability is around *h*^2^ = 0.8 in height and between *h*^2^ = 0.4 and *h*^2^ = 0.6 in BMI [97, 98]. Taken together, these estimates suggest that non-additive effects may play a greater role in BMI than height, as we have observed here.

**Figure 3.**
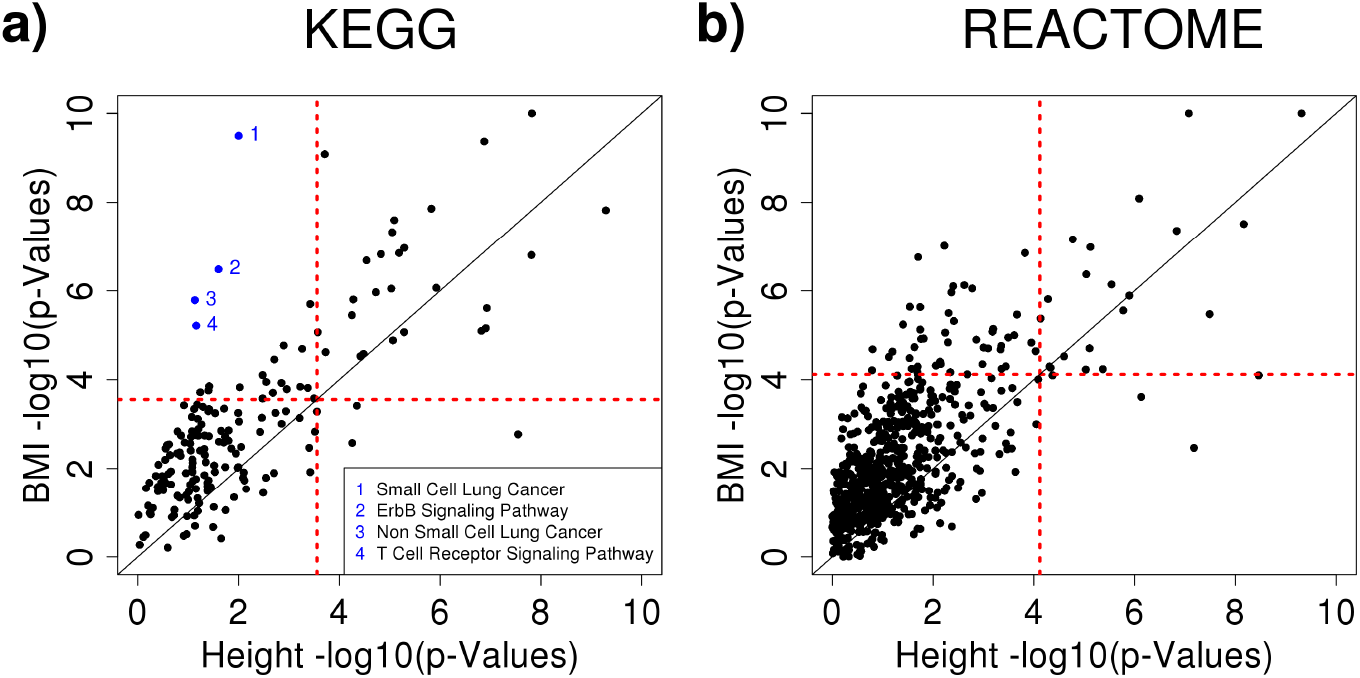
Scatterplots comparing the MAPIT-R *p*-values using (a) KEGG and (b) REACTOME pathways annotations in height and body mass index (BMI) within the African subgroup in the UK Biobank. For each plot, the *x*-axis shows the −log_10_ transformed MAPIT-R *p*-value for height, while the *y*-axis shows the same results for BMI. The red horizontal and vertical dashed lines are marked at the Bonferroni-corrected *p*-value thresholds for genome-wide significance in each pathway-phenotype combination (see Supplementary Table 1). Pathways in the top right quadrant have significant marginal epistatic effects in both traits; while, points in the bottom right and top left quadrants are pathways that are uniquely enriched in height or BMI, respectively. The four highlighted pathways in blue represent a cluster of oncogenic and signaling pathways whose loci have been functionally connected to BMI in previous studies [122–129]. Across both databases, BMI results have lower MAPIT-R *p*-values than height results on average. For these comparisons in all of the UK Biobank subgroups, see Supplementary Figure 21.

We detected one specific cluster of pathways in the KEGG database with notably divergent statistical evidence for marginal epistasis in height versus BMI (see Figure 3). These four highlighted pathways are related to oncogenic activity and include: genes associated with small cell lung cancer (*p*-value = 3.20 × 10^−10^), the ErbB signaling pathway (*p*-value = 3.30 × 10^−7^), genes associated with non-small cell lung cancer (*p*-value = 1.64 × 10^−6^), and T-cell receptor signaling (*p*-value = 6.12 × 10^−6^). There are predominantly two sets of gene families that appear in all four of these annotated gene sets: phosphatidylinositol 3-kinases (PI3Ks) and the AKT serine/threonine-protein kinases (see Supplementary Table 7). One particular gene in this group, *AKT2*, has been associated with multiple monogenic disorders of glucose metabolism, including severe insulin resistance and diabetes, and severe fasting hypoinsulinemic hypoglycemia [122–124], representing a possible driver of this cluster. Additionally, pharmacological inhibition of crosstalk between the PI3Ks has been shown to reduce adiposity and metabolic syndrome in both human beings and other model organisms [125–129].

### Testing Variability in MAPIT-R with British Replicate Subpopulations

One important consideration of our results is that the diverse non-European human ancestries in the UK Biobank have smaller sample sizes than recent GWA studies in individuals of European ancestry. Given the large sample size of over *N* = 470,000 individuals for the full white British cohort in the UK Biobank, we decided to test whether subsampled datasets from this group — similar in size to the non-European ancestry subgroups — would be large enough to gain insight into the genetic variation of height and BMI. Here, we sampled four additional, non-overlapping random subgroups of *N* = 4,000 British individuals and tested whether MAPIT-R results in these replicate subgroups were consistent with our results for the original British 4,000 subgroup. We also constructed larger non-overlapping British subsamples of *N* = 10,000 individuals to investigate how our results might vary with sample size. In total we analyzed five non-overlapping sets of *N* = 4,000 British individuals and five non-overlapping sets of *N* = 10,000 British individuals.

When applying MAPIT-R to these data replicates, we find that our results are robustly similar to what was observed in the original British 4,000 subgroup. Overall, there is a limited number of pathways with significant marginal epistatic effects, regardless of the pathway annotation scheme being used (i.e., KEGG versus REACTOME). Moreover, there is also limited overlap in the significant pathways that were detected between each of the subsampled replicates. These results are depicted and summarized in Supplementary Figures 7-12. As previously done with the individuals of non-European ancestry, we also checked that the null hypothesis of MAPIT-R remained well-calibrated on these subsampled British replicates by permuting the height and BMI measurements. Once again, we found that MAPIT-R continued to exhibit low empirical false discovery and type 1 error rates (Supplementary Tables 8 and 9). Altogether, the consistency of these analyses compared to the results with the original 4,000 individual British subgroup demonstrate that sample size does not appear to be a driving factor in the detection of pathway-level marginal epistasis.

### The Proteasome is Enriched for Marginal Epistasis Signals

To better identify the genes and genomic regions that are driving pathway-level marginal epistatic effects, we first investigated genes and gene families that are enriched amongst the significant pathways identified by MAPIT-R. To accomplish this, we conducted two types of hypergeometric tests for enrichment to detect genes that are overrepresented amongst the pathway annotations with low *p*-values (Supplementary Tables 3). In the first test, we took the annotations from a given database (i.e., KEGG or REACTOME) and implemented a standard hypergeometric test where we compared the number of times a gene appears within the set of significant epistatic pathways versus the number of times that same gene appears across all pathways in the database. This type of test, however, may be confounded by the fact that larger pathways naturally have more SNPs and are therefore more likely to be involved in non-additive genetic interactions (see Supplementary Figures 13 and 14). To mitigate this concern, we ran a second hypergeometric enrichment test using only pathways containing 1000 SNPs or fewer. By focusing on smaller pathways, we are better able to identify genes enriched for marginal epistasis versus spurious signals that may happen by chance in larger pathways.

Figure 4 shows the hypergeometric *p*-values for all genes in significant interacting pathways. Here, we focus on results for BMI within the African subgroup using annotations from the REACTOME database and we specifically highlight the only genes that were significant under both types of hypergeometric enrichment tests (i.e., the genes that were robustly identified as drivers regardless of the number of SNPs included in the test). Notably, these gene families (*PSMA, PSMB, PSMC, PSMD, PSME*, and *PSMF*) are all components of, or related to, the proteasome. The proteasome is a complex protein structure that acts as the catalytic half of the ubiquitin-proteasome system (UPS) — a critical system for the proper degradation of proteins within the cell [130–132]. The main proteasome isoform, 26S, is made up of two substructures: (i) the 20S core particle (CP) of four stacked rings (two outer structural rings encoded by *PSMA* genes and two inner catalytic rings encoded by *PSMB* genes), and (ii) the 19S regulatory particle (RP) which caps both ends of the CP (encoded by genes within both the *PSMC* and *PSMD* families). See Figure 5(a) for an illustration of this structure. Since these gene families covered both a large number of genomic sites, as well as biological functions known to be relevant to BMI, we used the proteasome as a test case to further refine the pathway-level signals identified by MAPIT-R.

**Figure 4.**
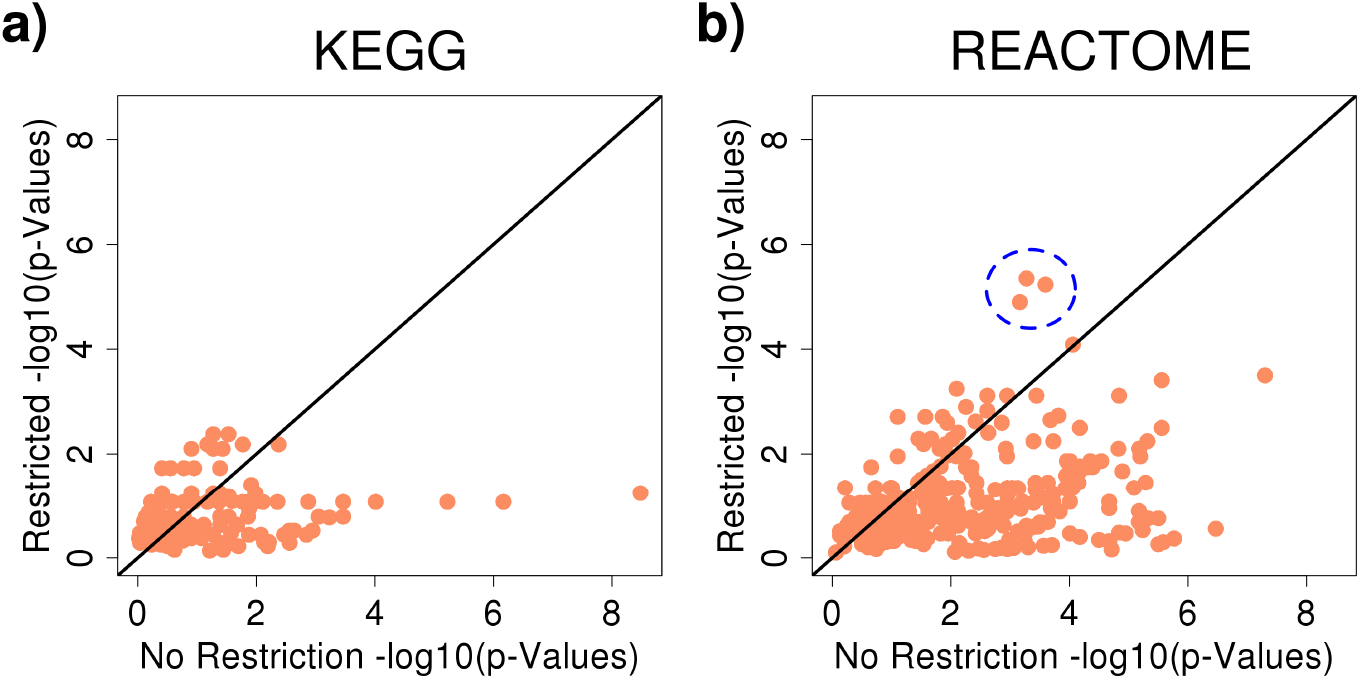
Scatterplots comparing the *p*-values from the hypergeometric enrichment analyses using only (a) KEGG and (b) REACTOME pathways annotations with at most 1000 SNPs within the African subgroup in the UK Biobank. Here, the gene-based *p*-values using the size restricted pathways are shown on the *y*-axis, while the results from the original unrestricted version of the analysis are shown on the *x*-axis. The blue dashed circle in panel **(b)** highlights the proteasome gene family cluster. For lists containing each gene’s original and size-restricted hypergeometric *p*-values, see Supplementary Table 11. Note that we only show results for BMI because few MAPIT-R significant pathways in the height analysis remained after imposing the size restriction. For lists containing gene counts for each database-phenotype-subgroup combination under both the original and size-restricted data sets, see Supplementary Tables 12 and 13.

**Figure 5.**
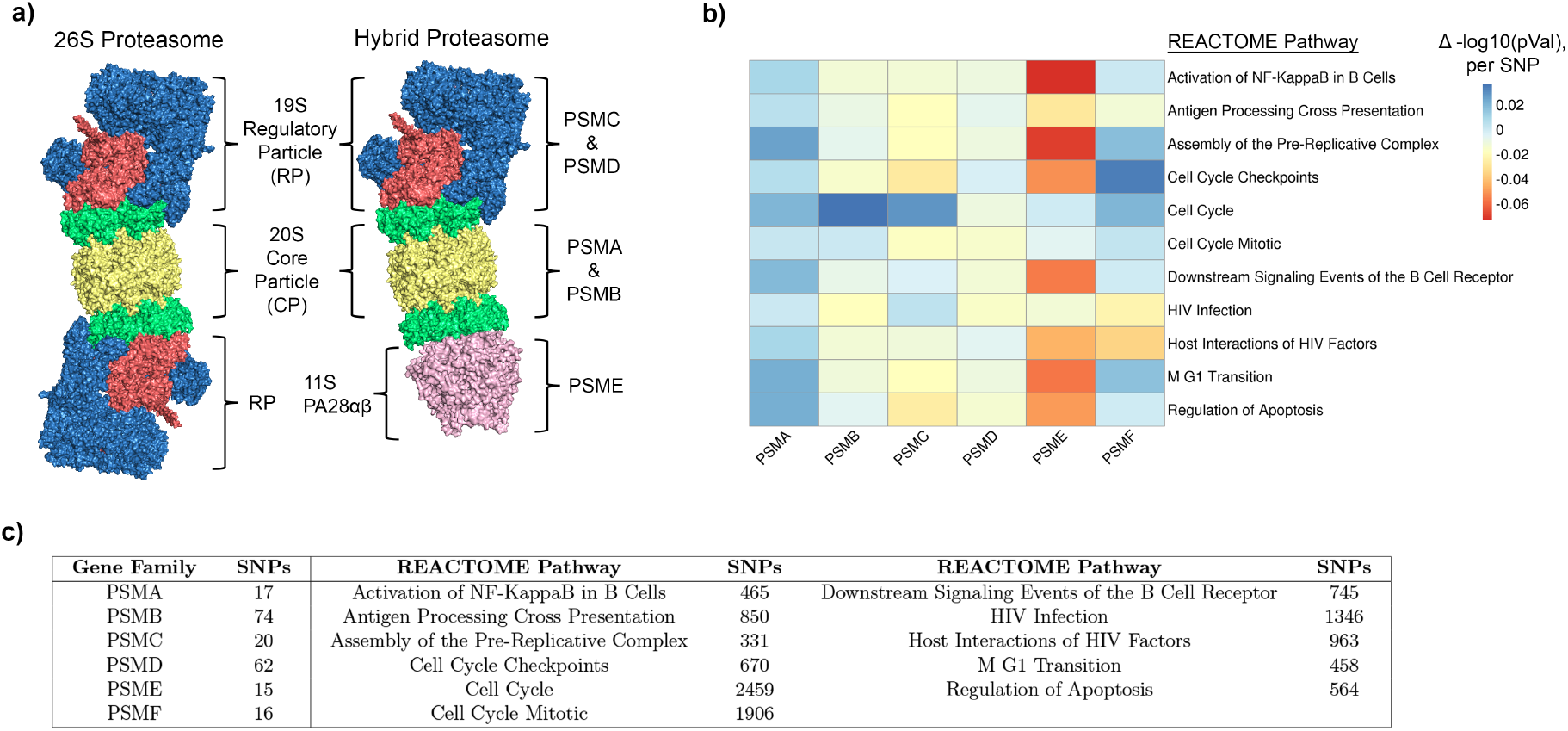
Structure of the proteasome and results from applying a “leave-one-out” approach to MAPIT-R with proteasome gene families. **(a)** Models of different isoforms of the proteasome, a complex protein structure required for proper degradation of many proteins in the cell. The “26S Proteasome” is the main isoform, composed of the 20S core particle (CP) and capped on both ends by the 19S regulatory particle (RP). The “Hybrid Proteasome” isoform is produced when the CP binds on one end with an RP and on the other end with the IFN-γ-inducible 11S complex PA28αβ. The *PSMA* and *PSMB* gene families encode components of the CP, the *PSMC* and *PSMD* gene families encode components of the RP, and members of the *PSME* gene family encode PA28αβ. Note that *PSMF* represents a proteasome inhibitor and is not shown. The structures shown were adopted and modified from the Protein Data Bank (human 26S proteasome, https://www.rcsb.org/structure/5GJR; mouse PA28αβ, https://www.rcsb.org/structure/5MX5) [141]. **(b)** The heatmap shows the change in original MAPIT-R −log_10_ *p*-value for different REACTOME pathways when each proteasome gene family is removed one at a time in a “leave-one-out” manner. The analyses were conducted in BMI for the African subgroup of the UK Biobank. The *x*-axis shows each proteasome gene family and the *y*-axis lists each REACTOME pathway. Each column has been scaled by the number of SNPs present in the given gene family and, as a result, the heatmap specifically shows the −log_10_ *p*-value change (△ in legend) per SNP. **(c)** The table shows the number of SNPs present in each proteasome gene family (left), as well as the number of SNPs present in each REACTOME pathway (right).

To investigate whether components of the proteasome served as a driver of significant marginal epistatic effects, we conducted a “leave-one-out” analysis with each of the gene families in the proteasome. More specifically, we first used MAPIT-R to reanalyze BMI after leaving out SNPs annotated within genes belonging to the *PSMA, PSMB, PSMC, PSMD, PSME*, and *PSMF* families, one family at a time. Next, we then compared these new “leave-one-out” MAPIT-R *p*-values to each pathway’s original *p*-value from running MAPIT-R on the full data. This enabled us to identify whether the removal of a particular gene family would lead to a notable loss of information regarding a pathway’s epistatic interactions with the rest of the genome.

Figure 5(b) shows the results from this analysis. We find that the *PSMA* and *PSME* gene families exhibit biologically interpretable changes in *p*-value magnitudes across multiple REACTOME pathways. For the *PSMA* gene family, we observe no examples where removing these genes leads to large increases in the MAPIT-R *p*-values. As previously mentioned, the *PSMA* gene family functionally encodes the outer two rings of the core four rings in the main 20S core. These outer “alpha” rings are gates which block entry into the core of the proteasome until they are opened by stimulation from the 19S regulatory particle [133–135]. And unlike the inner “beta” rings encoded by the *PSMB* family, which contain the proteolytic active sites, the outer rings do not have any catalytic functionality [136, 137]. This less direct role in the protein degradation process may explain the lack of increase in MAPIT-R *p*-values, or lack of information lost, when *PSMA* genes are removed from analysis.

For the *PSME* gene family, on the other hand, we find some of the largest increases in MAPIT-R *p*-values across multiple REACTOME pathways. Contextually, members of the *PSME* gene family encode an alternative regulatory particle, 11S PA28αβ, that also associates with the 20S core. PA28αβ is an Interferon-γ (IFN-γ) inducible regulatory protein that operates in a ubiquitin-independent manner and increases production of a particular subset of proteasomes known as immunoproteasomes [138–141]. Immunoproteasomes are specialized isoforms that are expressed at higher levels in hematopoietic cells and are more directly associated with immunity-related processes such as MHC antigen presentation [142–144]. Additionally, recent work has connected *PSME* genes to the regulation of NF-κB signaling [145, 146]. Altogether, these connections to immune activity may explain why removal of the *PSME* gene family affects marginal epistatic signals in pathways related to NF-κB, B-cells, HIV, and apoptosis. Lastly, conducting these “leave-one-out” MAPIT-R analyses in the other remaining UK Biobank subgroups, we observe that removing the *PSME* gene family also leads to some of the largest increases in MAPIT-R *p*-values in individuals of non-African ancestry as well (Supplementary Figures 15-20 and Supplementary Table 10). The consistency of this result across all subgroups suggests that *PSME* is a key contributor to proteasome epistatic interactions with other regions in the genome.

## Discussion

Here, we present the first scans for marginal epistasis within multiple human ancestries. We implement a new method, MAPIT-R, to test for evidence of non-additive genetic effects on the pathway-level and apply the framework to six different human ancestries sampled in the UK Biobank: African, British, Caribbean, Chinese, Indian, and Pakistani subgroups. Using two different pathway databases, we study continuous measurements of height and body mass index (BMI) and find a total of 245 pathways that have significant epistatic interactions with their polygenic background (see Figure 1). We find that the African subgroup produces the majority of these results, with over 65% of our 245 significant pathways being identified within this subgroup (see Figure 2). Additionally, we find that pathways related to immunity, cellular signaling, and metabolism have significant signals in our genome-wide marginal epistasis scans, and that BMI produces more significant marginal epistatic interactions at the pathway level than height (see Figure 3 and Table 1). In testing for drivers of our MAPIT-R results, we find evidence that the proteasome may be enriched for marginal epistatic interactions and characterize how proteasome gene families contribute to non-additive genetic architecture of complex traits (see Figures 4-5).

The fact that we find such an abundance of epistatic signals in the African subgroup underscores that African populations, and non-European ancestries in general, are particularly useful for complex trait genetics [66–68, 72, 147–152]. Past research has shown that African ancestry genomes offer a more complete characterization of the the genetic architecture of skin pigmentation [63, 64], reveal the evolutionary histories of *FOXP2* and other loci [153, 154], and are needed for more transferable polygenic risk scores [65, 70]. While many studies have generated a call for more GWA studies to be conducted in individuals of non-European ancestry [71, 73–75, 155], we believe this study reveals that our understanding of the role of epistasis in human complex trait architecture and broad-sense heritability will also expand with multiethnic analyses. Our results suggest that non-European ancestries, and African ancestries in particular, may be better suited for identifying signals of epistasis than European ancestries.

Our analyses are not without limitations. First, we are limited due to the computational costs of epistasis detection, although testing for marginal epistasis reduces our testing burden compared to standard exhaustive epistasis scans. Still, the MAPIT-R framework does not scale well to the full sample sizes of modern human genomic biobanks [43, 45, 84]. MAPIT-R encounters burdensome scalability when analyzing tens of thousands of individuals. One important future direction for research is to detect epistatic interactions using GWA summary statistics. Moving away from the need to have individual-level genotype-phenotype data to GWA study summary statistics has proven useful in both speeding up algorithmic efficiency as well as increasing power in multiple other GWA contexts [61, 156–160]. Another noticeable limitation is that MAPIT-R cannot be used to directly identify the interacting variant pairs that drive individual non-additive associations with a given trait. In particular, after identifying a pathway is involved in epistasis, it is still unclear which particular region of the genome it interacts with.

While the novel “leave-one-out” approach we implement here (see Figure 5(b)) helps narrow down the list of potential regions, MAPIT-R still does not directly identify pairs of interacting variants. Exploring marginal epistasis results *a posteriori* in a two step procedure can be one way to overcome these issues. For example, linking MAPIT-R with a framework that explicitly follows up on marginal epistasis signals with locus-focused methods such as fine-mapping [161–163] or co-localization [164–168] could further expand the power of the framework.

## URLs

MArginal ePIstasis Test for Regions (MAPIT-R) software, https://cran.r-project.org/web/packages/MAPITR/index.html; UK Biobank, https://www.ukbiobank.ac.uk; Molecular Signatures Database (MSigDB), https://www.gsea-msigdb.org/gsea/msigdb/index.jsp; Database of Genotypes and Phenotypes (db-GaP), https://www.ncbi.nlm.nih.gov/gap; NHGRI-EBI GWAS Catalog, https://www.ebi.ac.uk/gwas/; UCSC Genome Browser, https://genome.ucsc.edu/index.html; MArginal ePIstasis Test (MAPIT), https://github.com/lorinanthony/MAPIT; PLINK, https://www.cog-genomics.org/plink/1.9/.

## Acknowledgments

We thank Shigeo Murata for helpful feedback during the preparation of this manuscript. This research was conducted in part using computational resources and services at the Center for Computation and Visualization at Brown University. This research was also conducted using data from the UK Biobank Resource under Application Number 22419. G. Darnell was supported by NSF Grant No. DMS-1439786 while in residence at the Institute for Computational and Experimental Research in Mathematics (ICERM) in Providence, RI. This research was supported in part by grants P20GM109035 (COBRE Center for Computational Biology of Human Disease; PI Rand) and P20GM103645 (COBRE Center for Central Nervous; PI Sanes) from the NIH NIGMS, 2U10CA180794-06 from the NIH NCI and the Dana Farber Cancer Institute (PIs Gray and Gatsonis), as well as by an Alfred P. Sloan Research Fellowship (No. FG-2019-11622) awarded to L. Crawford. This research was also partly supported by the US National Institutes of Health (NIH) grant R01 GM118652, and the National Science Foundation (NSF) CAREER award DBI-1452622 to S. Ramachandran. Any opinions, findings, and conclusions or recommendations expressed in this material are those of the author(s) and do not necessarily reflect the views of any of the funders.

## Author Contributions

MCT, LC, and SR conceived the study design. LC and SR conceived the methods. MCT developed the software and carried out the analyses of the UK Biobank data. GD carried out the simulation studies. All authors wrote and reviewed the manuscript.

## References

1. Hirschhorn JN, Daly MJ. Genome-wide association studies for common diseases and complex traits [Journal Article]. Nat Rev Genet. 2005;6(2):95–108. Available from: http://www.ncbi.nlm.nih.gov/entrez/query.fcgi?cmd=Retrieve&db=PubMed&dopt=Citation&list_uids=15716906.

2. McCarthy MI, Abecasis GR, Cardon LR, Goldstein DB, Little J, Ioannidis JP, et al. Genome-wide association studies for complex traits: consensus, uncertainty and challenges [Journal Article]. Nat Rev Genet. 2008;9(5):356–69. Available from: http://www.ncbi.nlm.nih.gov/entrez/query.fcgi?cmd=Retrieve&db=PubMed&dopt=Citation&list_uids=18398418.

3. Stranger BE, Stahl EA, Raj T. Progress and promise of genome-wide association studies for human complex trait genetics [Journal Article]. Genetics. 2011;187(2):367–83. Available from: https://www.ncbi.nlm.nih.gov/pubmed/21115973.

4. Yang J, Zaitlen NA, Goddard ME, Visscher PM, Price AL. Advantages and pitfalls in the application of mixed-model association methods [Journal Article]. Nat Genet. 2014;46(2):100–6. Available from: https://www.ncbi.nlm.nih.gov/pubmed/24473328.

5. Pasaniuc B, Price AL. Dissecting the genetics of complex traits using summary association statistics [Journal Article]. Nat Rev Genet. 2017;18(2):117–127. Available from: https://www.ncbi.nlm.nih.gov/pubmed/27840428.

6. Visscher PM, Wray NR, Zhang Q, Sklar P, McCarthy MI, Brown MA, et al. 10 Years of GWAS Discovery: Biology, Function, and Translation [Journal Article]. Am J Hum Genet. 2017;101(1):5–22. Available from: https://www.ncbi.nlm.nih.gov/pubmed/28686856.

7. Buniello A, MacArthur JAL, Cerezo M, Harris LW, Hayhurst J, Malangone C, et al. The NHGRI-EBI GWAS Catalog of published genome-wide association studies, targeted arrays and summary statistics 2019 [Journal Article]. Nucleic Acids Res. 2019;47(D1):D1005–D1012. Available from: https://www.ncbi.nlm.nih.gov/pubmed/30445434.

8. Tam V, Patel N, Turcotte M, Bosse Y, Pare G, Meyre D. Benefits and limitations of genome-wide association studies [Journal Article]. Nat Rev Genet. 2019;20(8):467–484. Available from: https://www.ncbi.nlm.nih.gov/pubmed/31068683.

9. Lehner B, Crombie C, Tischler J, Fortunato A, Fraser AG. Systematic mapping of genetic interactions in Caenorhabditis elegans identifies common modifiers of diverse signaling pathways [Journal Article]. Nat Genet. 2006;38(8):896–903. Available from: https://www.ncbi.nlm.nih.gov/pubmed/16845399.

10. Rowe HC, Hansen BG, Halkier BA, Kliebenstein DJ. Biochemical networks and epistasis shape the Arabidopsis thaliana metabolome [Journal Article]. Plant Cell. 2008;20(5):1199–216. Available from: https://www.ncbi.nlm.nih.gov/pubmed/18515501.

11. Shao H, Burrage LC, Sinasac DS, Hill AE, Ernest SR, O’Brien W, et al. Genetic architecture of complex traits: large phenotypic effects and pervasive epistasis [Journal Article]. Proc Natl Acad Sci U S A. 2008;105(50):19910–4. Available from: https://www.ncbi.nlm.nih.gov/pubmed/19066216.

12. Flint J, Mackay TF. Genetic architecture of quantitative traits in mice, flies, and humans [Journal Article]. Genome Res. 2009;19(5):723–33. Available from: https://www.ncbi.nlm.nih.gov/pubmed/19411597.

13. Costanzo M, Baryshnikova A, Bellay J, Kim Y, Spear ED, Sevier CS, et al. The genetic landscape of a cell [Journal Article]. Science. 2010;327(5964):425–31. Available from: https://www.ncbi.nlm.nih.gov/pubmed/20093466.

14. He X, Qian W, Wang Z, Li Y, Zhang J. Prevalent positive epistasis in Escherichia coli and Saccharomyces cerevisiae metabolic networks [Journal Article]. Nat Genet. 2010;42(3):272–6. Available from: https://www.ncbi.nlm.nih.gov/pubmed/20101242.

15. Jarvis JP, Cheverud JM. Mapping the epistatic network underlying murine reproductive fatpad variation [Journal Article]. Genetics. 2011;187(2):597–610. Available from: https://www.ncbi.nlm.nih.gov/pubmed/21115969.

16. Pettersson M, Besnier F, Siegel PB, Carlborg O. Replication and explorations of high-order epistasis using a large advanced intercross line pedigree [Journal Article]. PLoS Genet. 2011;7(7):e1002180. Available from: https://www.ncbi.nlm.nih.gov/pubmed/21814519.

17. Bloom JS, Ehrenreich IM, Loo WT, Lite TL, Kruglyak L. Finding the sources of missing heritability in a yeast cross [Journal Article]. Nature. 2013;494(7436):234–7. Available from: https://www.ncbi.nlm.nih.gov/pubmed/23376951.

18. Monnahan PJ, Kelly JK. Epistasis Is a Major Determinant of the Additive Genetic Variance in Mimulus guttatus [Journal Article]. PLoS Genet. 2015;11(5):e1005201. Available from: https://www.ncbi.nlm.nih.gov/pubmed/25946702.

19. Carlborg O, Haley CS. Epistasis: too often neglected in complex trait studies? [Journal Article]. Nat Rev Genet. 2004;5(8):618–25. Available from: https://www.ncbi.nlm.nih.gov/pubmed/15266344.

20. Carlborg O, Jacobsson L, Ahgren P, Siegel P, Andersson L. Epistasis and the release of genetic variation during long-term selection [Journal Article]. Nat Genet. 2006;38(4):418–20. Available from: https://www.ncbi.nlm.nih.gov/pubmed/16532011.

21. Martin G, Elena SF, Lenormand T. Distributions of epistasis in microbes fit predictions from a fitness landscape model [Journal Article]. Nat Genet. 2007;39(4):555–60. Available from: https://www.ncbi.nlm.nih.gov/pubmed/17369829.

22. Phillips PC. Epistasis–the essential role of gene interactions in the structure and evolution of genetic systems [Journal Article]. Nat Rev Genet. 2008;9(11):855–67. Available from: https://www.ncbi.nlm.nih.gov/pubmed/18852697.

23. Moore JH, Williams SM. Epistasis and its implications for personal genetics [Journal Article]. Am J Hum Genet. 2009;85(3):309–20. Available from: https://www.ncbi.nlm.nih.gov/pubmed/19733727.

24. Zuk O, Hechter E, Sunyaev SR, Lander ES. The mystery of missing heritability: Genetic interactions create phantom heritability [Journal Article]. Proc Natl Acad Sci U S A. 2012;109(4):1193–8. Available from: https://www.ncbi.nlm.nih.gov/pubmed/22223662.

25. Jones AG, Burger R, Arnold SJ. Epistasis and natural selection shape the mutational architecture of complex traits [Journal Article]. Nat Commun. 2014;5:3709. Available from: https://www.ncbi.nlm.nih.gov/pubmed/24828461.

26. Mackay TF. Epistasis and quantitative traits: using model organisms to study gene-gene interactions [Journal Article]. Nat Rev Genet. 2014;15(1):22–33. Available from: https://www.ncbi.nlm.nih.gov/pubmed/24296533.

27. Hill WG, Goddard ME, Visscher PM. Data and theory point to mainly additive genetic variance for complex traits [Journal Article]. PLoS Genet. 2008;4(2):e1000008. Available from: https://www.ncbi.nlm.nih.gov/pubmed/18454194.

28. Crow JF. On epistasis: why it is unimportant in polygenic directional selection [Journal Article]. Philos Trans R Soc Lond B Biol Sci. 2010;365(1544):1241–4. Available from: https://www.ncbi.nlm.nih.gov/pubmed/20308099.

29. Yang J, Benyamin B, McEvoy BP, Gordon S, Henders AK, Nyholt DR, et al. Common SNPs explain a large proportion of the heritability for human height [Journal Article]. Nat Genet. 2010;42(7):565–9. Available from: https://www.ncbi.nlm.nih.gov/pubmed/20562875.

30. Aschard H, Chen J, Cornelis MC, Chibnik LB, Karlson EW, Kraft P. Inclusion of gene-gene and gene-environment interactions unlikely to dramatically improve risk prediction for complex diseases [Journal Article]. Am J Hum Genet. 2012;90(6):962–72. Available from: https://www.ncbi.nlm.nih.gov/pubmed/22633398.

31. Powell JE, Henders AK, McRae AF, Kim J, Hemani G, Martin NG, et al. Congruence of additive and non-additive effects on gene expression estimated from pedigree and SNP data [Journal Article]. PLoS Genet. 2013;9(5):e1003502. Available from: https://www.ncbi.nlm.nih.gov/pubmed/23696747.

32. Maki-Tanila A, Hill WG. Influence of gene interaction on complex trait variation with multilocus models [Journal Article]. Genetics. 2014;198(1):355–67. Available from: https://www.ncbi.nlm.nih.gov/pubmed/24990992.

33. Wood AR, Esko T, Yang J, Vedantam S, Pers TH, Gustafsson S, et al. Defining the role of common variation in the genomic and biological architecture of adult human height [Journal Article]. Nat Genet. 2014a;46(11):1173–86. Available from: https://www.ncbi.nlm.nih.gov/pubmed/25282103.

34. Yang J, Bakshi A, Zhu Z, Hemani G, Vinkhuyzen AA, Lee SH, et al. Genetic variance estimation with imputed variants finds negligible missing heritability for human height and body mass index [Journal Article]. Nat Genet. 2015;47(10):1114–20. Available from: https://www.ncbi.nlm.nih.gov/pubmed/26323059.

35. Huang W, Mackay TF. The Genetic Architecture of Quantitative Traits Cannot Be Inferred from Variance Component Analysis [Journal Article]. PLoS Genet. 2016;12(11):e1006421. Available from: https://www.ncbi.nlm.nih.gov/pubmed/27812106.

36. Maher B. Personal genomes: The case of the missing heritability [Journal Article]. Nature. 2008;456(7218):18–21. Available from: https://www.ncbi.nlm.nih.gov/pubmed/18987709.

37. Manolio TA, Collins FS, Cox NJ, Goldstein DB, Hindorff LA, Hunter DJ, et al. Finding the missing heritability of complex diseases [Journal Article]. Nature. 2009;461(7265):747–53. Available from: https://www.ncbi.nlm.nih.gov/pubmed/19812666.

38. Eichler EE, Flint J, Gibson G, Kong A, Leal SM, Moore JH, et al. Missing heritability and strategies for finding the underlying causes of complex disease [Journal Article]. Nat Rev Genet. 2010;11(6):446–50. Available from: https://www.ncbi.nlm.nih.gov/pubmed/20479774.

39. Slatkin M. Epigenetic inheritance and the missing heritability problem [Journal Article]. Genetics. 2009;182(3):845–50. Available from: https://www.ncbi.nlm.nih.gov/pubmed/19416939.

40. Hemani G, Knott S, Haley C. An evolutionary perspective on epistasis and the missing heritability [Journal Article]. PLoS Genet. 2013;9(2):e1003295. Available from: https://www.ncbi.nlm.nih.gov/pubmed/23509438.

41. Wainschtein P, Jain DP, Yengo L, Zheng Z, Cupples LA, Shadyab AH, et al. Recovery of trait heritability from whole genome sequence data [Journal Article]. bioRxiv. 2019;p. 588020. Available from: https://www.biorxiv.org/content/biorxiv/early/2019/03/25/588020.full.pdf.

42. Roberts GHL, Santorico SA, Spritz RA. Deep genotype imputation captures virtually all heritability of autoimmune vitiligo [Journal Article]. Hum Mol Genet. 2020;29(5):859–863. Available from: https://www.ncbi.nlm.nih.gov/pubmed/31943001.

43. Crawford L, Zeng P, Mukherjee S, Zhou X. Detecting epistasis with the marginal epistasis test in genetic mapping studies of quantitative traits [Journal Article]. PLoS Genet. 2017a;13(7):e1006869. Available from: https://www.ncbi.nlm.nih.gov/pubmed/28746338.

44. Crawford L, Zhou X. Genome-wide Marginal Epistatic Association Mapping in Case-Control Studies [Journal Article]. bioRxiv. 2018b;p. 374983. Available from: https://www.biorxiv.org/content/biorxiv/early/2018/07/23/374983.full.pdf.

45. Moore R, Casale FP, Jan Bonder M, Horta D, Consortium B, Franke L, et al. A linear mixed-model approach to study multivariate gene-environment interactions [Journal Article]. Nat Genet. 2019;51(1):180–186. Available from: https://www.ncbi.nlm.nih.gov/pubmed/30478441.

46. Wang H, Yue T, Yang J, Wu W, Xing EP. Deep mixed model for marginal epistasis detection and population stratification correction in genome-wide association studies [Journal Article]. BMC Bioinformatics. 2019;20(Suppl 23):656. Available from: https://www.ncbi.nlm.nih.gov/pubmed/31881907.

47. Zhou X, Carbonetto P, Stephens M. Polygenic modeling with bayesian sparse linear mixed models [Journal Article]. PLoS Genet. 2013;9(2):e1003264. Available from: https://www.ncbi.nlm.nih.gov/pubmed/23408905.

48. Bulik-Sullivan BK, Loh PR, Finucane HK, Ripke S, Yang J, Schizophrenia Working Group of the Psychiatric Genomics C, et al. LD Score regression distinguishes confounding from polygenicity in genome-wide association studies [Journal Article]. Nat Genet. 2015;47(3):291–5. Available from: https://www.ncbi.nlm.nih.gov/pubmed/25642630.

49. Wray NR, Wijmenga C, Sullivan PF, Yang J, Visscher PM. Common Disease Is More Complex Than Implied by the Core Gene Omnigenic Model [Journal Article]. Cell. 2018;173(7):1573–1580. Available from: https://www.ncbi.nlm.nih.gov/pubmed/29906445.

50. Subramanian A, Tamayo P, Mootha VK, Mukherjee S, Ebert BL, Gillette MA, et al. Gene set enrichment analysis: a knowledge-based approach for interpreting genome-wide expression profiles [Journal Article]. Proc Natl Acad Sci U S A. 2005;102(43):15545–50. Available from: https://www.ncbi.nlm.nih.gov/pubmed/16199517.

51. Cantor RM, Lange K, Sinsheimer JS. Prioritizing GWAS results: A review of statistical methods and recommendations for their application [Journal Article]. Am J Hum Genet. 2010;86(1):6–22. Available from: https://www.ncbi.nlm.nih.gov/pubmed/20074509.

52. Wang K, Li M, Hakonarson H. Analysing biological pathways in genome-wide association studies [Journal Article]. Nat Rev Genet. 2010b;11(12):843–54. Available from: https://www.ncbi.nlm.nih.gov/pubmed/21085203.

53. Lee S, Emond MJ, Bamshad MJ, Barnes KC, Rieder MJ, Nickerson DA, et al. Optimal unified approach for rare-variant association testing with application to small-sample case-control whole-exome sequencing studies [Journal Article]. Am J Hum Genet. 2012;91(2):224–37. Available from: http://www.ncbi.nlm.nih.gov/pubmed/22863193.

54. Carbonetto P, Stephens M. Integrated enrichment analysis of variants and pathways in genome-wide association studies indicates central role for IL-2 signaling genes in type 1 diabetes, and cytokine signaling genes in Crohn’s disease [Journal Article]. PLoS Genet. 2013;9(10):e1003770. Available from: https://www.ncbi.nlm.nih.gov/pubmed/24098138.

55. Mooney MA, Nigg JT, McWeeney SK, Wilmot B. Functional and genomic context in pathway analysis of GWAS data [Journal Article]. Trends Genet. 2014;30(9):390–400. Available from: https://www.ncbi.nlm.nih.gov/pubmed/25154796.

56. Gamazon ER, Wheeler HE, Shah KP, Mozaffari SV, Aquino-Michaels K, Carroll RJ, et al. A gene-based association method for mapping traits using reference transcriptome data [Journal Article]. Nat Genet. 2015;47(9):1091–8. Available from: https://www.ncbi.nlm.nih.gov/pubmed/26258848.

57. de Leeuw CA, Neale BM, Heskes T, Posthuma D. The statistical properties of gene-set analysis [Journal Article]. Nat Rev Genet. 2016;17(6):353–64. Available from: https://www.ncbi.nlm.nih.gov/pubmed/27070863.

58. Nakka P, Raphael BJ, Ramachandran S. Gene and Network Analysis of Common Variants Reveals Novel Associations in Multiple Complex Diseases [Journal Article]. Genetics. 2016;204(2):783–798. Available from: https://www.ncbi.nlm.nih.gov/pubmed/27489002.

59. Zhu X, Stephens M. Large-scale genome-wide enrichment analyses identify new trait-associated genes and pathways across 31 human phenotypes [Journal Article]. Nat Commun. 2018;9(1):4361. Available from: https://www.ncbi.nlm.nih.gov/pubmed/30341297.

60. Sun R, Hui S, Bader GD, Lin X, Kraft P. Powerful gene set analysis in GWAS with the Generalized Berk-Jones statistic [Journal Article]. PLoS Genet. 2019;15(3):e1007530. Available from: https://www.ncbi.nlm.nih.gov/pubmed/30875371.

61. Cheng W, Ramachandran S, Crawford L. Estimation of non-null SNP effect size distributions enables the detection of enriched genes underlying complex traits [Journal Article]. PLoS Genet. 2020;16(6):e1008855. Available from: https://www.ncbi.nlm.nih.gov/pubmed/32542026.

62. Nakka P, Archer NP, Xu H, Lupo PJ, Raphael BJ, Yang JJ, et al. Novel Gene and Network Associations Found for Acute Lymphoblastic Leukemia Using Case-Control and Family-Based Studies in Multiethnic Populations [Journal Article]. Cancer Epidemiol Biomarkers Prev. 2017;26(10):1531–1539. Available from: https://www.ncbi.nlm.nih.gov/pubmed/28751478.

63. Martin AR, Lin M, Granka JM, Myrick JW, Liu X, Sockell A, et al. An Unexpectedly Complex Architecture for Skin Pigmentation in Africans [Journal Article]. Cell. 2017b;171(6):1340–1353 e14. Available from: https://www.ncbi.nlm.nih.gov/pubmed/29195075.

64. Crawford NG, Kelly DE, Hansen MEB, Beltrame MH, Fan S, Bowman SL, et al. Loci associated with skin pigmentation identified in African populations [Journal Article]. Science. 2017b;358(6365). Available from: https://www.ncbi.nlm.nih.gov/pubmed/29025994.

65. Duncan L, Shen H, Gelaye B, Meijsen J, Ressler K, Feldman M, et al. Analysis of polygenic risk score usage and performance in diverse human populations [Journal Article]. Nat Commun. 2019;10(1):3328. Available from: https://www.ncbi.nlm.nih.gov/pubmed/31346163.

66. Kuchenbaecker K, Telkar N, Reiker T, Walters RG, Lin K, Eriksson A, et al. The transferability of lipid loci across African, Asian and European cohorts [Journal Article]. Nat Commun. 2019;10(1):4330. Available from: https://www.ncbi.nlm.nih.gov/pubmed/31551420.

67. Zhong Y, Perera MA, Gamazon ER. On Using Local Ancestry to Characterize the Genetic Architecture of Human Traits: Genetic Regulation of Gene Expression in Multiethnic or Admixed Populations [Journal Article]. Am J Hum Genet. 2019;104(6):1097–1115. Available from: https://www.ncbi.nlm.nih.gov/pubmed/31104770.

68. Wojcik GL, Graff M, Nishimura KK, Tao R, Haessler J, Gignoux CR, et al. Genetic analyses of diverse populations improves discovery for complex traits [Journal Article]. Nature. 2019;570(7762):514–518. Available from: https://www.ncbi.nlm.nih.gov/pubmed/31217584.

69. Chen MH, Raffield LM, Mousas A, Sakaue S, Huffman JE, Moscati A, et al. Trans-ethnic and Ancestry-Specific Blood-Cell Genetics in 746,667 Individuals from 5 Global Populations [Journal Article]. Cell. 2020;182(5):1198–1213.e14. Available from: http://www.sciencedirect.com/science/article/pii/S0092867420308229.

70. Marnetto D, Parna K, Lall K, Molinaro L, Montinaro F, Haller T, et al. Ancestry deconvolution and partial polygenic score can improve susceptibility predictions in recently admixed individuals [Journal Article]. Nat Commun. 2020;11(1):1628. Available from: https://www.ncbi.nlm.nih.gov/pubmed/32242022.

71. Popejoy AB, Fullerton SM. Genomics is failing on diversity [Journal Article]. Nature. 2016;538(7624):161–164. Available from: https://www.ncbi.nlm.nih.gov/pubmed/27734877.

72. Martin AR, Teferra S, Moller M, Hoal EG, Daly MJ. The critical needs and challenges for genetic architecture studies in Africa [Journal Article]. Curr Opin Genet Dev. 2018;53:113–120. Available from: https://www.ncbi.nlm.nih.gov/pubmed/30240950.

73. Martin AR, Kanai M, Kamatani Y, Okada Y, Neale BM, Daly MJ. Clinical use of current polygenic risk scores may exacerbate health disparities [Journal Article]. Nat Genet. 2019;51(4):584–591. Available from: https://www.ncbi.nlm.nih.gov/pubmed/30926966.

74. Gurdasani D, Barroso I, Zeggini E, Sandhu MS. Genomics of disease risk in globally diverse populations [Journal Article]. Nat Rev Genet. 2019;20(9):520–535. Available from: https://www.ncbi.nlm.nih.gov/pubmed/31235872.

75. Sirugo G, Williams SM, Tishkoff SA. The Missing Diversity in Human Genetic Studies [Journal Article]. Cell. 2019;177(1):26–31. Available from: https://www.ncbi.nlm.nih.gov/pubmed/30901543.

76. Ma L, Brautbar A, Boerwinkle E, Sing CF, Clark AG, Keinan A. Knowledge-driven analysis identifies a gene-gene interaction affecting high-density lipoprotein cholesterol levels in multiethnic populations [Journal Article]. PLoS Genet. 2012;8(5):e1002714. Available from: https://www.ncbi.nlm.nih.gov/pubmed/22654671.

77. Fish AE, Capra JA, Bush WS. Are Interactions between cis-Regulatory Variants Evidence for Biological Epistasis or Statistical Artifacts? [Journal Article]. Am J Hum Genet. 2016;99(4):817–830. Available from: https://www.ncbi.nlm.nih.gov/pubmed/27640306.

78. Choquet H, Paylakhi S, Kneeland SC, Thai KK, Hoffmann TJ, Yin J, et al. A multiethnic genome-wide association study of primary open-angle glaucoma identifies novel risk loci [Journal Article]. Nat Commun. 2018;9(1):2278. Available from: https://www.ncbi.nlm.nih.gov/pubmed/29891935.

79. Hoffmann TJ, Choquet H, Yin J, Banda Y, Kvale MN, Glymour M, et al. A Large Multiethnic Genome-Wide Association Study of Adult Body Mass Index Identifies Novel Loci [Journal Article]. Genetics. 2018;210(2):499–515. Available from: https://www.ncbi.nlm.nih.gov/pubmed/30108127.

80. Liberzon A, Subramanian A, Pinchback R, Thorvaldsdottir H, Tamayo P, Mesirov JP. Molecular signatures database (MSigDB) 3.0 [Journal Article]. Bioinformatics. 2011;27(12):1739–40. Available from: https://www.ncbi.nlm.nih.gov/pubmed/21546393.

81. Sudlow C, Gallacher J, Allen N, Beral V, Burton P, Danesh J, et al. UK biobank: an open access resource for identifying the causes of a wide range of complex diseases of middle and old age [Journal Article]. PLoS Med. 2015;12(3):e1001779. Available from: https://www.ncbi.nlm.nih.gov/pubmed/25826379.

82. Jiang Y, Reif JC. Modeling Epistasis in Genomic Selection [Journal Article]. Genetics. 2015;201(2):759–68. Available from: https://www.ncbi.nlm.nih.gov/pubmed/26219298.

83. Zhou X. A Unified Framework for Variance Component Estimation with Summary Statistics in Genome-Wide Association Studies [Journal Article]. Ann Appl Stat. 2017;11(4):2027–2051. Available from: https://www.ncbi.nlm.nih.gov/pubmed/29515717.

84. Crawford L, Wood KC, Zhou X, Mukherjee S. Bayesian Approximate Kernel Regression with Variable Selection [Journal Article]. J Am Stat Assoc. 2018a;113(524):1710–1721. Available from: https://www.ncbi.nlm.nih.gov/pubmed/30799887.

85. Haseman JK, Elston RC. The investigation of linkage between a quantitative trait and a marker locus [Journal Article]. Behav Genet. 1972;2(1):3–19. Available from: https://www.ncbi.nlm.nih.gov/pubmed/4157472.

86. Liu JZ, McRae AF, Nyholt DR, Medland SE, Wray NR, Brown KM, et al. A versatile gene-based test for genome-wide association studies [Journal Article]. Am J Hum Genet. 2010;87(1):139–45. Available from: https://www.ncbi.nlm.nih.gov/pubmed/20598278.

87. Wu MC, Lee S, Cai T, Li Y, Boehnke M, Lin X. Rare-variant association testing for sequencing data with the sequence kernel association test [Journal Article]. Am J Hum Genet. 2011;89(1):82–93. Available from: http://www.ncbi.nlm.nih.gov/pubmed/21737059.

88. Ionita-Laza I, Lee S, Makarov V, Buxbaum JD, Lin X. Sequence kernel association tests for the combined effect of rare and common variants [Journal Article]. Am J Hum Genet. 2013;92(6):841–53. Available from: https://www.ncbi.nlm.nih.gov/pubmed/23684009.

89. Wang M, Huang J, Liu Y, Ma L, Potash JB, Han S. COMBAT: A Combined Association Test for Genes Using Summary Statistics [Journal Article]. Genetics. 2017;207(3):883–891. Available from: https://www.ncbi.nlm.nih.gov/pubmed/28878002.

90. Davies RB. Algorithm AS 155: The Distribution of a Linear Combination of χ^2^ Squared Random Variables [Journal Article]. Journal of the Royal Statistical Society Series C (Applied Statistics). 1980;29(3):323–333. Available from: http://www.jstor.org/stable/2346911.

91. Chen H, Meigs JB, Dupuis J. Sequence kernel association test for quantitative traits in family samples [Journal Article]. Genet Epidemiol. 2013;37(2):196–204. Available from: https://www.ncbi.nlm.nih.gov/pubmed/23280576.

92. Kuonen D. Saddlepoint Approximations for Distributions of Quadratic Forms in Normal Variables [Journal Article]. Biometrika. 1999;86(4):929–935. Available from: http://www.jstor.org.revproxy.brown.edu/stable/2673596.

93. Satterthwaite FE. An approximate distribution of estimates of variance components [Journal Article]. Biometrics. 1946;2(6):110–4. Available from: https://www.ncbi.nlm.nih.gov/pubmed/20287815.

94. Wang K, Li M, Hakonarson H. ANNOVAR: functional annotation of genetic variants from high-throughput sequencing data [Journal Article]. Nucleic Acids Res. 2010a;38(16):e164. Available from: http://www.ncbi.nlm.nih.gov/pubmed/20601685.

95. Das S, Forer L, Schonherr S, Sidore C, Locke AE, Kwong A, et al. Next-generation genotype imputation service and methods [Journal Article]. Nat Genet. 2016;48(10):1284–1287. Available from: https://www.ncbi.nlm.nih.gov/pubmed/27571263.

96. Locke AE, Kahali B, Berndt SI, Justice AE, Pers TH, Day FR, et al. Genetic studies of body mass index yield new insights for obesity biology [Journal Article]. Nature. 2015;518(7538):197–206. Available from: https://www.ncbi.nlm.nih.gov/pubmed/25673413.

97. Elks CE, den Hoed M, Zhao JH, Sharp SJ, Wareham NJ, Loos RJ, et al. Variability in the heritability of body mass index: a systematic review and meta-regression [Journal Article]. Front Endocrinol (Lausanne). 2012;3:29. Available from: https://www.ncbi.nlm.nih.gov/pubmed/22645519.

98. Visscher PM, Brown MA, McCarthy MI, Yang J. Five years of GWAS discovery [Journal Article]. Am J Hum Genet. 2012;90(1):7–24. Available from: https://www.ncbi.nlm.nih.gov/pubmed/22243964.

99. Finucane HK, Bulik-Sullivan B, Gusev A, Trynka G, Reshef Y, Loh PR, et al. Partitioning heritability by functional annotation using genome-wide association summary statistics [Journal Article]. Nat Genet. 2015;47(11):1228–35. Available from: https://www.ncbi.nlm.nih.gov/pubmed/26414678.

100. Speed D, Cai N, Consortium U, Johnson MR, Nejentsev S, Balding DJ. Reevaluation of SNP heritability in complex human traits [Journal Article]. Nat Genet. 2017;49(7):986–992. Available from: https://www.ncbi.nlm.nih.gov/pubmed/28530675.

101. Martin MP, Gao X, Lee JH, Nelson GW, Detels R, Goedert JJ, et al. Epistatic interaction between KIR3DS1 and HLA-B delays the progression to AIDS [Journal Article]. Nat Genet. 2002;31(4):429–34. Available from: https://www.ncbi.nlm.nih.gov/pubmed/12134147.

102. Williams TN, Mwangi TW, Wambua S, Peto TE, Weatherall DJ, Gupta S, et al. Negative epistasis between the malaria-protective effects of alpha+-thalassemia and the sickle cell trait [Journal Article]. Nat Genet. 2005;37(11):1253–7. Available from: https://www.ncbi.nlm.nih.gov/pubmed/16227994.

103. Wan X, Yang C, Yang Q, Xue H, Fan X, Tang NL, et al. BOOST: A fast approach to detecting gene-gene interactions in genome-wide case-control studies [Journal Article]. Am J Hum Genet. 2010;87(3):325–40. Available from: https://www.ncbi.nlm.nih.gov/pubmed/20817139.

104. Rose AM, Bell LC. Epistasis and immunity: the role of genetic interactions in autoimmune diseases [Journal Article]. Immunology. 2012;137(2):131–8. Available from: https://www.ncbi.nlm.nih.gov/pubmed/22804709.

105. Lareau CA, White BC, Oberg AL, Kennedy RB, Poland GA, McKinney BA. An interaction quantitative trait loci tool implicates epistatic functional variants in an apoptosis pathway in smallpox vaccine eQTL data [Journal Article]. Genes Immun. 2016;17(4):244–50. Available from: https://www.ncbi.nlm.nih.gov/pubmed/27052692.

106. Opi DH, Swann O, Macharia A, Uyoga S, Band G, Ndila CM, et al. Two complement receptor one alleles have opposing associations with cerebral malaria and interact with alpha(+)thalassaemia [Journal Article]. Elife. 2018;7. Available from: https://www.ncbi.nlm.nih.gov/pubmed/29690995.

107. Zhang J, Wei Z, Cardinale CJ, Gusareva ES, Van Steen K, Sleiman P, et al. Multiple Epistasis Interactions Within MHC Are Associated With Ulcerative Colitis [Journal Article]. Front Genet. 2019;10:257. Available from: https://www.ncbi.nlm.nih.gov/pubmed/31001315.

108. Segre D, Deluna A, Church GM, Kishony R. Modular epistasis in yeast metabolism [Journal Article]. Nat Genet. 2005;37(1):77–83. Available from: https://www.ncbi.nlm.nih.gov/pubmed/15592468.

109. Snitkin ES, Segre D. Epistatic interaction maps relative to multiple metabolic phenotypes [Journal Article]. PLoS Genet. 2011;7(2):e1001294. Available from: https://www.ncbi.nlm.nih.gov/pubmed/21347328.

110. Podgornaia AI, Laub MT. Protein evolution. Pervasive degeneracy and epistasis in a proteinprotein interface [Journal Article]. Science. 2015;347(6222):673–7. Available from: https://www.ncbi.nlm.nih.gov/pubmed/25657251.

111. Sorrells TR, Booth LN, Tuch BB, Johnson AD. Intersecting transcription networks constrain gene regulatory evolution [Journal Article]. Nature. 2015;523(7560):361–5. Available from: https://www.ncbi.nlm.nih.gov/pubmed/26153861.

112. Tyler AL, Ji B, Gatti DM, Munger SC, Churchill GA, Svenson KL, et al. Epistatic Networks Jointly Influence Phenotypes Related to Metabolic Disease and Gene Expression in Diversity Outbred Mice [Journal Article]. Genetics. 2017;206(2):621–639. Available from: https://www.ncbi.nlm.nih.gov/pubmed/28592500.

113. Nghe P, Kogenaru M, Tans SJ. Sign epistasis caused by hierarchy within signalling cascades [Journal Article]. Nat Commun. 2018;9(1):1451. Available from: https://www.ncbi.nlm.nih.gov/pubmed/29654280.

114. Jiao H, Zang Y, Zhang M, Zhang Y, Wang Y, Wang K, et al. Genome-Wide Interaction and Pathway Association Studies for Body Mass Index [Journal Article]. Front Genet. 2019;10:404. Available from: https://www.ncbi.nlm.nih.gov/pubmed/31118946.

115. Marouli E, Graff M, Medina-Gomez C, Lo KS, Wood AR, Kjaer TR, et al. Rare and low-frequency coding variants alter human adult height [Journal Article]. Nature. 2017;542(7640):186–190. Available from: https://www.ncbi.nlm.nih.gov/pubmed/28146470.

116. Wood AR, Tuke MA, Nalls MA, Hernandez DG, Bandinelli S, Singleton AB, et al. Another explanation for apparent epistasis [Journal Article]. Nature. 2014b;514(7520):E3–5. Available from: https://www.ncbi.nlm.nih.gov/pubmed/25279928.

117. Salinas YD, Wang L, DeWan AT. Multiethnic genome-wide association study identifies ethnicspecific associations with body mass index in Hispanics and African Americans [Journal Article]. BMC Genet. 2016;17(1):78. Available from: https://www.ncbi.nlm.nih.gov/pubmed/27296613.

118. Jha P, McDevitt MT, Halilbasic E, Williams EG, Quiros PM, Gariani K, et al. Genetic Regulation of Plasma Lipid Species and Their Association with Metabolic Phenotypes [Journal Article]. Cell Syst. 2018;6(6):709–721 e6. Available from: https://www.ncbi.nlm.nih.gov/pubmed/29909275.

119. Cousminer DL, Berry DJ, Timpson NJ, Ang W, Thiering E, Byrne EM, et al. Genome-wide association and longitudinal analyses reveal genetic loci linking pubertal height growth, pubertal timing and childhood adiposity [Journal Article]. Hum Mol Genet. 2013;22(13):2735–47. Available from: https://www.ncbi.nlm.nih.gov/pubmed/23449627.

120. Stahl EA, Wegmann D, Trynka G, Gutierrez-Achury J, Do R, Voight BF, et al. Bayesian inference analyses of the polygenic architecture of rheumatoid arthritis [Journal Article]. Nat Genet. 2012;44(5):483–9. Available from: https://www.ncbi.nlm.nih.gov/pubmed/22446960.

121. Deng Z, Zhen J, Harrison GF, Zhang G, Chen R, Sun G, et al. Genetically Determined Strength of Natural Killer Cells is Enhanced by Adaptive HLA class I Admixture in East Asians [Journal Article]. bioRxiv. 2020;p. 2020.07.29.227579. Available from: https://www.biorxiv.org/content/biorxiv/early/2020/07/30/2020.07.29.227579.full.pdf.

122. George S, Rochford JJ, Wolfrum C, Gray SL, Schinner S, Wilson JC, et al. A family with severe insulin resistance and diabetes due to a mutation in AKT2 [Journal Article]. Science. 2004;304(5675):1325–8. Available from: https://www.ncbi.nlm.nih.gov/pubmed/15166380.

123. Manning A, Highland HM, Gasser J, Sim X, Tukiainen T, Fontanillas P, et al. A Low-Frequency Inactivating AKT2 Variant Enriched in the Finnish Population Is Associated With Fasting Insulin Levels and Type 2 Diabetes Risk [Journal Article]. Diabetes. 2017;66(7):2019–2032. Available from: https://www.ncbi.nlm.nih.gov/pubmed/28341696.

124. Latva-Rasku A, Honka MJ, Stancakova A, Koistinen HA, Kuusisto J, Guan L, et al. A Partial Loss-of-Function Variant in AKT2 Is Associated With Reduced Insulin-Mediated Glucose Uptake in Multiple Insulin-Sensitive Tissues: A Genotype-Based Callback Positron Emission Tomography Study [Journal Article]. Diabetes. 2018;67(2):334–342. Available from: https://www.ncbi.nlm.nih.gov/pubmed/29141982.

125. Ortega-Molina A, Lopez-Guadamillas E, Mattison JA, Mitchell SJ, Munoz-Martin M, Iglesias G, et al. Pharmacological inhibition of PI3K reduces adiposity and metabolic syndrome in obese mice and rhesus monkeys [Journal Article]. Cell Metab. 2015;21(4):558–70. Available from: https://www.ncbi.nlm.nih.gov/pubmed/25817535.

126. Justice AE, Winkler TW, Feitosa MF, Graff M, Fisher VA, Young K, et al. Genome-wide meta-analysis of 241,258 adults accounting for smoking behaviour identifies novel loci for obesity traits [Journal Article]. Nat Commun. 2017;8:14977. Available from: https://www.ncbi.nlm.nih.gov/pubmed/28443625.

127. Grigsby P, Elhammali A, Ruiz F, Markovina S, McLellan MD, Miller CA, et al. Clinical outcomes and differential effects of PI3K pathway mutation in obese versus non-obese patients with cervical cancer [Journal Article]. Oncotarget. 2018;9(3):4061–4073. Available from: https://www.ncbi.nlm.nih.gov/pubmed/29423104.

128. Huang X, Liu G, Guo J, Su Z. The PI3K/AKT pathway in obesity and type 2 diabetes [Journal Article]. Int J Biol Sci. 2018;14(11):1483–1496. Available from: https://www.ncbi.nlm.nih.gov/pubmed/30263000.

129. Couto Alves A, De Silva NMG, Karhunen V, Sovio U, Das S, Taal HR, et al. GWAS on longitudinal growth traits reveals different genetic factors influencing infant, child, and adult BMI [Journal Article]. Sci Adv. 2019;5(9):eaaw3095. Available from: https://www.ncbi.nlm.nih.gov/pubmed/31840077.

130. Voges D, Zwickl P, Baumeister W. The 26S proteasome: a molecular machine designed for controlled proteolysis [Journal Article]. Annu Rev Biochem. 1999;68:1015–68. Available from: https://www.ncbi.nlm.nih.gov/pubmed/10872471.

131. Livneh I, Cohen-Kaplan V, Cohen-Rosenzweig C, Avni N, Ciechanover A. The life cycle of the 26S proteasome: from birth, through regulation and function, and onto its death [Journal Article]. Cell Res. 2016;26(8):869–85. Available from: https://www.ncbi.nlm.nih.gov/pubmed/27444871.

132. Collins GA, Goldberg AL. The Logic of the 26S Proteasome [Journal Article]. Cell. 2017;169(5):792–806. Available from: https://www.ncbi.nlm.nih.gov/pubmed/28525752.

133. Groll M, Bajorek M, Kohler A, Moroder L, Rubin DM, Huber R, et al. A gated channel into the proteasome core particle [Journal Article]. Nat Struct Biol. 2000;7(11):1062–7. Available from: https://www.ncbi.nlm.nih.gov/pubmed/11062564.

134. Kohler A, Cascio P, Leggett DS, Woo KM, Goldberg AL, Finley D. The axial channel of the proteasome core particle is gated by the Rpt2 ATPase and controls both substrate entry and product release [Journal Article]. Mol Cell. 2001;7(6):1143–52. Available from: https://www.ncbi.nlm.nih.gov/pubmed/11430818.

135. Smith DM, Chang SC, Park S, Finley D, Cheng Y, Goldberg AL. Docking of the proteasomal ATPases’ carboxyl termini in the 20S proteasome’s alpha ring opens the gate for substrate entry [Journal Article]. Mol Cell. 2007;27(5):731–44. Available from: https://www.ncbi.nlm.nih.gov/pubmed/17803938.

136. Baumeister W, Walz J, Zuhl F, Seemuller E. The proteasome: paradigm of a self-compartmentalizing protease [Journal Article]. Cell. 1998;92(3):367–80. Available from: https://www.ncbi.nlm.nih.gov/pubmed/9476896.

137. Groll M, Heinemeyer W, Jager S, Ullrich T, Bochtler M, Wolf DH, et al. The catalytic sites of 20S proteasomes and their role in subunit maturation: a mutational and crystallographic study [Journal Article]. Proc Natl Acad Sci U S A. 1999;96(20):10976–83. Available from: https://www.ncbi.nlm.nih.gov/pubmed/10500111.

138. Groettrup M, Soza A, Eggers M, Kuehn L, Dick TP, Schild H, et al. A role for the proteasome regulator PA28alpha in antigen presentation [Journal Article]. Nature. 1996;381(6578):166–8. Available from: https://www.ncbi.nlm.nih.gov/pubmed/8610016.

139. de Graaf N, van Helden MJ, Textoris-Taube K, Chiba T, Topham DJ, Kloetzel PM, et al. PA28 and the proteasome immunosubunits play a central and independent role in the production of MHC class I-binding peptides in vivo [Journal Article]. Eur J Immunol. 2011;41(4):926–35. Available from: https://www.ncbi.nlm.nih.gov/pubmed/21360704.

140. Raule M, Cerruti F, Benaroudj N, Migotti R, Kikuchi J, Bachi A, et al. PA28alphabeta reduces size and increases hydrophilicity of 20S immunoproteasome peptide products [Journal Article]. Chem Biol. 2014;21(4):470–480. Available from: https://www.ncbi.nlm.nih.gov/pubmed/24631123.

141. Murata S, Takahama Y, Kasahara M, Tanaka K. The immunoproteasome and thymoproteasome: functions, evolution and human disease [Journal Article]. Nat Immunol. 2018;19(9):923–931. Available from: https://www.ncbi.nlm.nih.gov/pubmed/30104634.

142. Ferrington DA, Gregerson DS. Immunoproteasomes: structure, function, and antigen presentation [Journal Article]. Prog Mol Biol Transl Sci. 2012;109:75–112. Available from: https://www.ncbi.nlm.nih.gov/pubmed/22727420.

143. Basler M, Kirk CJ, Groettrup M. The immunoproteasome in antigen processing and other immunological functions [Journal Article]. Curr Opin Immunol. 2013;25(1):74–80. Available from: https://www.ncbi.nlm.nih.gov/pubmed/23219269.

144. McCarthy MK, Weinberg JB. The immunoproteasome and viral infection: a complex regulator of inflammation [Journal Article]. Front Microbiol. 2015;6:21. Available from: https://www.ncbi.nlm.nih.gov/pubmed/25688236.

145. Sun J, Luan Y, Xiang D, Tan X, Chen H, Deng Q, et al. The 11S Proteasome Subunit PSME3 Is a Positive Feedforward Regulator of NF-kappaB and Important for Host Defense against Bacterial Pathogens [Journal Article]. Cell Rep. 2016;14(4):737–749. Available from: https://www.ncbi.nlm.nih.gov/pubmed/26776519.

146. Mitchell S, Mercado EL, Adelaja A, Ho JQ, Cheng QJ, Ghosh G, et al. An NFkappaB Activity Calculator to Delineate Signaling Crosstalk: Type I and II Interferons Enhance NFkappaB via Distinct Mechanisms [Journal Article]. Front Immunol. 2019;10:1425. Available from: https://www.ncbi.nlm.nih.gov/pubmed/31293585.

147. Dumitrescu L, Carty CL, Taylor K, Schumacher FR, Hindorff LA, Ambite JL, et al. Genetic determinants of lipid traits in diverse populations from the population architecture using genomics and epidemiology (PAGE) study [Journal Article]. PLoS Genet. 2011;7(6):e1002138. Available from: https://www.ncbi.nlm.nih.gov/pubmed/21738485.

148. Rotimi CN, Bentley AR, Doumatey AP, Chen G, Shriner D, Adeyemo A. The genomic landscape of African populations in health and disease [Journal Article]. Hum Mol Genet. 2017;26(R2):R225–R236. Available from: https://www.ncbi.nlm.nih.gov/pubmed/28977439.

149. Choudhury A, Aron S, Sengupta D, Hazelhurst S, Ramsay M. African genetic diversity provides novel insights into evolutionary history and local adaptations [Journal Article]. Hum Mol Genet. 2018;27(R2):R209–R218. Available from: https://www.ncbi.nlm.nih.gov/pubmed/29741686.

150. Mogil LS, Andaleon A, Badalamenti A, Dickinson SP, Guo X, Rotter JI, et al. Genetic architecture of gene expression traits across diverse populations [Journal Article]. PLoS Genet. 2018;14(8):e1007586. Available from: https://www.ncbi.nlm.nih.gov/pubmed/30096133.

151. Bien SA, Wojcik GL, Hodonsky CJ, Gignoux CR, Cheng I, Matise TC, et al. The Future of Genomic Studies Must Be Globally Representative: Perspectives from PAGE [Journal Article]. Annu Rev Genomics Hum Genet. 2019;20:181–200. Available from: https://www.ncbi.nlm.nih.gov/pubmed/30978304.

152. Bentley AR, Callier SL, Rotimi CN. Evaluating the promise of inclusion of African ancestry populations in genomics [Journal Article]. NPJ Genom Med. 2020;5:5. Available from: https://www.ncbi.nlm.nih.gov/pubmed/32140257.

153. Atkinson EG, Audesse AJ, Palacios JA, Bobo DM, Webb AE, Ramachandran S, et al. No Evidence for Recent Selection at FOXP2 among Diverse Human Populations [Journal Article]. Cell. 2018;174(6):1424–1435 e15. Available from: https://www.ncbi.nlm.nih.gov/pubmed/30078708.

154. Sugden LA, Atkinson EG, Fischer AP, Rong S, Henn BM, Ramachandran S. Localization of adaptive variants in human genomes using averaged one-dependence estimation [Journal Article]. Nat Commun. 2018;9(1):703. Available from: https://www.ncbi.nlm.nih.gov/pubmed/29459739.

155. Need AC, Goldstein DB. Next generation disparities in human genomics: concerns and remedies [Journal Article]. Trends Genet. 2009;25(11):489–94. Available from: https://www.ncbi.nlm.nih.gov/pubmed/19836853.

156. Shi H, Kichaev G, Pasaniuc B. Contrasting the Genetic Architecture of 30 Complex Traits from Summary Association Data [Journal Article]. Am J Hum Genet. 2016;99(1):139–53. Available from: https://www.ncbi.nlm.nih.gov/pubmed/27346688.

157. Johnson R, Shi H, Pasaniuc B, Sankararaman S. A unifying framework for joint trait analysis under a non-infinitesimal model [Journal Article]. Bioinformatics. 2018;34(13):i195–i201. Available from: https://www.ncbi.nlm.nih.gov/pubmed/29949958.

158. Ray D, Boehnke M. Methods for meta-analysis of multiple traits using GWAS summary statistics [Journal Article]. Genet Epidemiol. 2018;42(2):134–145. Available from: https://www.ncbi.nlm.nih.gov/pubmed/29226385.

159. Turchin MC, Stephens M. Bayesian multivariate reanalysis of large genetic studies identifies many new associations [Journal Article]. PLoS Genet. 2019;15(10):e1008431. Available from: https://www.ncbi.nlm.nih.gov/pubmed/31596850.

160. Urbut SM, Wang G, Carbonetto P, Stephens M. Flexible statistical methods for estimating and testing effects in genomic studies with multiple conditions [Journal Article]. Nat Genet. 2019;51(1):187–195. Available from: https://www.ncbi.nlm.nih.gov/pubmed/30478440.

161. Kichaev G, Yang WY, Lindstrom S, Hormozdiari F, Eskin E, Price AL, et al. Integrating functional data to prioritize causal variants in statistical fine-mapping studies [Journal Article]. PLoS Genet. 2014;10(10):e1004722. Available from: https://www.ncbi.nlm.nih.gov/pubmed/25357204.

162. Chen W, Larrabee BR, Ovsyannikova IG, Kennedy RB, Haralambieva IH, Poland GA, et al. Fine Mapping Causal Variants with an Approximate Bayesian Method Using Marginal Test Statistics [Journal Article]. Genetics. 2015;200(3):719–36. Available from: https://www.ncbi.nlm.nih.gov/pubmed/25948564.

163. Benner C, Spencer CC, Havulinna AS, Salomaa V, Ripatti S, Pirinen M. FINEMAP: efficient variable selection using summary data from genome-wide association studies [Journal Article]. Bioinformatics. 2016;32(10):1493–501. Available from: https://www.ncbi.nlm.nih.gov/pubmed/26773131.

164. Hormozdiari F, van de Bunt M, Segre AV, Li X, Joo JWJ, Bilow M, et al. Colocalization of GWAS and eQTL Signals Detects Target Genes [Journal Article]. Am J Hum Genet. 2016;99(6):1245–1260. Available from: https://www.ncbi.nlm.nih.gov/pubmed/27866706.

165. Zhu Z, Zhang F, Hu H, Bakshi A, Robinson MR, Powell JE, et al. Integration of summary data from GWAS and eQTL studies predicts complex trait gene targets [Journal Article]. Nat Genet. 2016;48(5):481–7. Available from: https://www.ncbi.nlm.nih.gov/pubmed/27019110.

166. Wen X, Pique-Regi R, Luca F. Integrating molecular QTL data into genome-wide genetic association analysis: Probabilistic assessment of enrichment and colocalization [Journal Article]. PLoS Genet. 2017;13(3):e1006646. Available from: https://www.ncbi.nlm.nih.gov/pubmed/28278150.

167. Giambartolomei C, Zhenli Liu J, Zhang W, Hauberg M, Shi H, Boocock J, et al. A Bayesian framework for multiple trait colocalization from summary association statistics [Journal Article]. Bioinformatics. 2018;34(15):2538–2545. Available from: https://www.ncbi.nlm.nih.gov/pubmed/29579179.

168. Wallace C. Eliciting priors and relaxing the single causal variant assumption in colocalisation analyses [Journal Article]. PLoS Genet. 2020;16(4):e1008720. Available from: https://www.ncbi.nlm.nih.gov/pubmed/32310995.

169. He M, Xu M, Zhang B, Liang J, Chen P, Lee JY, et al. Meta-analysis of genome-wide association studies of adult height in East Asians identifies 17 novel loci [Journal Article]. Hum Mol Genet. 2015;24(6):1791–800. Available from: https://www.ncbi.nlm.nih.gov/pubmed/25429064.

170. Tachmazidou I, Suveges D, Min JL, Ritchie GRS, Steinberg J, Walter K, et al. Whole-Genome Sequencing Coupled to Imputation Discovers Genetic Signals for Anthropometric Traits [Journal Article]. Am J Hum Genet. 2017;100(6):865–884. Available from: https://www.ncbi.nlm.nih.gov/pubmed/28552196.

171. Akiyama M, Ishigaki K, Sakaue S, Momozawa Y, Horikoshi M, Hirata M, et al. Characterizing rare and low-frequency height-associated variants in the Japanese population [Journal Article]. Nat Commun. 2019;10(1):4393. Available from: https://www.ncbi.nlm.nih.gov/pubmed/31562340.

172. Saeed S, Bonnefond A, Tamanini F, Mirza MU, Manzoor J, Janjua QM, et al. Loss-of-function mutations in ADCY3 cause monogenic severe obesity [Journal Article]. Nat Genet. 2018;50(2):175–179. Available from: https://www.ncbi.nlm.nih.gov/pubmed/29311637.

173. Cao Y, Li L, Xu M, Feng Z, Sun X, Lu J, et al. The ChinaMAP analytics of deep whole genome sequences in 10,588 individuals [Journal Article]. Cell Res. 2020;30(9):717–731. Available from: https://www.ncbi.nlm.nih.gov/pubmed/32355288.

174. Kichaev G, Bhatia G, Loh PR, Gazal S, Burch K, Freund MK, et al. Leveraging Polygenic Functional Enrichment to Improve GWAS Power [Journal Article]. Am J Hum Genet. 2019;104(1):65–75. Available from: https://www.ncbi.nlm.nih.gov/pubmed/30595370.

175. Berndt SI, Gustafsson S, Magi R, Ganna A, Wheeler E, Feitosa MF, et al. Genome-wide meta-analysis identifies 11 new loci for anthropometric traits and provides insights into genetic architecture [Journal Article]. Nat Genet. 2013;45(5):501–12. Available from: https://www.ncbi.nlm.nih.gov/pubmed/23563607.

176. Shungin D, Winkler TW, Croteau-Chonka DC, Ferreira T, Locke AE, Magi R, et al. New genetic loci link adipose and insulin biology to body fat distribution [Journal Article]. Nature. 2015;518(7538):187–196. Available from: https://www.ncbi.nlm.nih.gov/pubmed/25673412.

177. Graff M, Scott RA, Justice AE, Young KL, Feitosa MF, Barata L, et al. Genome-wide physical activity interactions in adiposity - A meta-analysis of 200,452 adults [Journal Article]. PLoS Genet. 2017;13(4):e1006528. Available from: https://www.ncbi.nlm.nih.gov/pubmed/28448500.

178. Pulit SL, Stoneman C, Morris AP, Wood AR, Glastonbury CA, Tyrrell J, et al. Meta-analysis of genome-wide association studies for body fat distribution in 694 649 individuals of European ancestry [Journal Article]. Hum Mol Genet. 2019;28(1):166–174. Available from: https://www.ncbi.nlm.nih.gov/pubmed/30239722.

179. Lotta LA, Wittemans LBL, Zuber V, Stewart ID, Sharp SJ, Luan J, et al. Association of Genetic Variants Related to Gluteofemoral vs Abdominal Fat Distribution With Type 2 Diabetes, Coronary Disease, and Cardiovascular Risk Factors [Journal Article]. JAMA. 2018;320(24):2553–2563. Available from: https://www.ncbi.nlm.nih.gov/pubmed/30575882.

180. Justice AE, Karaderi T, Highland HM, Young KL, Graff M, Lu Y, et al. Protein-coding variants implicate novel genes related to lipid homeostasis contributing to body-fat distribution [Journal Article]. Nat Genet. 2019;51(3):452–469. Available from: https://www.ncbi.nlm.nih.gov/pubmed/30778226.

181. Zhu Z, Guo Y, Shi H, Liu CL, Panganiban RA, Chung W, et al. Shared genetic and experimental links between obesity-related traits and asthma subtypes in UK Biobank [Journal Article]. J Allergy Clin Immunol. 2020;145(2):537–549. Available from: https://www.ncbi.nlm.nih.gov/pubmed/31669095.

182. Chu AY, Deng X, Fisher VA, Drong A, Zhang Y, Feitosa MF, et al. Multiethnic genome-wide meta-analysis of ectopic fat depots identifies loci associated with adipocyte development and differentiation [Journal Article]. Nat Genet. 2017;49(1):125–130. Available from: https://www.ncbi.nlm.nih.gov/pubmed/27918534.

183. Winkler TW, Justice AE, Graff M, Barata L, Feitosa MF, Chu S, et al. The Influence of Age and Sex on Genetic Associations with Adult Body Size and Shape: A Large-Scale Genome-Wide Interaction Study [Journal Article]. PLoS Genet. 2015;11(10):e1005378. Available from: https://www.ncbi.nlm.nih.gov/pubmed/26426971.

184. Lango Allen H, Estrada K, Lettre G, Berndt SI, Weedon MN, Rivadeneira F, et al. Hundreds of variants clustered in genomic loci and biological pathways affect human height [Journal Article]. Nature. 2010;467(7317):832–8. Available from: http://www.ncbi.nlm.nih.gov/entrez/query.fcgi?cmd=Retrieve&db=PubMed&dopt=Citation&list_uids=20881960.

185. Rueger S, McDaid A, Kutalik Z. Evaluation and application of summary statistic imputation to discover new height-associated loci [Journal Article]. PLoS Genet. 2018;14(5):e1007371. Available from: https://www.ncbi.nlm.nih.gov/pubmed/29782485.

186. Manning AK, Hivert MF, Scott RA, Grimsby JL, Bouatia-Naji N, Chen H, et al. A genome-wide approach accounting for body mass index identifies genetic variants influencing fasting glycemic traits and insulin resistance [Journal Article]. Nat Genet. 2012;44(6):659–69. Available from: https://www.ncbi.nlm.nih.gov/pubmed/22581228.

187. Spracklen CN, Chen P, Kim YJ, Wang X, Cai H, Li S, et al. Association analyses of East Asian individuals and trans-ancestry analyses with European individuals reveal new loci associated with cholesterol and triglyceride levels [Journal Article]. Hum Mol Genet. 2017;26(9):1770–1784. Available from: https://www.ncbi.nlm.nih.gov/pubmed/28334899.

188. Ng MC, Hester JM, Wing MR, Li J, Xu J, Hicks PJ, et al. Genome-wide association of BMI in African Americans [Journal Article]. Obesity (Silver Spring). 2012;20(3):622–7. Available from: https://www.ncbi.nlm.nih.gov/pubmed/21701570.

