## Supplementary Material for "Pathway Analysis within Multiple Human Ancestries Reveals Novel Signals for Epistasis in Complex Traits"

### Contents

**1 Data Quality Control Procedures for UK Biobank** **2**

**2 Hypergeometric Enrichment Analyses** **2**

**3 Supplementary Figures** **3**

**4 Supplementary Tables** **25**

References **35**

### 1 Data Quality Control Procedures for UK Biobank

The results presented in the main text made use of genotyped chip data released from the UK Biobank [1]. Here, we focus on subgroups from the UK Biobank that we created first by grouping individuals based on their self-identified ancestries: “African”, “British”, “Caribbean”, “Chinese”, “Indian”, and “Pakistani”. We then conducted the following quality control procedures on each of these subgroups. To start, we removed SNPs by checking three sets of criteria. First, we removed SNPs with minor allele frequency (MAF) less than 1%. Second, we removed SNPs with missingness greater than 5%. Lastly, we removed SNPs not in Hardy-Weinberg Equilibrium (Fisher's exact test $p$-value $> 10^{-6}$). We then removed individuals by checking four sets of criteria. First, we removed individuals with greater than 5% genotype missingness. Second, we removed individuals if they were a 3$^{\text{rd}}$ degree relative (or more) to someone else in the dataset — specifically, the KING relatedness values provided with the UK Biobank dataset were used to identify related individuals, and one individual from every pair of 3$^{\text{rd}}$ degree or more relatives was randomly removed. Third, individuals were also removed if they were tagged by any of the following three flags from the UK Biobank dataset: `het.missing.outliers`, `putative.sex.chromosome.aneuploidy`, and `excess.relatives`. Lastly, individuals were removed if they were determined to be outliers via principal component analysis (PCA) — this was conducted by running `FlashPCA` (version 2.1) [2] in `R` on each population subgroup separately and identifying individuals that had any of their top six principal component (PC) values greater than seven standard deviations away from the mean. We refer to conducting PCA on each subgroup separately as “local” PCA to help distinguish from the alternative setup of conducting PCA on the entire dataset jointly, which we refer to as “global” PCA. Note that global PCs were only used to create the PCA plots shown in Supplementary Figure 1, while local PCs were used for quality control and to adjust for population structure in the MAPIT-R analyses.

After the first round of quality control procedures, we then proceeded to impute missing genotypes for each population subgroup. To conduct this imputation, we uploaded our population subgroups to the University of Michigan Imputation Server [3] and used the following options: `Minimac3` for the imputation software, 1000 Genomes (phase 3 version 5) for the reference panel, and `Eagle` (version 2.3) for the phasing software. Completed imputed files were then downloaded from the Imputation Server afterwards and treated to one last round of quality control. Variants with imputation quality scores less than .3 were removed, and variants that had genotype missingness greater than 0% were also removed. Information on the final forms of our UK Biobank population subgroups can be found in Supplementary Table 1 and information on the number of individuals dropped at each relevant quality control step can be found in Supplementary Table 2.

### 2 Hypergeometric Enrichment Analyses

Hypergeometric analyses were conducted by counting the number of times a given gene is present among the significant pathways identified by MAPIT-R for a given database-phenotype-subgroup combination versus the number of times the gene is present among the background distribution of pathways from the same database. For the size-restriction analyses, pathways with greater than 1,000 SNPs were removed. The 1,000 SNP threshold was chosen to balance between the total number of pathways included in analyses and the average pathway size across all subgroups (see Supplementary Figure 13 and 14 for the relationship between MAPIT-R $p$-values and pathway sizes). Note that for a number of size thresholds ranging from 500 to 1,500 SNPs, results regarding the proteasome gene families remain robust (see Supplementary Figure 22).

### 3 Supplementary Figures

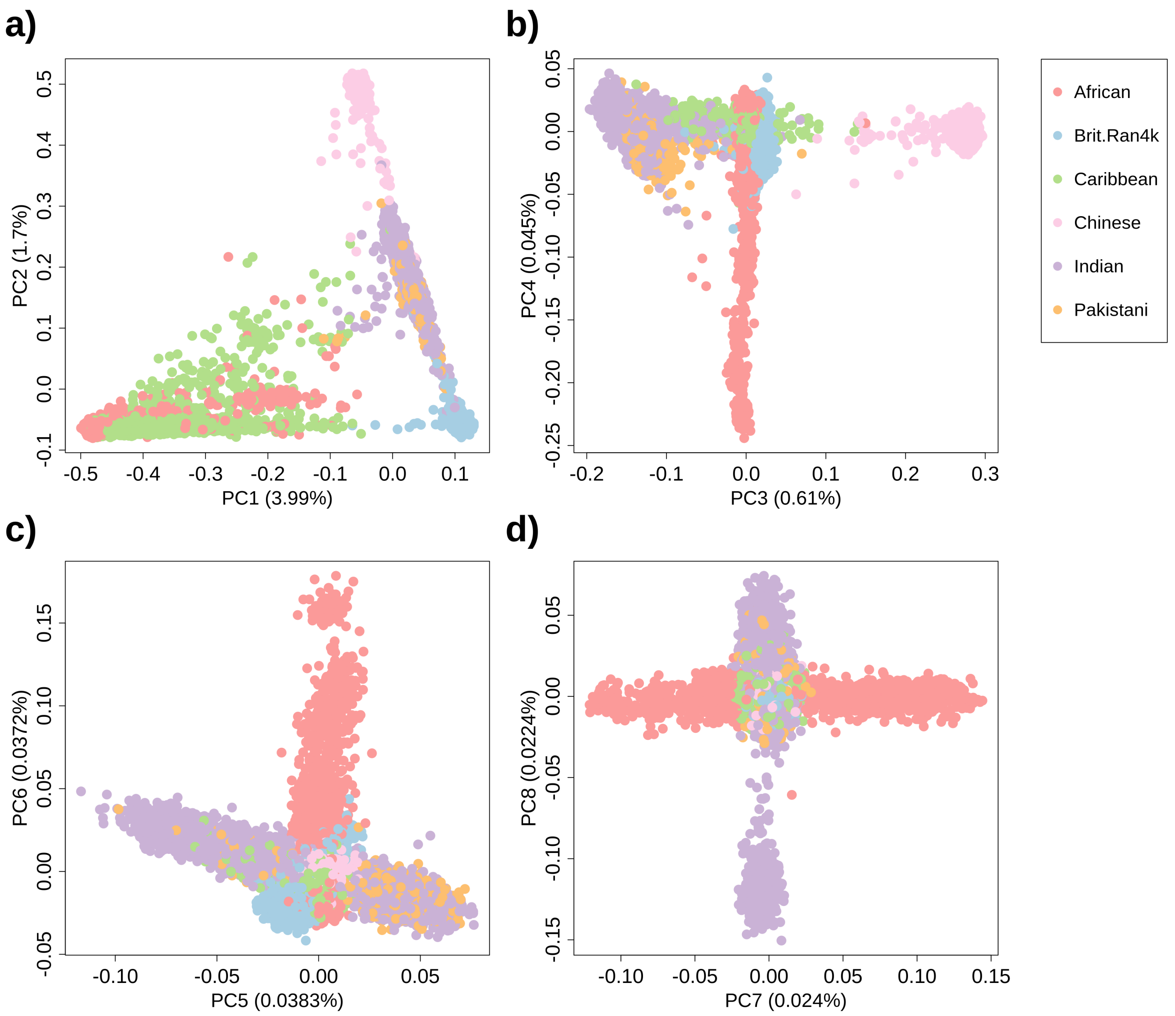


**Supplementary Figure 1. Global principal component analysis (PCA) of the different ancestry-specific subgroups in the UK Biobank analyzed in this study.** Here, subgroups in the UK Biobank included the following self-identified ancestries: "African", "British", "Caribbean", "Chinese", "Indian", and "Pakistani" (see legend to the right of plots). The figure shows the top eight global principal components (PCs) for each subgroup plotted against one another. The comparisons include: **(a)** PC1 vs. PC2, **(b)** PC3 vs. PC4, **(c)** PC5 vs. PC6, and **(d)** PC7 vs. PC8. PCA was conducted using `FlashPCA` [2]. Here, "global" refers to running PCA on all subgroups jointly. This is in contrast to "local" PCA in which we conduct principal component analysis on each subgroup separately. Note that global PCs were only used to create this figure, while local PCs were used for quality control and to adjust for population structure in the MAPIT-R analyses (see 'UK Biobank Data' in Materials and Methods and 'Data Quality Control Procedures for UK Biobank' in Supplementary Note). On each axis, we also list the percent of total variation explained by each PC in parentheses.

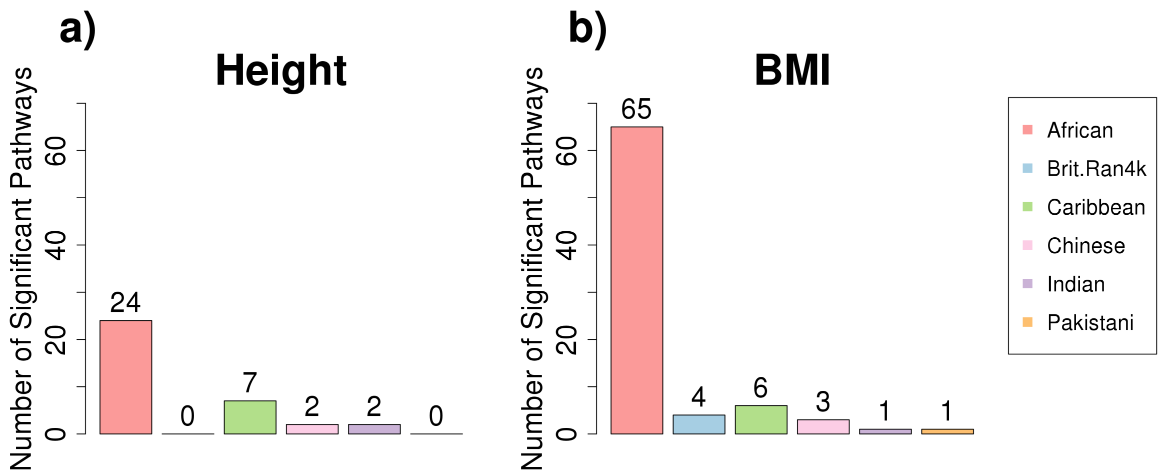


**Supplementary Figure 2. Number of REACTOME pathways identified by MAPIT-R that have significant marginal epistatic effects within (a) standing height and (b) body mass index (BMI) per subgroup in the UK Biobank.** Here, subgroups in the UK Biobank included individuals based on their self-identified ancestries: "African", "British", "Caribbean", "Chinese", "Indian", and "Pakistani" (see legend to the right of panel **(b)**). Genome-wide significance was determined by using Bonferroni-corrected $p$-value thresholds based on the number of pathways tested in each database-phenotype-subgroup combination (see Supplementary Table 1). Across all database-phenotype combinations, the African subgroup has the largest numbers of significant pathways. For lists of the specific significant pathways per database-phenotype-subgroup combination, see Supplementary Table 3.

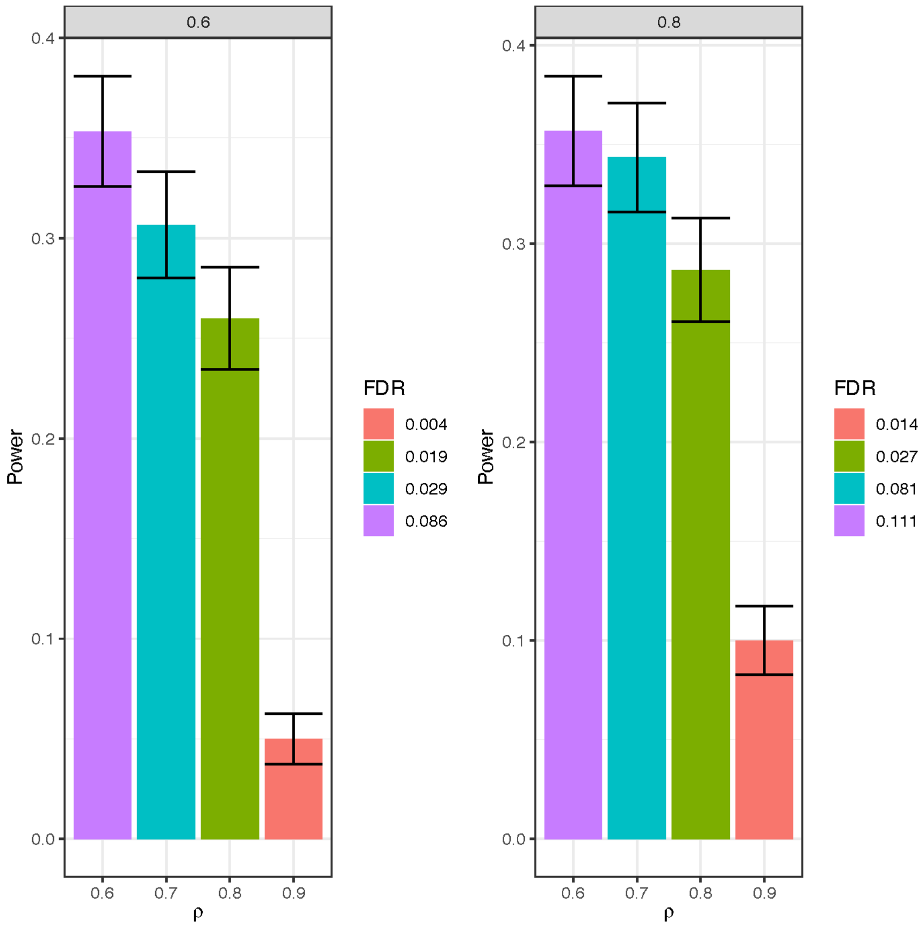


**Supplementary Figure 3. Barplots of MAPIT-R performance under different simulated genetic architectures**. Here, we show results from running simulations of MAPIT-R under various architectures using a simulation scheme similar to that used in Crawford et al. [4]. For all simulations, 3,000 random individuals from the British subgroup and 10,000 random SNPs from chromosome 1 were used. We simulated continuous phenotypes using the following model: $\mathbf{y} = \mathbf{X}_{\mathcal{C}}\boldsymbol{\beta}_{\mathcal{C}} + \mathbf{Z}\mathbf{u} + \mathbf{W}\boldsymbol{\alpha} + \boldsymbol{\epsilon}$, where $\mathbf{X}_{\mathcal{C}}$ is a genotype matrix of randomly chosen causal SNPs with additive effects, $\mathbf{Z}$ is a covariate matrix representing population structure, $\mathbf{W}$ is a design matrix that contains SNP-by-SNP interaction designations, and $\mathbf{u}$, $\boldsymbol{\beta}$, and $\boldsymbol{\alpha}$ are all effect sizes that are drawn from standard normal distributions, or $\mathcal{N}(\mathbf{0}, \mathbf{I})$. The random noise term, $\boldsymbol{\epsilon}$, is also drawn as a standard normal variable. To create the matrix $\mathbf{W}$, we first randomly create 10 enriched pathways, each containing 50 causal SNPs. We then allow each causal SNP to interact with 200 SNPs from the additive genetic background. This is done by simply taking the Hadamard (element-wise) product between genotypic vectors. In total, 100,000 interactions are created per simulation (i.e., 10 pathways each with 50 SNPs that interact with 200 partnering SNPs). We scale both the additive and pairwise interaction effects so that collectively they explain a fixed proportion of the phenotypic variance. Namely, the additive effects make up $\rho\%$ while the epistatic terms make up the remaining $(1-\rho)\%$. Alternatively, the proportion of broad-sense heritability that is explained by additive effects is said to be $\mathbb{V}[\mathbf{X}_{\mathcal{C}}\boldsymbol{\beta}_{\mathcal{C}}] = H^2\rho$, while the proportion of heritability that is detailed by interactions is $H^2(1-\rho)$. Lastly, we introduce population stratification effects by using the top 10 local principal components as the covariates in $\mathbf{Z}$ and scaling their effects to account for an additional 10% of the overall phenotypic variance. Overall, we analyze 50 simulated datasets per scenario. The barplots show mean power and false discovery rates (FDR) as a function of $\rho$ for **(a)** $H^2 = 0.6$ and **(b)** $H^2 = 0.8$, respectively. Standard error bars are also included.

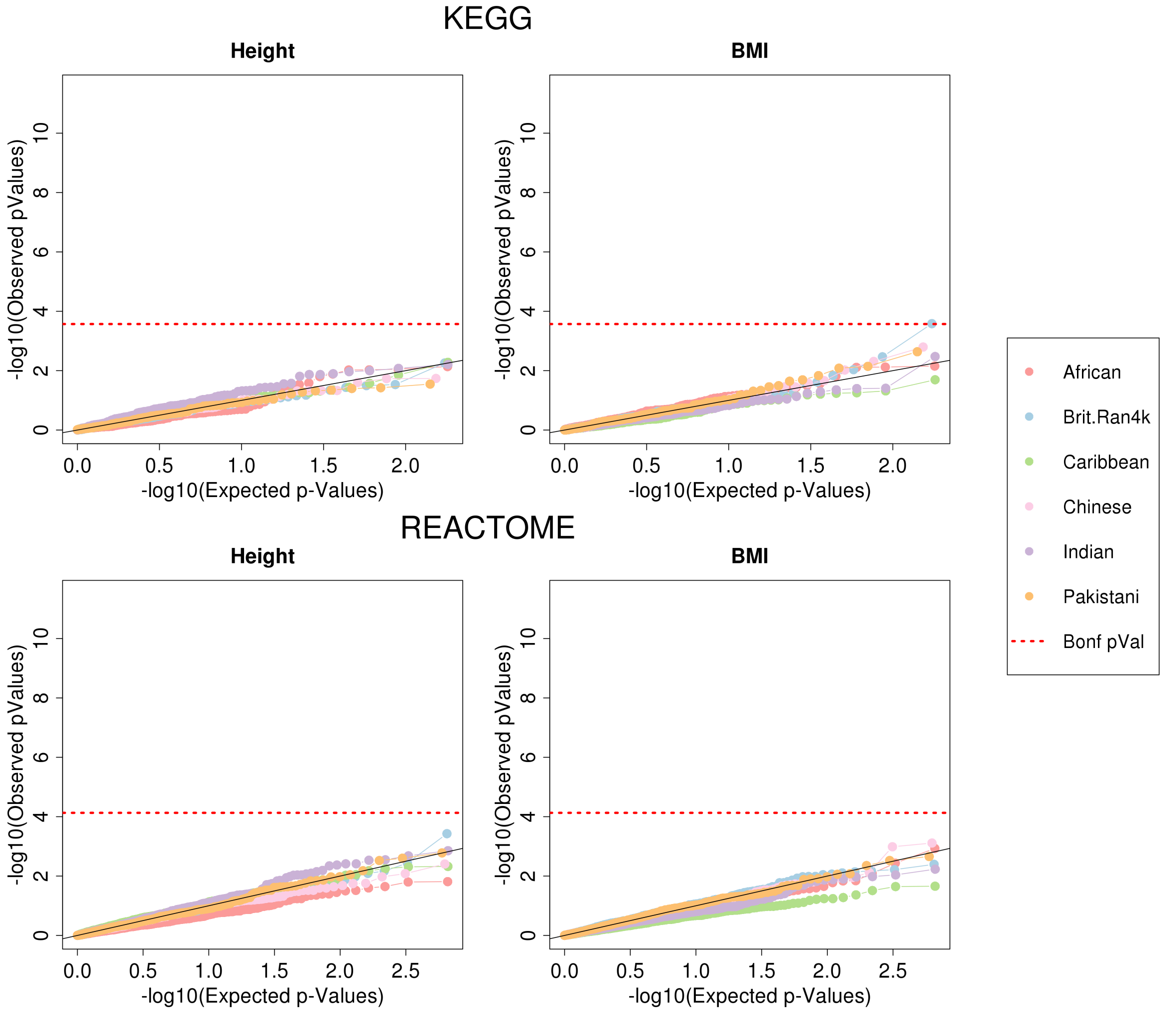


**Supplementary Figure 4. QQ-Plots of MAPIT-R $p$-values using KEGG and REACTOME pathway annotations and randomly permuted phenotypes in each ancestry-specific subgroup in the UK Biobank.** Here, we run MAPIT-R after conducting a single random permutation of either height or BMI measurements. Note that traits were permuted within each population subgroup 10 different times. The $x$-axis shows the -$\log_{10}$ transformed expected $p$-values, while the $y$-axis shows the -$\log_{10}$ observed $p$-values. The dotted red line is the Bonferroni-corrected $p$-value threshold based on the number of pathways tested per database-phenotype combination (Supplementary Table 1). Subgroups in the UK Biobank included individuals based on their self-identified ancestries: “African”, “British”, “Caribbean”, “Chinese”, “Indian”, and “Pakistani” (see legend to the right of plots). Overall, we find that MAPIT-R properly controls for the type 1 error rate across varying significance cutoff thresholds (Supplementary Table 4).

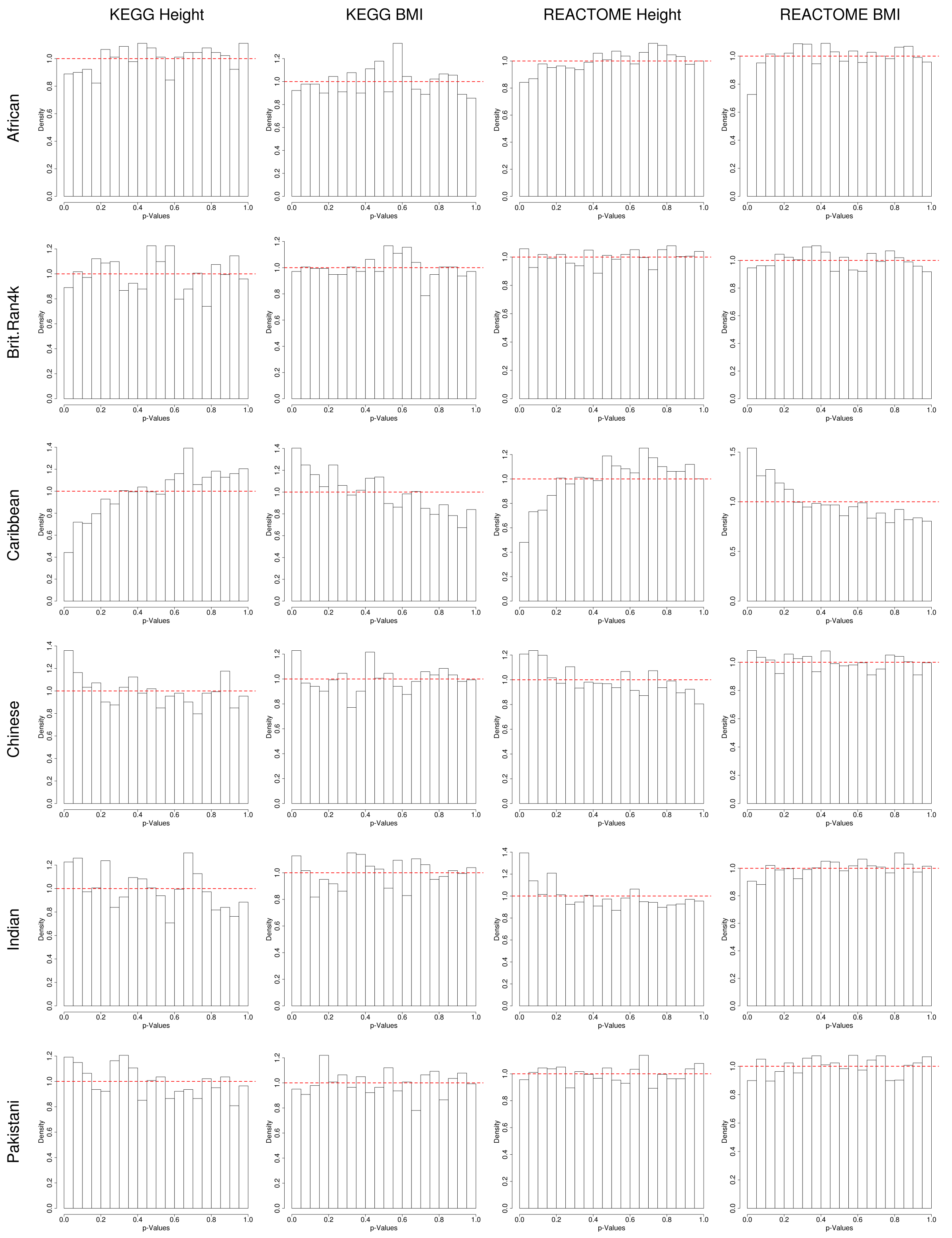


**Supplementary Figure 5. Histograms of MAPIT-R *p*-values using KEGG and REACTOME pathway annotations and randomly permuted phenotypes in each ancestry-specific subgroup in the UK Biobank.** Here, we run MAPIT-R after conducting a single random permutation of either height or BMI measurements. Note that traits were independently permuted within each population subgroup ten different times. Subgroups in the UK Biobank included individuals based on their self-identified ancestries: "African", "British", "Caribbean", "Chinese", "Indian", and "Pakistani" (ordered here from top-to-bottom). The dotted red line corresponds to a uniform distribution of *p*-values. Overall, we find that MAPIT-R properly controls for type 1 error rate at varying significance cutoff thresholds (Supplementary Table 4).

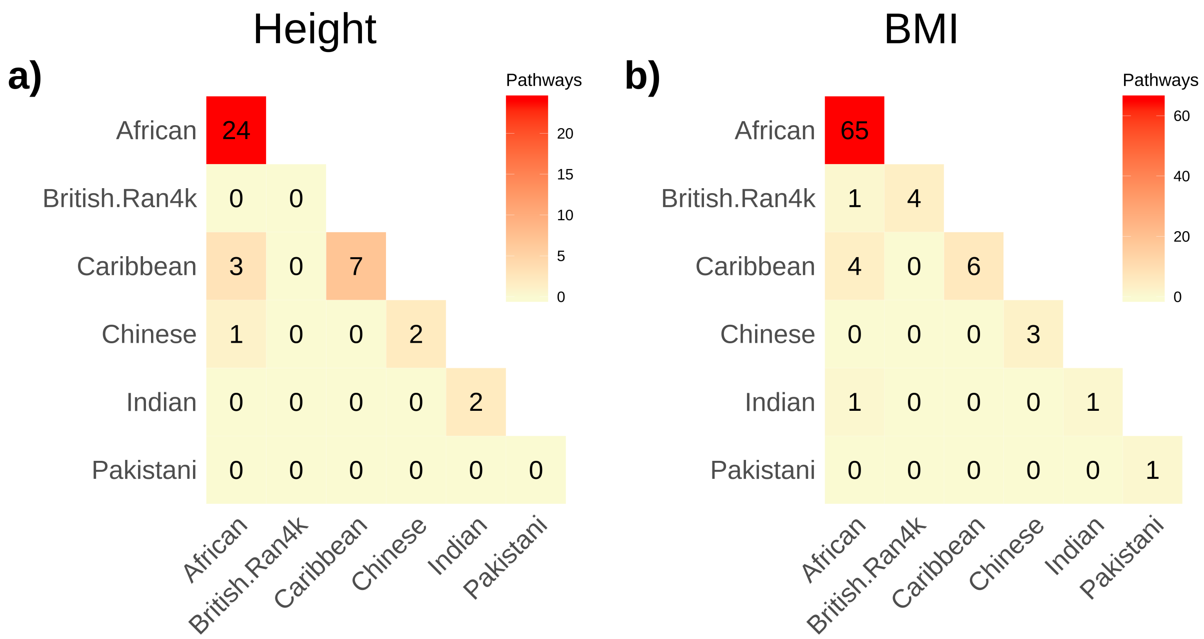


**Supplementary Figure 6. Heatmaps depicting the overlap of MAPIT-R significant REACTOME pathways for (a) standing height and (b) body mass index (BMI) between the different ancestry-specific subgroups in the UK Biobank**. Here, subgroups in the UK Biobank included individuals based on their self-identified ancestries: "African", "British", "Caribbean", "Chinese", "Indian", and "Pakistani" (ordered here from top-to-bottom and left-to-right). Genome-wide significance was determined by using Bonferroni-corrected $p$-value thresholds based on the number of pathways tested in each database-phenotype-subgroup combination (see Supplementary Table 1). The diagonal shows the total number of genome-wide significant pathways per subgroup. We observe that significant pathways observed in non-African subgroups overlap more often with pathways from the African subgroup than they do with pathways from the other, remaining non-African subgroups. Results for both phenotypes in the KEGG database can be seen in Figure 2 in the main text.

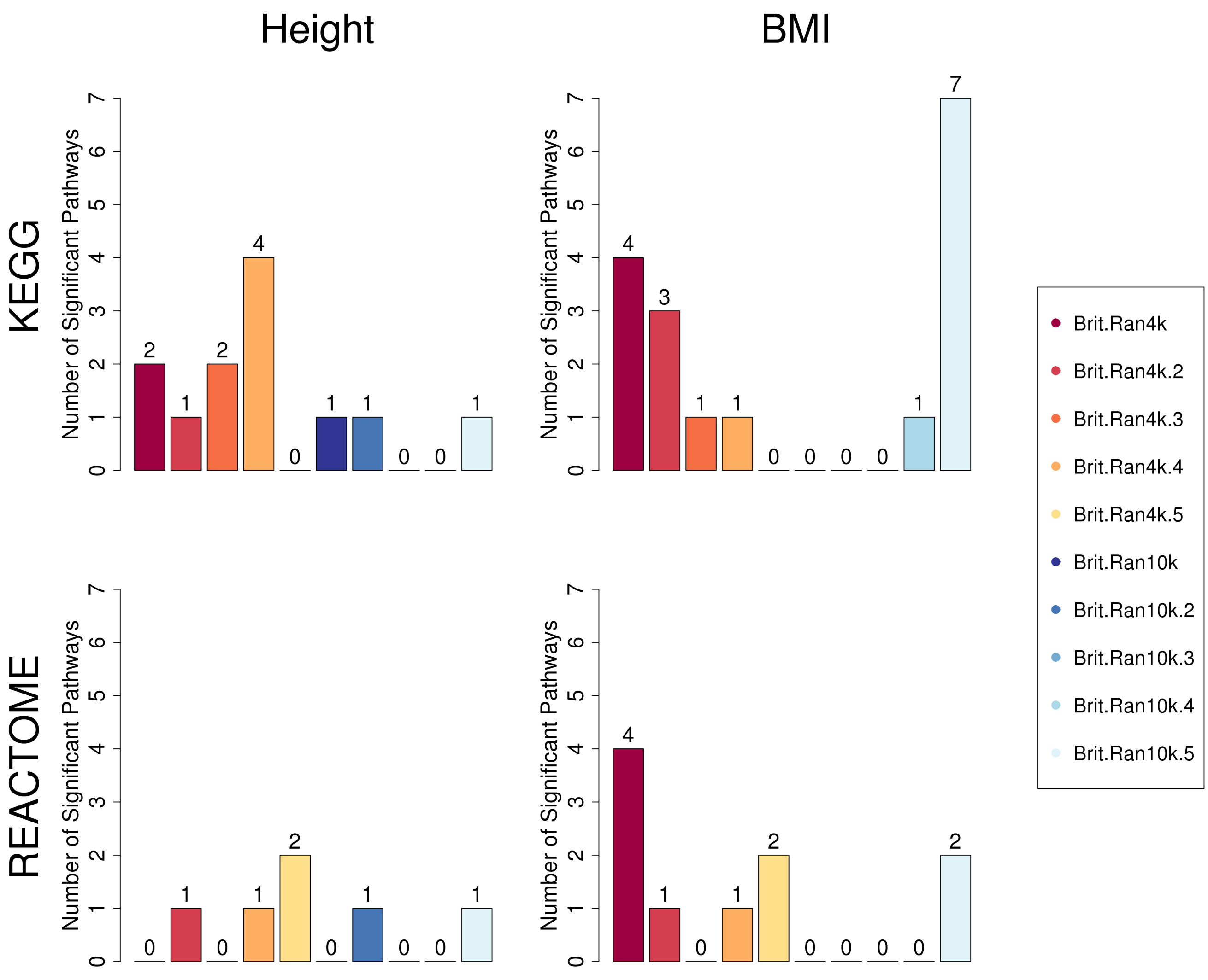


**Supplementary Figure 7. Number of KEGG and REACTOME pathways identified by MAPIT-R that have significant marginal epistatic effects within standing height and body mass index (BMI) per British replicate subgroup in the UK Biobank.** Here, the random subgroups were created by subsampling either $N = 4{,}000$ (reds and oranges) or 10,000 (blues) individuals from the full British cohort. In the legend, replicates are marked as numbers after the root tag name (e.g., `Brit.Ran4k.4` denotes the fourth replicate in the $N = 4{,}000$ subsampling scheme). Genome-wide significance was determined by using Bonferroni-corrected $p$-value thresholds based on the number of pathways tested in each database-phenotype-subgroup combination (see Supplementary Table 1). Note that most replicate runs did not produce many significant results. See Supplementary Figures 8-12 and Supplementary Tables 8-9 for other analyses using the 4,000 and 10,000 British replicate subgroups.

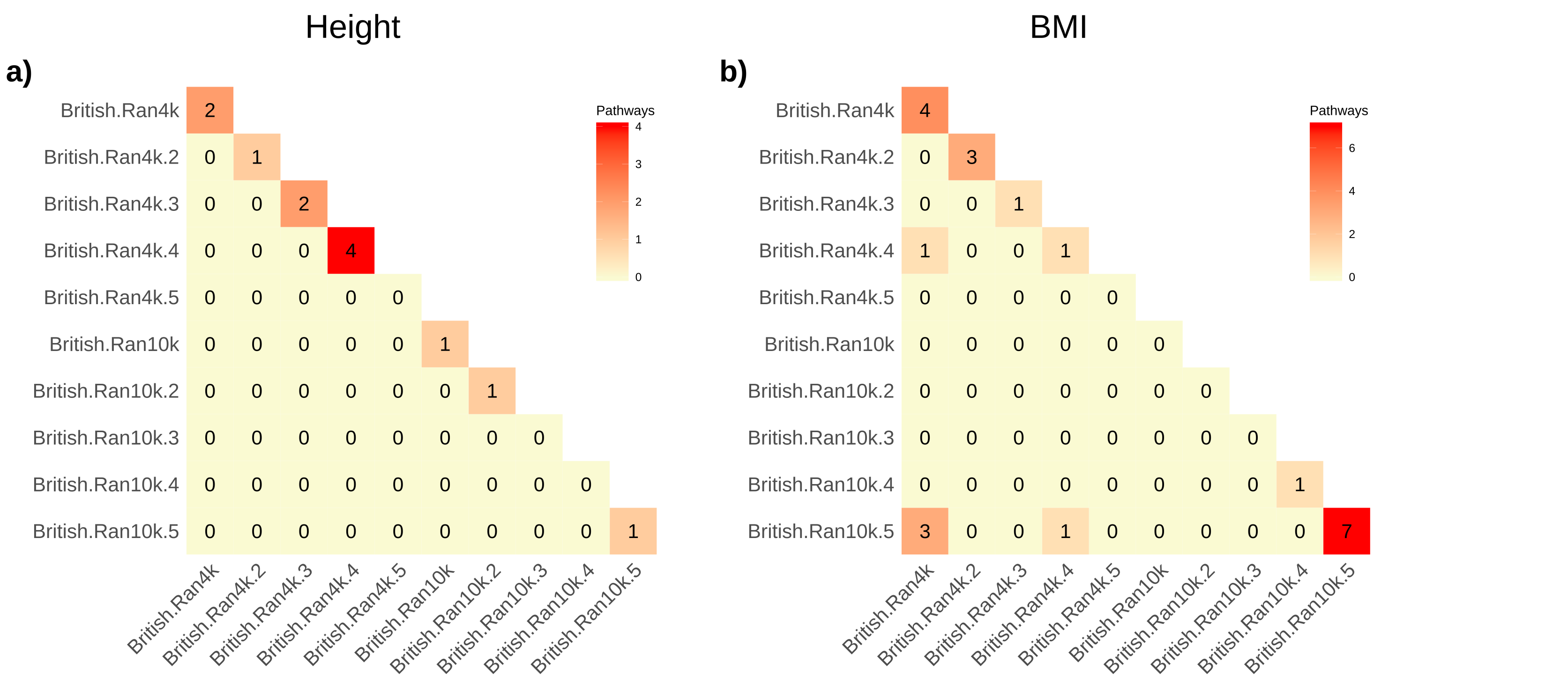


**Supplementary Figure 8. Heatmaps depicting the overlap of MAPIT-R significant KEGG pathways for (a) standing height and (b) body mass index (BMI) per British replicate subgroup in the UK Biobank**. Here, the random subgroups were created by subsampling either $N = 4{,}000$ or 10,000 individuals from the full British cohort. In the legend, replicates are marked as numbers after the root tag name (e.g., `Brit.Ran4k.4` denotes the fourth replicate in the $N = 4{,}000$ subsampling scheme). Genome-wide significance was determined by using Bonferroni-corrected $p$-value thresholds based on the number of pathways tested in each database-phenotype-subgroup combination (see Supplementary Table 1). The diagonal shows the total number of genome-wide significant pathways per subgroup. We do not observe much overlap between enriched pathways between the replicate subgroups — even though they all consist of individuals of the same ancestry. See Supplementary Figures 7,9-12 and Supplementary Tables 8-9 for other analyses using the 4,000 and 10,000 British replicate subgroups.

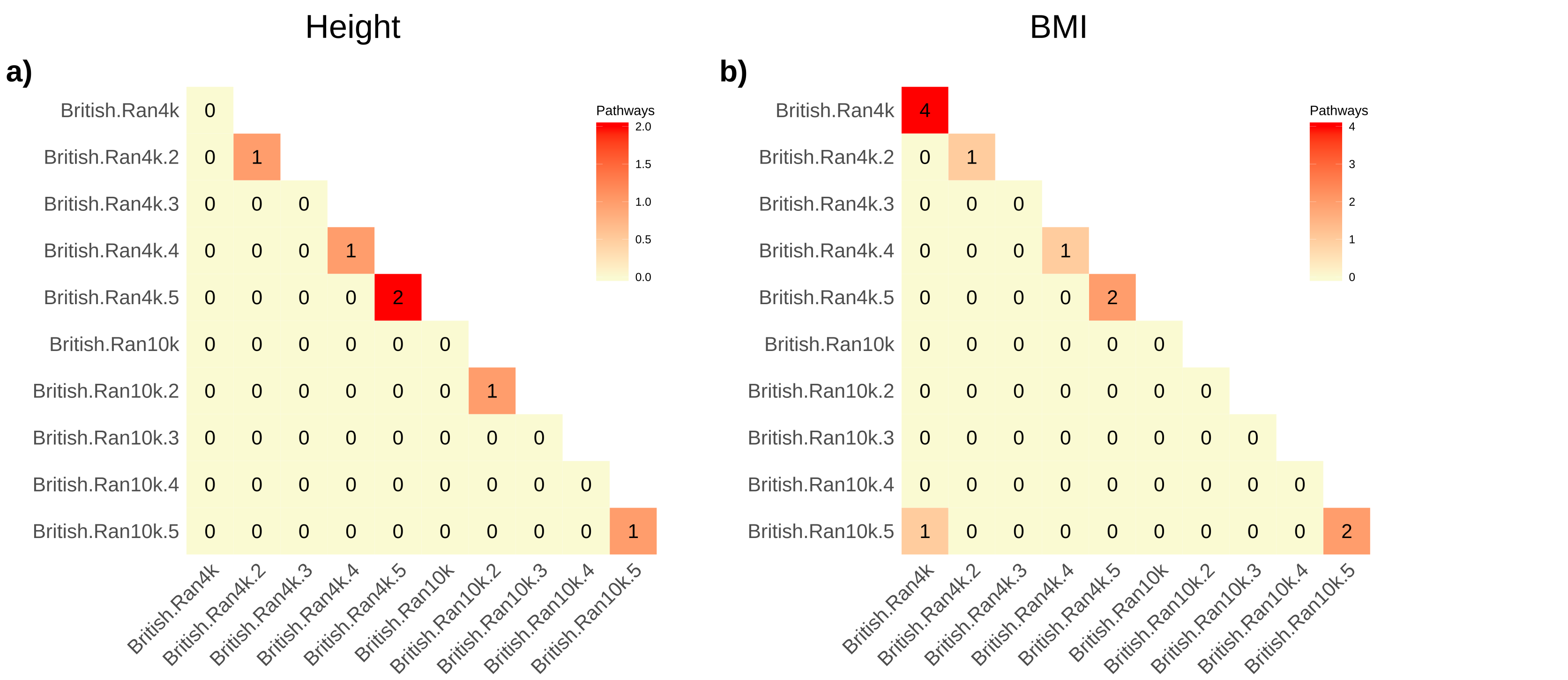


**Supplementary Figure 9. Heatmaps depicting the overlap of MAPIT-R significant REACTOME pathways for (a) standing height and (b) body mass index (BMI) per British replicate subgroup in the UK Biobank**. Here, the random subgroups were created by subsampling either $N = 4{,}000$ or 10,000 individuals from the full British cohort. In the legend, replicates are marked as numbers after the root tag name (e.g., `Brit.Ran4k.4` denotes the fourth replicate in the $N = 4{,}000$ subsampling scheme). Genome-wide significance was determined by using Bonferroni-corrected $p$-value thresholds based on the number of pathways tested in each database-phenotype-subgroup combination (see Supplementary Table 1). The diagonal shows the total number of genome-wide significant pathways per subgroup. We do not observe much overlap between enriched pathways between the replicate subgroups — even though they all consist of individuals of the same ancestry. See Supplementary Figures 7-8,10-12 and Supplementary Tables 8-9 for other analyses using the 4,000 and 10,000 British replicate subgroups.

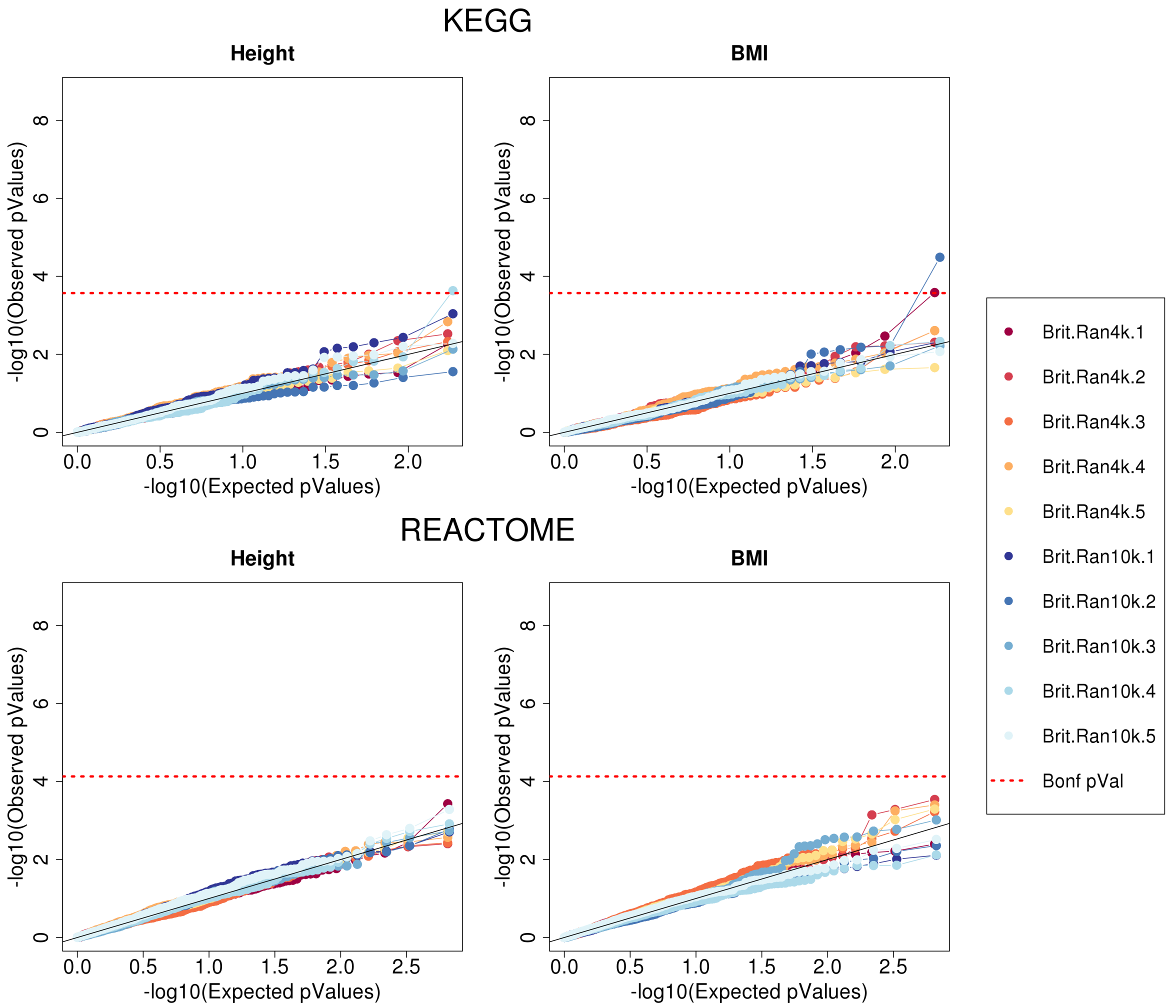


**Supplementary Figure 10. QQ-Plots of MAPIT-R $p$-values using KEGG and REACTOME pathway annotations and randomly permuted phenotypes in British replicate subgroups from the UK Biobank.** Here, we run MAPIT-R after conducting a single random permutation of either height or BMI measurements. Note that traits were permuted within each population subgroup ten different times. The $x$-axis shows the -$\log_{10}$ transformed expected $p$-values, while the $y$-axis shows the -$\log_{10}$ observed $p$-values. The random subgroups were created by subsampling either $N = 4{,}000$ (reds and oranges) or 10,000 (blues) individuals from the full British cohort. In the legend, replicates are marked as numbers after the root tag name (e.g., `Brit.Ran4k.4` denotes the fourth replicate in the $N = 4{,}000$ subsampling scheme). The dotted red line is the Bonferroni-corrected $p$-value threshold based on the number of pathways tested per database-phenotype combination (Supplementary Table 1). Overall, we find that MAPIT-R continues to exhibit well-calibrated behavior under the null hypothesis and properly controls for type 1 error rate. See Supplementary Figures 7-9,11-12 and Supplementary Tables 8-9 for other analyses using the 4,000 and 10,000 British replicate subgroups.

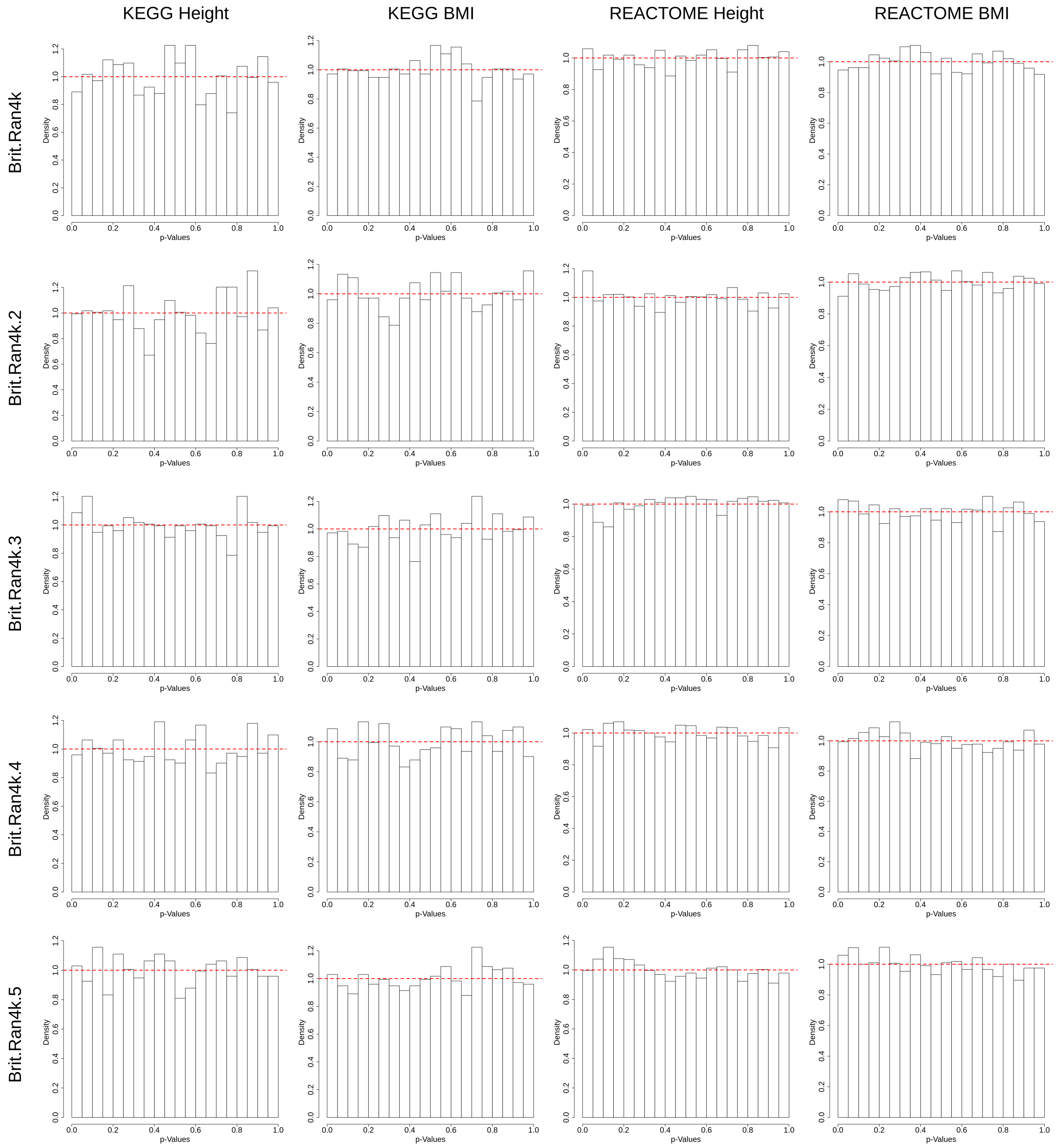


**Supplementary Figure 11. Histograms of MAPIT-R $p$-values using KEGG and REACTOME pathway annotations and randomly permuted phenotypes in each British 4,000 replicate subgroup in the UK Biobank.** Here, we run MAPIT-R after conducting a single random permutation of either height or BMI measurements. Note that traits were independently permuted within each population subgroup ten different times. The random subgroups were created by subsampling $N =$ 4,000 individuals from the full British cohort. In the row labels, replicates are marked as numbers after the root tag name (e.g., `Brit.Ran4k.4` denotes the fourth replicate in the $N =$ 4,000 subsampling scheme). The dotted red line corresponds to a uniform distribution of $p$-values. Overall, we find that MAPIT-R continues to exhibit well-calibrated behavior under the null hypothesis. See Supplementary Figures 7-10,12 and Supplementary Tables 8-9 for other analyses using the 4,000 and 10,000 British replicate subgroups.

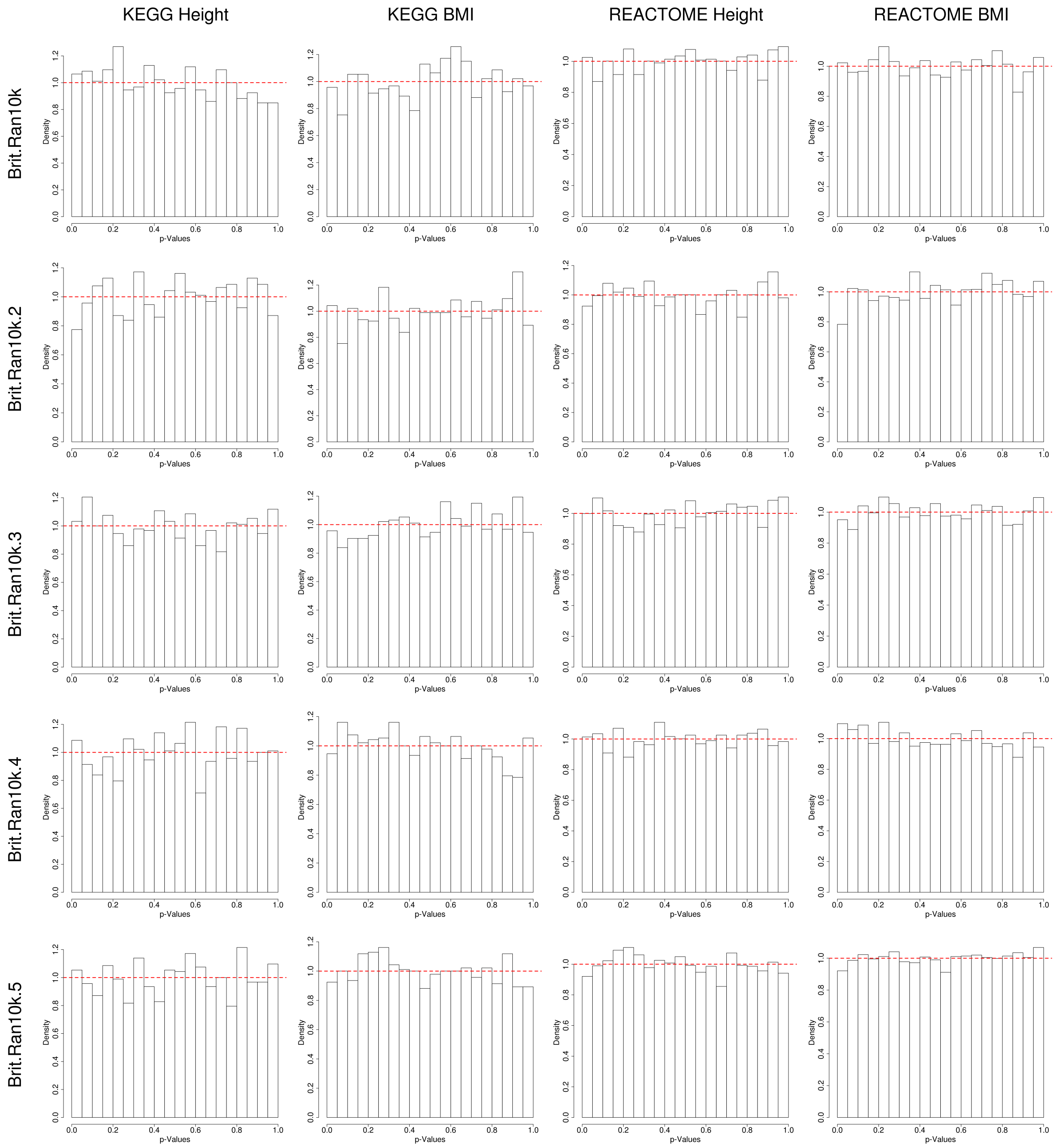


**Supplementary Figure 12. Histograms of MAPIT-R *p*-values using KEGG and REACTOME pathway annotations and randomly permuted phenotypes in each British 10,000 replicate subgroup in the UK Biobank.** Here, we run MAPIT-R after conducting a single random permutation of either height or BMI measurements. Note that traits were independently permuted within each population subgroup ten different times. The random subgroups were created by subsampling $N =$ 10,000 individuals from the full British cohort. In the row labels, replicates are marked as numbers after the root tag name (e.g., `Brit.Ran10k.4` denotes the fourth replicate in the $N$ = 10,000 subsampling scheme). The dotted red line corresponds to a uniform distribution of $p$-values. Overall, we find that MAPIT-R continues to exhibit well-calibrated behavior under the null hypothesis. See Supplementary Figures 7-11 and Supplementary Tables 8-9 for other analyses using the 4,000 and 10,000 British replicate subgroups.

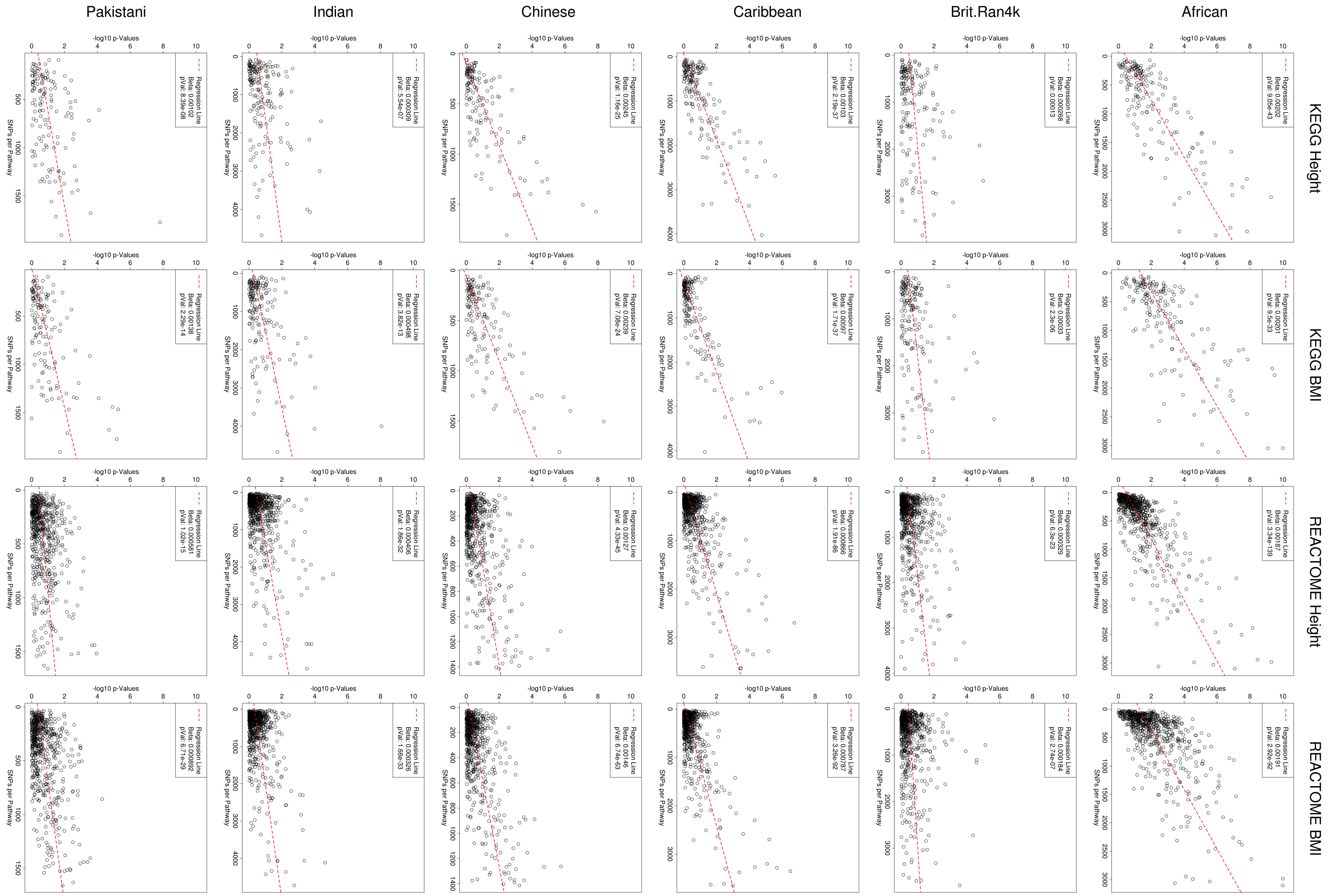


**Supplementary Figure 13. Scatterplots comparing the MAPIT-R *p*-values as function of the number of SNPs annotated within each pathway for KEGG and REACTOME in height and body mass index (BMI) across each ancestry-specific subgroup in the UK Biobank.** Here, subgroups in the UK Biobank included individuals based on their self-identified ancestries: "African", "British", "Caribbean", "Chinese", "Indian", and "Pakistani" (ordered here from right-to-left). The dotted red line represents the best fit using a univariate model, and the legend provides the corresponding regression coefficient and its associated *p*-value. We observe that for most combinations there is a significant relationship between the MAPIT-R *p*-value and the number of SNPs present in a pathway. This follows our hypothesis that combining SNPs together in a joint analysis might provide greater power to detect marginal epistasis than analyzing each SNP independently. We note, however, that these results appear to not solely be driven just by the presence or absence of large SNP counts — conducting this same analysis on one of our sets of permuted phenotypes, we find very few significant relationships between MAPIT-R *p*-values and pathway SNP counts (Supplementary Figure 14).

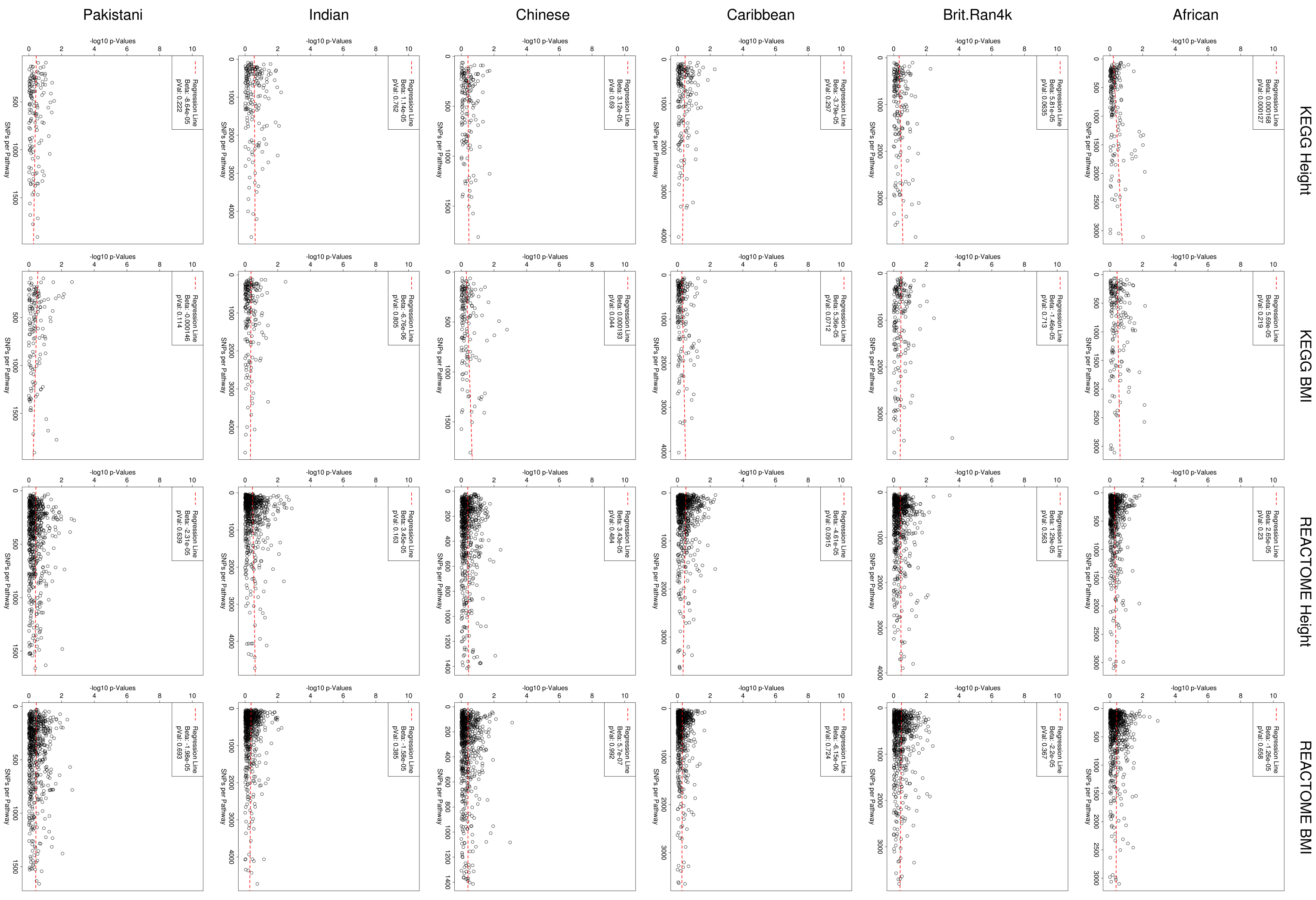


**Supplementary Figure 14. Scatterplots comparing the MAPIT-R *p*-values as function of the number of SNPs annotated within each pathway for KEGG and REACTOME using permuted phenotypes across each ancestry-specific subgroup in the UK Biobank.** Here, we run MAPIT-R after conducting a single random permutation of either height or BMI measurements. Note that traits were independently permuted within each population subgroup ten different times. The dotted red line represents the best fit using a univariate model, and the legend provides the corresponding regression coefficient and its associated *p*-value. Subgroups in the UK Biobank included individuals based on their self-identified ancestries: “African”, “British”, “Caribbean”, “Chinese”, “Indian”, and “Pakistani” (ordered here from right-to-left). In these permuted cases, we observe hardly any relationship between MAPIT-R results and the number of SNPs present in a pathway. For this same analysis on the original set of observed phenotypes, see Supplementary Figure 13.

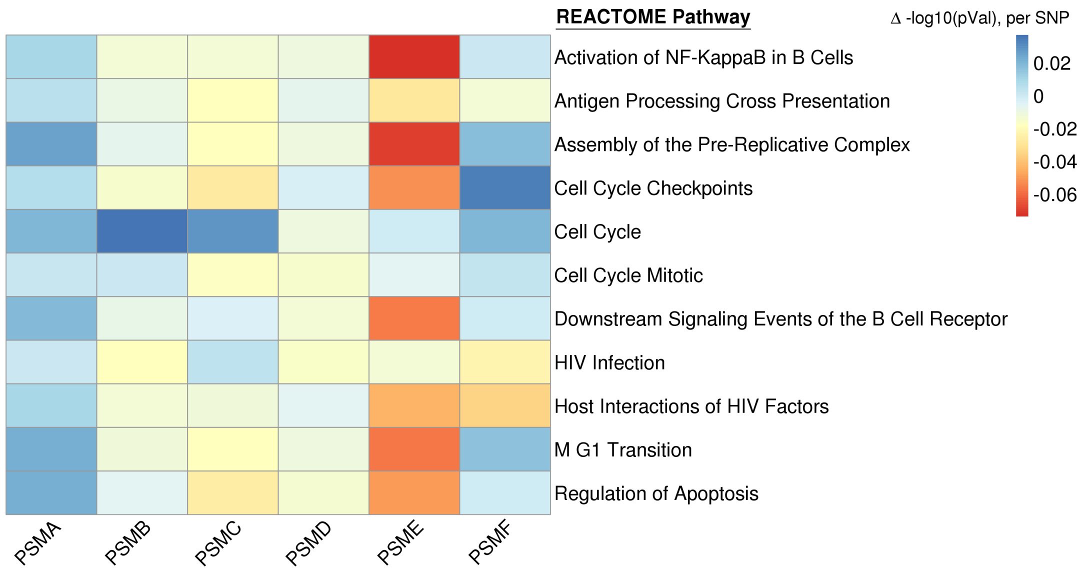


(a) Δ in MAPIT-R -$\log_{10}$ $p$-value per SNP

| Gene Family | # SNPs | REACTOME Pathway | # SNPs | $p$-Value |
|---|---|---|---|---|
| PSMA | 17 | Activation of NF-KappaB in B Cells | 465 | $1.86 \times 10^{-5}$ |
| PSMB | 74 | Antigen Processing Cross Presentation | 850 | $3.96 \times 10^{-5}$ |
| PSMC | 20 | Assembly of the Pre-Replicative Complex | 331 | $3.29 \times 10^{-5}$ |
| PSMD | 62 | Cell Cycle Checkpoints | 670 | $5.78 \times 10^{-6}$ |
| PSME | 15 | Cell Cycle | 2459 | $1.29 \times 10^{-6}$ |
| PSMF | 16 | Cell Cycle Mitotic | 1906 | $4.51 \times 10^{-5}$ |
| | | Downstream Signaling Events of the B Cell Receptor | 745 | $1.19 \times 10^{-5}$ |
| | | HIV Infection | 1346 | $7.53 \times 10^{-7}$ |
| | | Host Interactions of HIV Factors | 963 | $7.52 \times 10^{-6}$ |
| | | M G1 Transition | 458 | $3.19 \times 10^{-6}$ |
| | | Regulation of Apoptosis | 564 | $1.22 \times 10^{-5}$ |

(b) Number of SNPs in each Proteasome Gene Family and REACTOME Pathway

**Supplementary Figure 15. Results from applying a "leave-one-out" approach to MAPIT-R with proteasome gene families in body mass index (BMI) within the African subgroup in the UK Biobank. (a)** The heatmap shows the change in original MAPIT-R -$\log_{10}$ $p$-value for different REACTOME pathways when each proteasome gene family is removed one at a time in a "leave-one-out" manner. The $x$-axis shows each proteasome gene family and the $y$-axis lists each REACTOME pathway. Each column has been scaled by the number of SNPs present in the given gene family and, as a result, the heatmap specifically shows the -$\log_{10}$ $p$-value change (Δ in legend) per SNP. Raw MAPIT-R $p$-values underlying the heatmap can be found in Supplementary Table 10. **(b)** The table shows the number of SNPs present in each proteasome gene family (left), as well as the number of SNPs present in each REACTOME pathway (right). The original MAPIT-R $p$-values are also shown for each pathway (right).

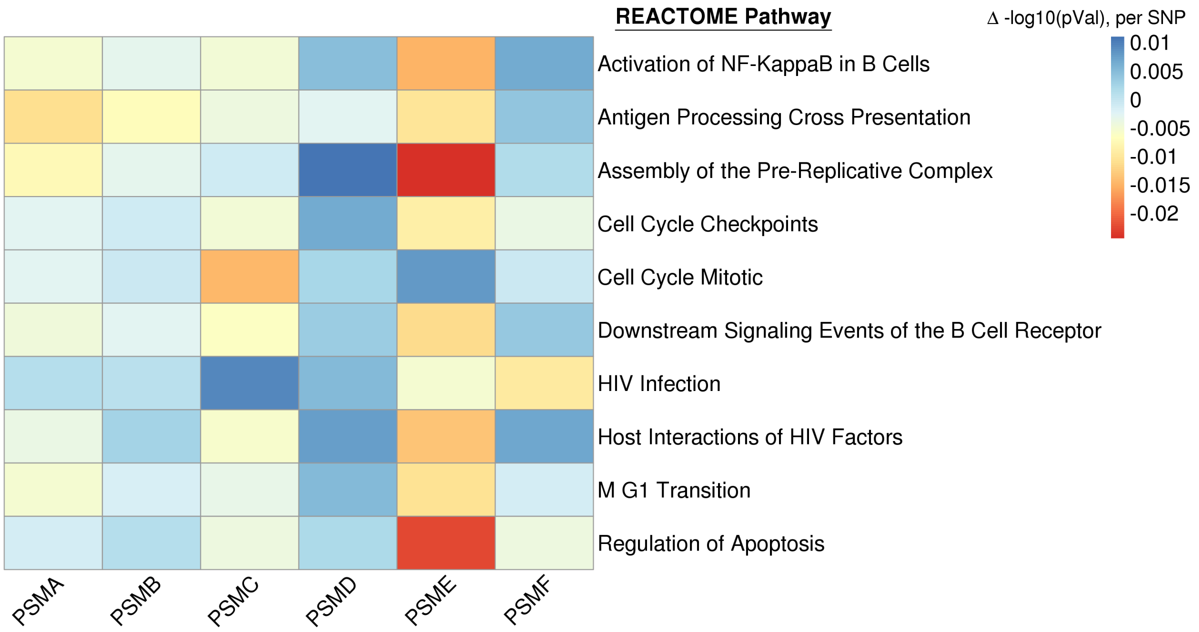


(a) Δ in MAPIT-R -$\log_{10}$ $p$-value per SNP

| Gene Family | # SNPs | REACTOME Pathway | # SNPs | $p$-Value |
|---|---|---|---|---|
| PSMA | 40 | Activation of NF-KappaB in B Cells | 756 | $2.28\times10^{-1}$ |
| PSMB | 91 | Antigen Processing Cross Presentation | 1104 | $2.48\times10^{-5}$ |
| PSMC | 29 | Assembly of the Pre-Replicative Complex | 507 | $1.84\times10^{-2}$ |
| PSMD | 101 | Cell Cycle Checkpoints | 1121 | $6.77\times10^{-2}$ |
| PSME | 25 | Cell Cycle Mitotic | 3304 | $2.34\times10^{-1}$ |
| PSMF | 26 | Downstream Signaling Events of the B Cell Receptor | 1248 | $3.25\times10^{-1}$ |
| | | HIV Infection | 2221 | $8.26\times10^{-2}$ |
| | | Host Interactions of HIV Factors | 1541 | $1.55\times10^{-2}$ |
| | | M G1 Transition | 697 | $3.02\times10^{-1}$ |
| | | Regulation of Apoptosis | 906 | $2.98\times10^{-2}$ |

(b) Number of SNPs in each Proteasome Gene Family and REACTOME Pathway

**Supplementary Figure 16. Results from applying a "leave-one-out" approach to MAPIT-R with proteasome gene families in body mass index (BMI) within 4,000 randomly subsampled individuals of British ancestry (`Brit.Ran4k`) in the UK Biobank. (a)** The heatmap shows the change in original MAPIT-R -$\log_{10}$ $p$-value for different REACTOME pathways when each proteasome gene family is removed one at a time in a "leave-one-out" manner. The $x$-axis shows each proteasome gene family and the $y$-axis lists each REACTOME pathway. Each column has been scaled by the number of SNPs present in the given gene family and, as a result, the heatmap specifically shows the -$\log_{10}$ $p$-value change (Δ in legend) per SNP. Raw MAPIT-R $p$-values underlying the heatmap can be found in Supplementary Table 10. **(b)** The table shows the number of SNPs present in each proteasome gene family (left), as well as the number of SNPs present in each REACTOME pathway (right). The original MAPIT-R $p$-values are also shown for each pathway (right).

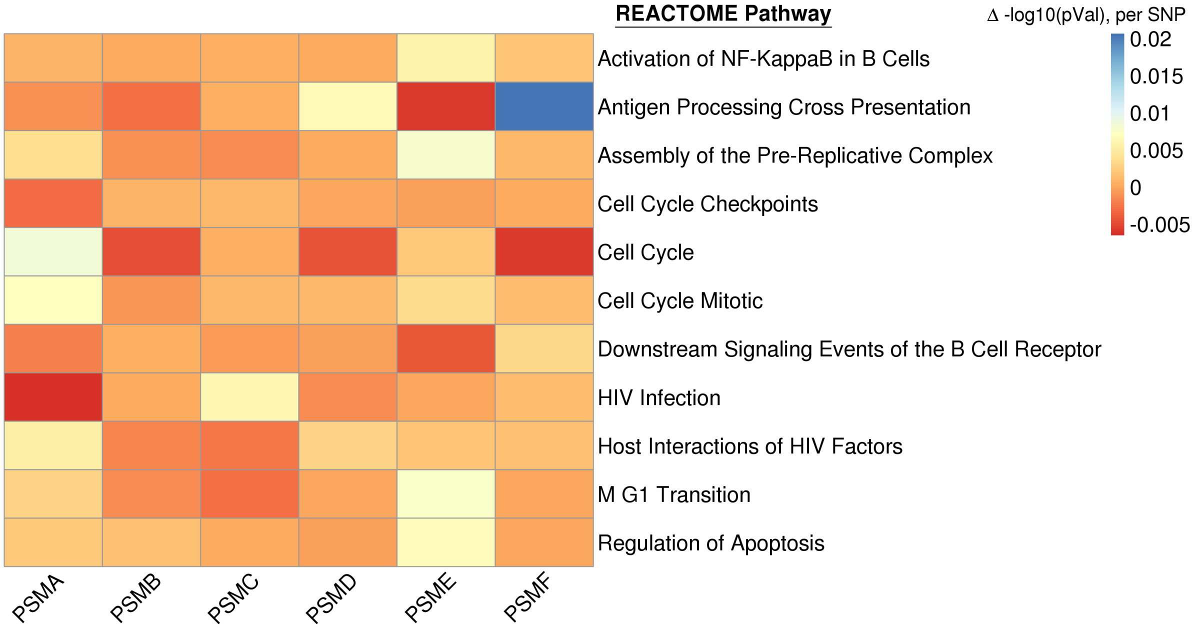


(a) Δ in MAPIT-R -$\log_{10}$ $p$-value per SNP

| Gene Family | # SNPs | REACTOME Pathway | # SNPs | $p$-Value |
|---|---|---|---|---|
| PSMA | 18 | Activation of NF-KappaB in B Cells | 507 | $9.93\times10^{-1}$ |
| PSMB | 77 | Antigen Processing Cross Presentation | 871 | $5.15\times10^{-2}$ |
| PSMC | 21 | Assembly of the Pre-Replicative Complex | 357 | $7.25\times10^{-1}$ |
| PSMD | 69 | Cell Cycle Checkpoints | 736 | $8.83\times10^{-1}$ |
| PSME | 15 | Cell Cycle | 2711 | $2.51\times10^{-1}$ |
| PSMF | 16 | Cell Cycle Mitotic | 2111 | $7.98\times10^{-1}$ |
| | | Downstream Signaling Events of the B Cell Receptor | 829 | $8.37\times10^{-1}$ |
| | | HIV Infection | 1483 | $3.11\times10^{-1}$ |
| | | Host Interactions of HIV Factors | 1055 | $6.68\times10^{-1}$ |
| | | M G1 Transition | 500 | $6.37\times10^{-1}$ |
| | | Regulation of Apoptosis | 615 | $9.42\times10^{-1}$ |

(b) Number of SNPs in each Proteasome Gene Family and REACTOME Pathway

**Supplementary Figure 17. Results from applying a "leave-one-out" approach to MAPIT-R with proteasome gene families in body mass index (BMI) within the Caribbean subgroup in the UK Biobank. (a)** The heatmap shows the change in original MAPIT-R -$\log_{10}$ $p$-value for different REACTOME pathways when each proteasome gene family is removed one at a time in a "leave-one-out" manner. The $x$-axis shows each proteasome gene family and the $y$-axis lists each REACTOME pathway. Each column has been scaled by the number of SNPs present in the given gene family and, as a result, the heatmap specifically shows the -$\log_{10}$ $p$-value change (Δ in legend) per SNP. Raw MAPIT-R $p$-values underlying the heatmap can be found in Supplementary Table 10. **(b)** The table shows the number of SNPs present in each proteasome gene family (left), as well as the number of SNPs present in each REACTOME pathway (right). The original MAPIT-R $p$-values are also shown for each pathway (right).

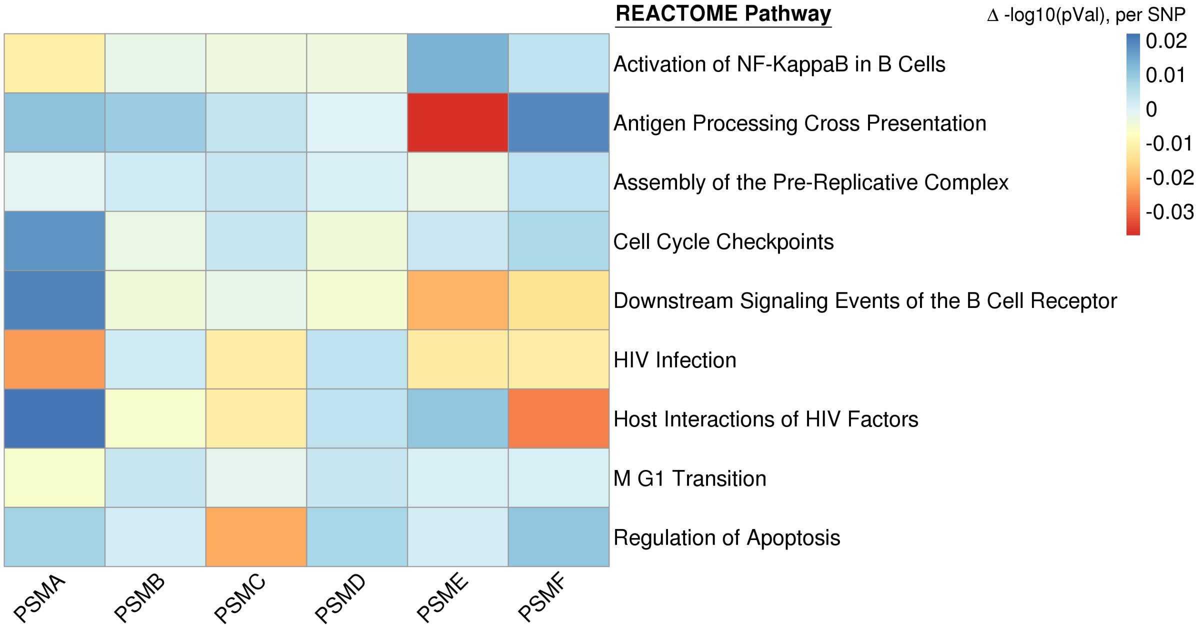


(a) Δ in MAPIT-R -$\log_{10}$ $p$-value per SNP

| Gene Family | # SNPs | REACTOME Pathway | # SNPs | $p$-Value |
|---|---|---|---|---|
| PSMA | 13 | Activation of NF-KappaB in B Cells | 433 | $5.27\times10^{-1}$ |
| PSMB | 74 | Antigen Processing Cross Presentation | 771 | $1.11\times10^{-2}$ |
| PSMC | 18 | Assembly of the Pre-Replicative Complex | 292 | $8.77\times10^{-1}$ |
| PSMD | 58 | Cell Cycle Checkpoints | 589 | $5.41\times10^{-1}$ |
| PSME | 12 | Downstream Signaling Events of the B Cell Receptor | 698 | $1.40\times10^{-1}$ |
| PSMF | 16 | HIV Infection | 1266 | $3.68\times10^{-3}$ |
| | | Host Interactions of HIV Factors | 902 | $4.24\times10^{-2}$ |
| | | M G1 Transition | 400 | $8.20\times10^{-1}$ |
| | | Regulation of Apoptosis | 527 | $2.26\times10^{-2}$ |

(b) Number of SNPs in each Proteasome Gene Family and REACTOME Pathway

**Supplementary Figure 18. Results from applying a "leave-one-out" approach to MAPIT-R with proteasome gene families in body mass index (BMI) within the Chinese subgroup in the UK Biobank. (a)** The heatmap shows the change in original MAPIT-R -$\log_{10}$ $p$-value for different REACTOME pathways when each proteasome gene family is removed one at a time in a "leave-one-out" manner. The $x$-axis shows each proteasome gene family and the $y$-axis lists each REACTOME pathway. Each column has been scaled by the number of SNPs present in the given gene family and, as a result, the heatmap specifically shows the -$\log_{10}$ $p$-value change (Δ in legend) per SNP. Raw MAPIT-R $p$-values underlying the heatmap can be found in Supplementary Table 10. **(b)** The table shows the number of SNPs present in each proteasome gene family (left), as well as the number of SNPs present in each REACTOME pathway (right). The original MAPIT-R $p$-values are also shown for each pathway (right).

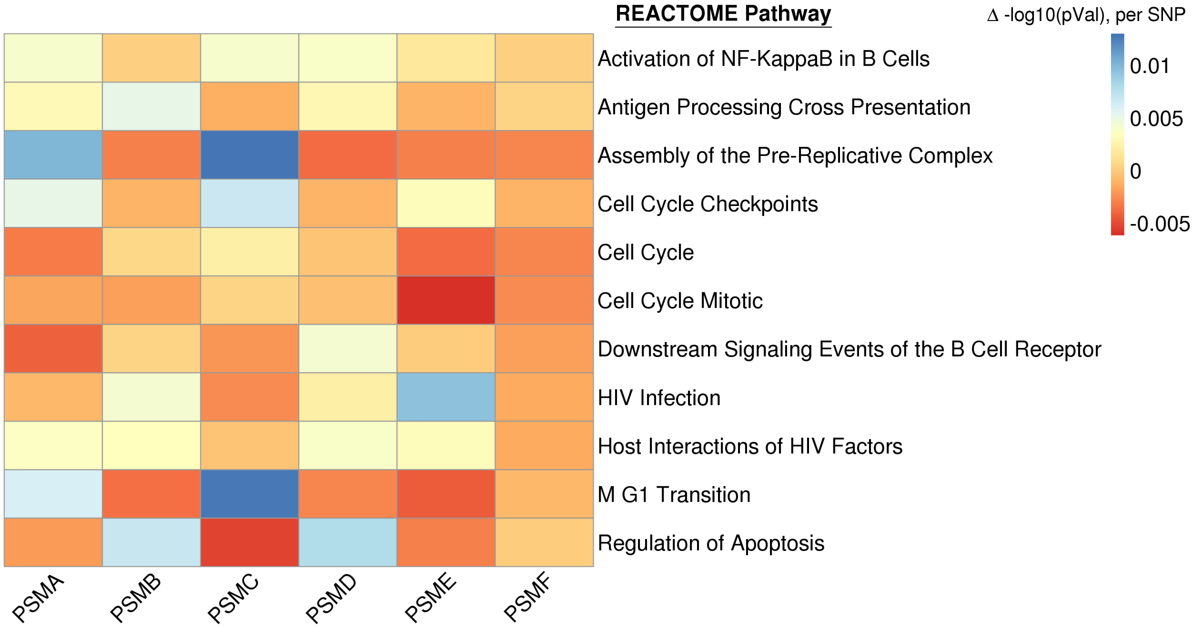


(a) $\Delta$ in MAPIT-R -$\log_{10}$ $p$-value per SNP

| Gene Family | # SNPs | REACTOME Pathway | # SNPs | $p$-Value |
|---|---|---|---|---|
| PSMA | 29 | Activation of NF-KappaB in B Cells | 618 | $9.98\times10^{-1}$ |
| PSMB | 85 | Antigen Processing Cross Presentation | 977 | $4.48\times10^{-1}$ |
| PSMC | 25 | Assembly of the Pre-Replicative Complex | 427 | $5.02\times10^{-1}$ |
| PSMD | 78 | Cell Cycle Checkpoints | 909 | $8.04\times10^{-1}$ |
| PSME | 21 | Cell Cycle | 3361 | $1.87\times10^{-1}$ |
| PSMF | 21 | Cell Cycle Mitotic | 2656 | $2.32\times10^{-1}$ |
| | | Downstream Signaling Events of the B Cell Receptor | 1037 | $5.53\times10^{-1}$ |
| | | HIV Infection | 1836 | $9.80\times10^{-2}$ |
| | | Host Interactions of HIV Factors | 1278 | $3.79\times10^{-1}$ |
| | | M G1 Transition | 590 | $4.65\times10^{-1}$ |
| | | Regulation of Apoptosis | 754 | $5.61\times10^{-1}$ |

(b) Number of SNPs in each Proteasome Gene Family and REACTOME Pathway

**Supplementary Figure 19. Results from applying a "leave-one-out" approach to MAPIT-R with proteasome gene families in body mass index (BMI) within the Indian subgroup in the UK Biobank. (a)** The heatmap shows the change in original MAPIT-R -$\log_{10}$ $p$-value for different REACTOME pathways when each proteasome gene family is removed one at a time in a "leave-one-out" manner. The $x$-axis shows each proteasome gene family and the $y$-axis lists each REACTOME pathway. Each column has been scaled by the number of SNPs present in the given gene family and, as a result, the heatmap specifically shows the -$\log_{10}$ $p$-value change ($\Delta$ in legend) per SNP. Raw MAPIT-R $p$-values underlying the heatmap can be found in Supplementary Table 10. **(b)** The table shows the number of SNPs present in each proteasome gene family (left), as well as the number of SNPs present in each REACTOME pathway (right). The original MAPIT-R $p$-values are also shown for each pathway (right).

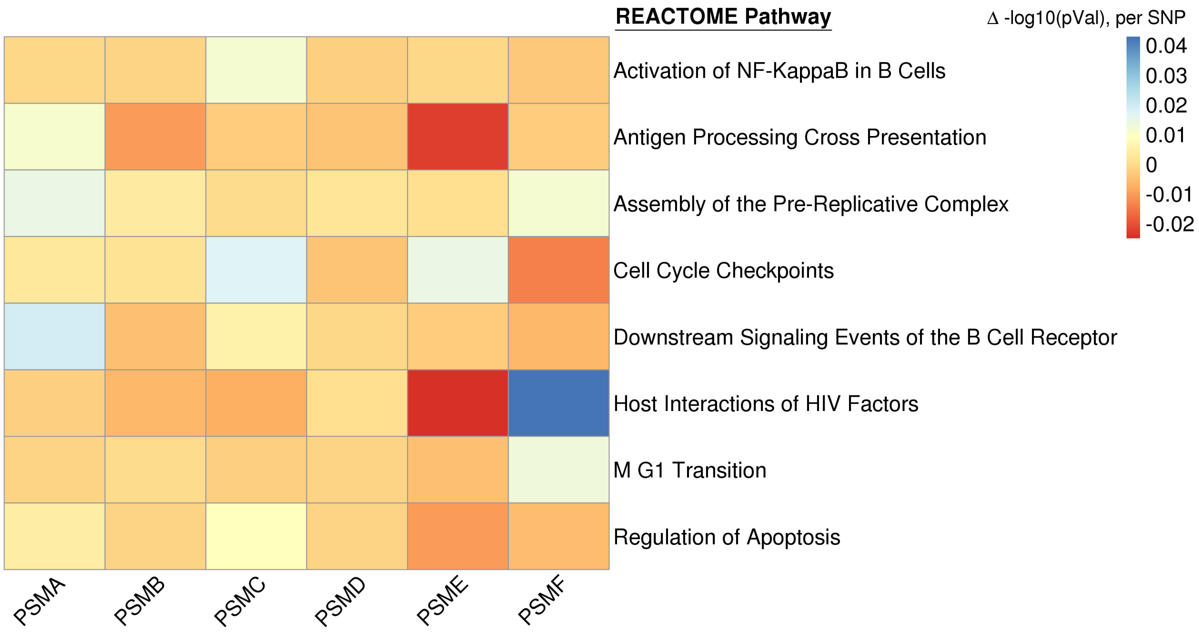


(a) Δ in MAPIT-R -$\log_{10}$ $p$-value per SNP

| Gene Family | # SNPs | REACTOME Pathway | # SNPs | $p$-Value |
|---|---|---|---|---|
| PSMA | 30 | Activation of NF-KappaB in B Cells | 643 | $4.61\times10^{-1}$ |
| PSMB | 90 | Antigen Processing Cross Presentation | 1036 | $8.74\times10^{-3}$ |
| PSMC | 24 | Assembly of the Pre-Replicative Complex | 444 | $9.73\times10^{-1}$ |
| PSMD | 86 | Cell Cycle Checkpoints | 940 | $1.63\times10^{-1}$ |
| PSME | 21 | Downstream Signaling Events of the B Cell Receptor | 1073 | $6.57\times10^{-2}$ |
| PSMF | 22 | Host Interactions of HIV Factors | 1315 | $1.00\times10^{-1}$ |
| | | M G1 Transition | 612 | $6.66\times10^{-1}$ |
| | | Regulation of Apoptosis | 774 | $5.16\times10^{-2}$ |

(b) Number of SNPs in each Proteasome Gene Family and REACTOME Pathway

**Supplementary Figure 20. Results from applying a "leave-one-out" approach to MAPIT-R with proteasome gene families in body mass index (BMI) within the Pakistani subgroup in the UK Biobank. (a)** The heatmap shows the change in original MAPIT-R -$\log_{10}$ $p$-value for different REACTOME pathways when each proteasome gene family is removed one at a time in a "leave-one-out" manner. The $x$-axis shows each proteasome gene family and the $y$-axis lists each REACTOME pathway. Each column has been scaled by the number of SNPs present in the given gene family and, as a result, the heatmap specifically shows the -$\log_{10}$ $p$-value change (Δ in legend) per SNP. Raw MAPIT-R $p$-values underlying the heatmap can be found in Supplementary Table 10. **(b)** The table shows the number of SNPs present in each proteasome gene family (left), as well as the number of SNPs present in each REACTOME pathway (right). The original MAPIT-R $p$-values are also shown for each pathway (right).

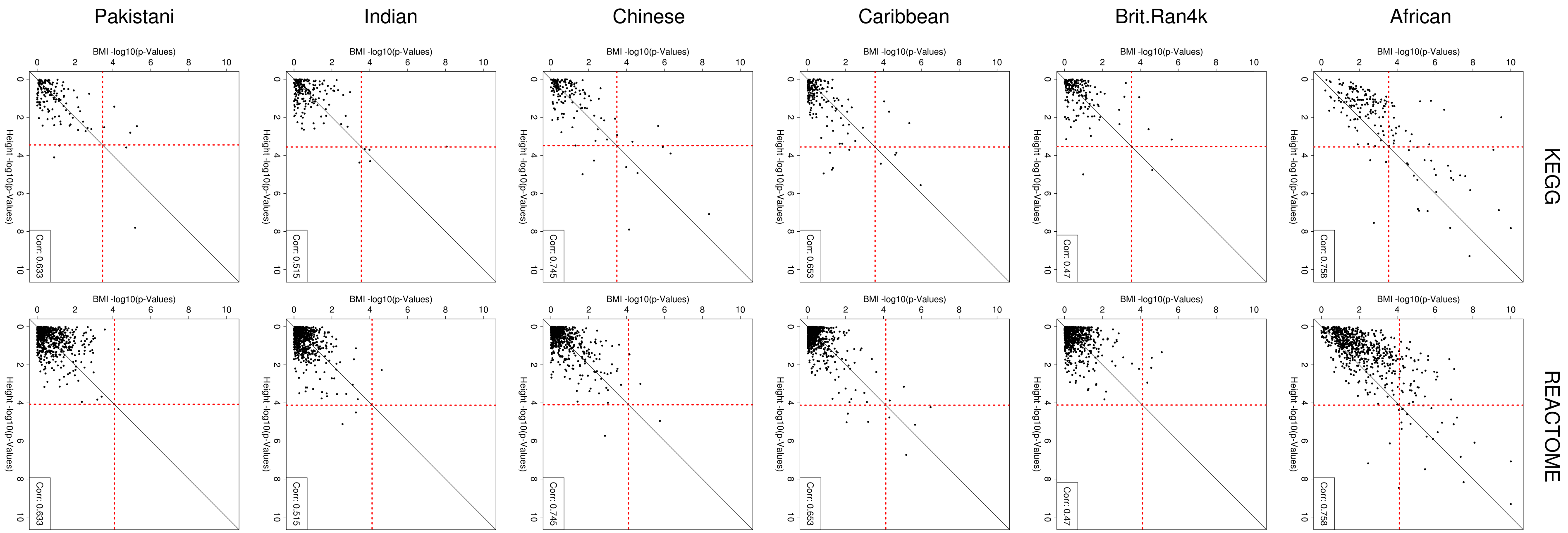


**Supplementary Figure 21. Scatterplots comparing the MAPIT-R $p$-values using (top) KEGG and (bottom) REACTOME pathway annotations in height and body mass index (BMI) within all of the different ancestry-specific subgroups in the UK Biobank.** Here, subgroups in the UK Biobank included individuals based on their self-identified ancestries: "African", "British", "Caribbean", "Chinese", "Indian", and "Pakistani", (ordered here from right-to-left). For each plot, the $x$-axis shows the -$\log_{10}$ transformed MAPIT-R $p$-value for height, while the $y$-axis shows the same results for BMI. The red horizontal and vertical dashed lines are marked at the Bonferroni-corrected $p$-value thresholds for genome-wide significance in each pathway-phenotype combination (see Supplementary Table 1). Pathways in the top right quadrant have significant marginal epistatic effects in both traits; while, points in the bottom right and top left quadrants are pathways that are uniquely enriched in height or BMI, respectively. Within each plot we also list the Pearson correlation coefficient describing the similarity of marginally epistatic enriched pathways in both traits for each database-subgroup combination.

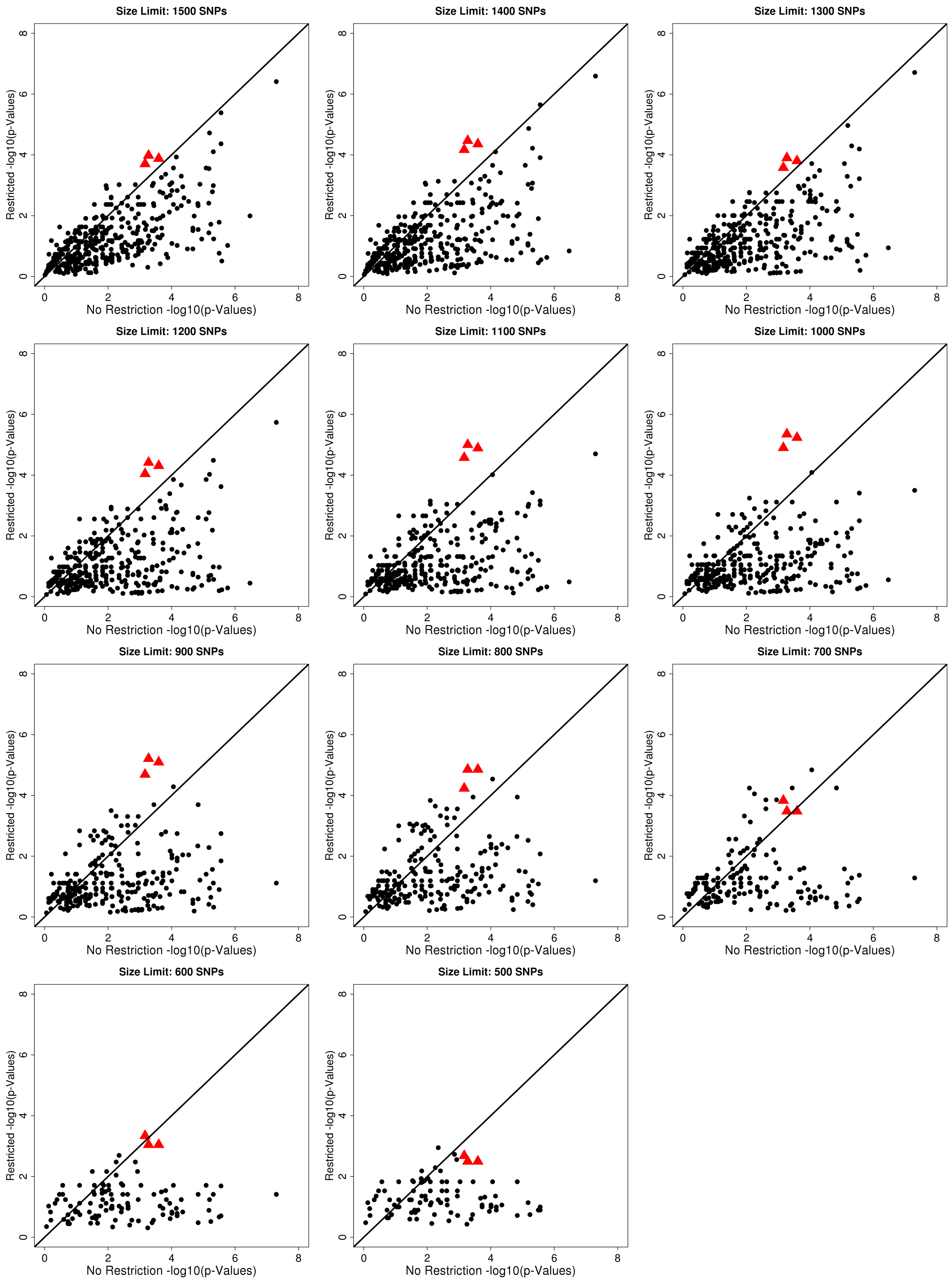


**Supplementary Figure 22. Impact of pathway size thresholds on hypergeometric enrichment analyses.** Here, we show multiple comparisons of size-restricted versus unrestricted hypergeometric enrichment analyses using incrementally decreasing pathway size thresholds. Size thresholds range from $T = 500$ SNPs to 1,500 SNPs in 100-SNP increments. In each plot, the $x$-axis is the unrestricted hypergeometric $p$-value for each gene, while the $y$-axis is the size-restricted hypergeometric $p$-value for each gene. Red triangles represent results for each gene belonging to a proteasome gene family: *PSMA*, *PSMB*, *PSMC*, *PSMD*, *PSME*, and *PSMF*. Note that there are only three triangles since most proteasome genes are assigned to the same pathways.

### 4 Supplementary Tables

| Subgroups | Ancestry | Individuals | SNPs | KEGG | REACTOME |
|---|---|---|---|---|---|
| **Original** | African | 3111 | 374466 | 180 | 658 |
| | British (4k, Rep. 1) | 3848 | 600006 | 173 | 650 |
| | Caribbean | 3833 | 410017 | 181 | 661 |
| | Chinese | 1448 | 345221 | 153 | 626 |
| | Indian | 5077 | 505854 | 181 | 662 |
| | Pakistani | 1581 | 516806 | 141 | 596 |
| **British** | British (4k, Rep. 2) | 3869 | 599381 | 173 | 650 |
| | British (4k, Rep. 3) | 3836 | 600654 | 173 | 649 |
| | British (4k, Rep. 4) | 3838 | 599829 | 173 | 650 |
| | British (4k, Rep. 5) | 3853 | 599442 | 173 | 650 |
| | British (10k, Rep. 1) | 9603 | 597298 | 186 | 669 |
| | British (10k, Rep. 2) | 9628 | 597577 | 186 | 669 |
| | British (10k, Rep. 3) | 9636 | 597486 | 186 | 669 |
| | British (10k, Rep. 4) | 9593 | 597369 | 186 | 669 |
| | British (10k, Rep. 5) | 9596 | 597507 | 186 | 669 |

**Supplementary Table 1. Tabulation of the number of subgroups, individuals, SNPs, and pathways analyzed in this study**. Here, the `Subgroups` category "Original" refers to the multiple global human ancestries from the UK Biobank that were initially analyzed at the beginning of this study, while the `Subgroups` category "British" refers specifically to the different replicates that were created by taking random independent subsamples of the full British cohort in the UK Biobank. The number of individuals and SNPs in the second and third columns are those post quality control (see Supplementary Note for details and Supplementary Table 2 for number of individuals dropped at each relevant quality control processing step), and the number of KEGG and REACTOME pathways analyzed within each subgroup are shown in the fourth and fifth columns. Note that for the "British replicate" subgroups, each dataset initially began with either 4,000 or 10,000 individuals before quality control. Additionally, pathway numbers across British replicates vary slightly due to some replicates not containing enough individuals to analyze pathways with the most SNPs; this leads to those largest pathways being dropped from the analysis.

| Subgroups | Ancestry | Initial Numbers | Missingness Drops | Relatedness Drops | UKBiobank Drops | PCA Drops |
|---|---|---|---|---|---|---|
| **Original** | African | 3205 | 3202 | 3117 | 3116 | 3111 |
| | British (4k, Rep. 1) | 3889 | 3888 | 3878 | 3866 | 3848 |
| | Caribbean | 4299 | 4296 | 3861 | 3854 | 3833 |
| | Chinese | 1504 | 1502 | 1454 | 1454 | 1448 |
| | Indian | 5716 | 5715 | 5261 | 5203 | 5077 |
| | Pakistani | 1748 | 1746 | 1609 | 1604 | 1581 |
| **British** | British (4k, Rep. 2) | 3907 | 3906 | 3900 | 3890 | 3869 |
| | British (4k, Rep. 3) | 3884 | 3881 | 3875 | 3859 | 3836 |
| | British (4k, Rep. 4) | 3882 | 3879 | 3868 | 3855 | 3838 |
| | British (4k, Rep. 5) | 3894 | 3891 | 3886 | 3871 | 3853 |
| | British (10k, Rep. 1) | 9733 | 9726 | 9683 | 9639 | 9603 |
| | British (10k, Rep. 2) | 9747 | 9745 | 9698 | 9669 | 9628 |
| | British (10k, Rep. 3) | 9740 | 9736 | 9692 | 9671 | 9636 |
| | British (10k, Rep. 4) | 9750 | 9748 | 9697 | 9665 | 9593 |
| | British (10k, Rep. 5) | 9739 | 9731 | 9687 | 9654 | 9596 |

**Supplementary Table 2. Tabulation of the number of individuals dropped per subgroup after each quality control step**. Here, the `Subgroups` category "Original" refers to the multiple global human ancestries from the UK Biobank that were initially analyzed at the beginning of this study, while the `Subgroups` category "British" refers specifically to the different replicates that were created by taking random independent subsamples of the full British cohort in the UK Biobank. The third column shows the number of individuals each subgroup began with prior to any quality control steps. Note that British replicates were initially created by extracting individual IDs from the associated UK Biobank individual information file. However, not all individuals present in this file are also present in the genotype files, so a set of individuals from the original groups of 4,000 and 10,000 were initially dropped at this point. The fourth column shows the remaining sample sizes after dropping individuals due to genotype missingness being greater than 5%. The fifth column shows the remaining sample sizes after dropping a random individual from every pair of individuals that were 3$^{\text{rd}}$ degree relatives or more based on the KING relatedness values provided with the UK Biobank dataset. The sixth column shows the remaining sample sizes after dropping individuals due to being flagged by the following three UK Biobank dataset quality control categories: `het.missing.outliers`, `putative.sex.chromosome.aneuploidy`, and `excess.relatives`. The seventh column shows the remaining sample sizes after dropping individuals due to being principal component analysis (PCA) outliers based on the top six local principal components (PCs). We refer to conducting PCA on each subgroup separately as "local" PCA to help distinguish from the alternative setup of conducting PCA on the entire dataset jointly, which we refer to as "global" PCA. Note that global PCs were only used to create the PCA plots shown in Supplementary Figure 1, while local PCs were used for quality control and to adjust for population structure in the MAPIT-R analyses. For other final dataset statistics see Supplementary Table 1 and for other details regarding dataset quality control steps see Supplementary Note.

**Supplementary Table 3. List of MAPIT-R significant pathways genome-wide for standing height and body mass index (BMI) across the different ancestry-specific subgroups in the UK Biobank.** Here, subgroups in the UK Biobank included individuals based on their self-identified ancestries: "African", "British", "Caribbean", "Chinese", "Indian", and "Pakistani" (ordered here from top-to-bottom). Phenotype and pathway database combinations include: (*i*) KEGG+Height, (*ii*) KEGG+BMI, (*iii*) REACTOME+Height, and (*iv*) REACTOME+BMI. Genome-wide significance was determined by using Bonferroni-corrected $p$-value thresholds based on the number of pathways tested in each database-phenotype-subgroup combination with significance value $\alpha = 0.05$ (see Supplementary Table 1). (XLSX)

| Database (Trait) | Subgroup | # Pathways | $\alpha$ = Bonferroni Correction | | | $\alpha = 0.001$ Thresh. | | $\alpha = 0.01$ Thresh. | |
|---|---|---|---|---|---|---|---|---|---|
| | | | Thresh. | # FPs | FDR (%) | # FPs | FDR (%) | # FPs | FDR (%) |
| **KEGG (Height)** | African | 1800 | $2.78\times10^{-4}$ | 0 | 0.000 | 1 | 0.056 | 10 | 0.556 |
| | British (4k) | 1730 | $2.89\times10^{-4}$ | 0 | 0.000 | 0 | 0.000 | 13 | 0.751 |
| | Caribbean | 1810 | $2.76\times10^{-4}$ | 1 | 0.055 | 1 | 0.055 | 5 | 0.276 |
| | Chinese | 1530 | $3.27\times10^{-4}$ | 0 | 0.000 | 3 | 0.196 | 24 | 1.569 |
| | Indian | 1810 | $2.76\times10^{-4}$ | 1 | 0.055 | 1 | 0.055 | 20 | 1.105 |
| | Pakistani | 1410 | $3.55\times10^{-4}$ | 0 | 0.000 | 2 | 0.142 | 17 | 1.206 |
| **KEGG (BMI)** | African | 1800 | $2.778\times10^{-4}$ | 0 | 0.000 | 1 | 0.056 | 16 | 0.889 |
| | Caribbean | 1810 | $2.762\times10^{-4}$ | 0 | 0.000 | 1 | 0.055 | 25 | 1.381 |
| | Chinese | 1530 | $3.268\times10^{-4}$ | 0 | 0.000 | 1 | 0.065 | 24 | 1.569 |
| | Indian | 1810 | $2.762\times10^{-4}$ | 1 | 0.055 | 2 | 0.110 | 20 | 1.105 |
| | Irish | 1860 | $2.688\times10^{-4}$ | 0 | 0.000 | 0 | 0.000 | 14 | 0.753 |
| | Pakistani | 1410 | $3.546\times10^{-4}$ | 0 | 0.000 | 2 | 0.142 | 15 | 1.064 |
| **REACTOME (Height)** | African | 6580 | $7.599\times10^{-5}$ | 0 | 0.000 | 6 | 0.091 | 47 | 0.714 |
| | Caribbean | 6610 | $7.564\times10^{-5}$ | 0 | 0.000 | 2 | 0.030 | 22 | 0.333 |
| | Chinese | 6260 | $7.987\times10^{-5}$ | 0 | 0.000 | 13 | 0.208 | 85 | 1.358 |
| | Indian | 6620 | $7.553\times10^{-5}$ | 1 | 0.015 | 5 | 0.076 | 90 | 1.360 |
| | Irish | 6710 | $7.452\times10^{-5}$ | 0 | 0.000 | 1 | 0.015 | 55 | 0.820 |
| | Pakistani | 5960 | $8.389\times10^{-5}$ | 0 | 0.000 | 3 | 0.050 | 63 | 1.057 |
| **REACTOME (BMI)** | African | 6580 | $7.60\times10^{-5}$ | 1 | 0.015 | 4 | 0.061 | 39 | 0.593 |
| | British (4k) | 6490 | $7.70\times10^{-5}$ | 1 | 0.015 | 4 | 0.062 | 52 | 0.801 |
| | Caribbean | 6610 | $7.56\times10^{-5}$ | 0 | 0.000 | 13 | 0.197 | 97 | 1.467 |
| | Chinese | 6260 | $7.99\times10^{-5}$ | 0 | 0.000 | 6 | 0.096 | 49 | 0.783 |
| | Indian | 6620 | $7.55\times10^{-5}$ | 0 | 0.000 | 3 | 0.045 | 66 | 0.997 |
| | Pakistani | 5960 | $8.39\times10^{-5}$ | 0 | 0.000 | 4 | 0.067 | 45 | 0.755 |

**Supplementary Table 4. MAPIT-R false discovery rates at different significance thresholds using permuted phenotypes across the different ancestry-specific subgroups in the UK Biobank.** Here, we run MAPIT-R after conducting a single random permutation of either height or BMI measurements. Note that traits were independently permuted within each population subgroup ten different times. Subgroups in the UK Biobank included individuals based on their self-identified ancestries: "African", "British", "Caribbean", "Chinese", "Indian", and "Pakistani" (ordered here from top-to-bottom in each section). Phenotype and pathway database combinations include: (*i*) KEGG+Height, (*ii*) KEGG+BMI, (*iii*) REACTOME+Height, and (*iv*) REACTOME+BMI. Genome-wide significance at a Bonferroni-corrected $p$-value threshold was based on the number of pathways tested in each database-phenotype-subgroup combination with significance value set to $\alpha = 0.05$. The second column lists the total number of pathways that were tested across each of the ten phenotype permutations. # FPs denotes the number of false positives at that threshold and FDR reports the corresponding false discovery rate (listed as percentages). The remaining columns list the same information but at significance levels $\alpha = 0.001$ and 0.01, respectively.

| Population Comparison | Enriched Pathways |
|---|---|
| **African and Caribbean** | KEGG_ARRHYTHMOGENIC_RIGHT_VENTRICULAR_CARDIOMYOPATHY_ARVC |
| | KEGG_AXON_GUIDANCE |
| | KEGG_CHEMOKINE_SIGNALING_PATHWAY |
| | KEGG_HYPERTROPHIC_CARDIOMYOPATHY_HCM |
| | KEGG_NATURAL_KILLER_CELL_MEDIATED_CYTOTOXICITY |
| | KEGG_VASCULAR_SMOOTH_MUSCLE_CONTRACTION |
| **African and Chinese** | KEGG_ALLOGRAFT_REJECTION |
| | KEGG_ANTIGEN_PROCESSING_AND_PRESENTATION |
| | KEGG_GRAFT_VERSUS_HOST_DISEASE |
| | KEGG_SYSTEMIC_LUPUS_ERYTHEMATOSUS |
| | KEGG_TYPE_I_DIABETES_MELLITUS |

**Supplementary Table 5. List of overlap between the MAPIT-R enriched pathways genome-wide for standing height and body mass index (BMI) across the different ancestry-specific subgroups in the UK Biobank.** Pathway annotations were determined according to the KEGG database. Genome-wide significance was determined by using Bonferroni-corrected $p$-value thresholds based on the number of pathways tested in each database-phenotype-subgroup combination with significance value $\alpha = 0.05$ (see Supplementary Table 1).

| Population Comparison | Frequency | Genes in Enriched Pathways |
|---|---|---|
| **African and Caribbean** | 4 | *MAPK1*, *MAPK3* |
| | 3 | *ROCK2*, *ROCK1*, *RHOA*, *RAF1*, *RAC2*, *RAC1*, *PRKCB*, *PAK1*, *NRAS*, *MAP2K1*, *KRAS*, *ITGB1*, *HRAS*, *CACNA1S*, *CACNA1D*, *CACNA1C*, *BRAF* |
| **African and Chinese** | 5 | *HLA-DRB1*, *HLA-DRA*, *HLA-DQB1*, *HLA-DQA2*, *HLA-DQA1*, *HLA-DPB1*, *HLA-DPA1*, *HLA-DOB*, *HLA-DOA*, *HLA-DMB*, *HLA-DM* |
| | 4 | *TNF*, *IFNG*, *HLA-G*, *HLA-F*, *HLA-E*, *HLA-C*, *HLA-B*, *HLA-A*, *CD86*, *CD80*, *CD28* |
| | 3 | *PRF1*, *IL2*, *GZMB*, *FASLG*, *FAS* |

**Supplementary Table 6. Genes that are annotated for multiple MAPIT-R enriched pathways in both standing height and body mass index (BMI) for different ancestry-specific subgroups in the UK Biobank.** Genome-wide significance was determined by using Bonferroni-corrected $p$-value thresholds based on the number of pathways tested in each database-phenotype-subgroup combination with significance value $\alpha = 0.05$ (see Supplementary Table 1). The second column records the number of enriched pathways that each gene belongs to, according to the KEGG database.

| Trait Comparison | Frequency | Genes in Enriched Pathways |
|---|---|---|
| **Height vs. BMI** | 4 | *PIK3R5*, *PIK3R3*, *PIK3R2*, *PIK3R1*, *PIK3CG*, *PIK3CD*, *PIK3CB*, *PIK3CA*, *AKT3*, *AKT2*, *AKT1* |
| | 3 | *SOS2*, *SOS1*, *RAF1*, *PLCG1*, *NRAS*, *MAPK3*, *MAPK1*, *MAP2K2*, *MAP2K1*, *KRAS*, *HRAS*, *GRB2*, *CDK4* |
| | 2 | *TP53*, *TGFA*, *RXRG*, *RXRB*, *RXRA*, *RELA*, *RB1*, *RARB*, *PTK2*, *PRKCG*, *PRKCB*, *PRKCA*, *PLCG2*, *PDPK1*, *PAK6*, *PAK4*, *PAK2*, *PAK1*, *NFKBIA*, *NFKB1*, *NCK2*, *NCK1*, *MYC*, *MAPK9*, *MAP2K7*, *JUN*, *IKBKB*, *GSK3B*, *FHIT*, *ERBB2*, *EGFR*, *EGF*, *E2F3*, *E2F2*, *E2F1*, *CHUK*, *CDKN1B*, *CDK6*, *CCND1*, *CBLC*, *CBLB*, *CBL*, *CASP9*, *BRAF*, *BAD* |

**Supplementary Table 7. Genes that are annotated for multiple enriched pathways in both standing height and body mass index (BMI) within the African subgroup from the UK Biobank.** Here, we compare genes that are present across multiple of the pathways highlighted in blue in Figure 3; these are specific pathways that have more significant MAPIT-R $p$-values in BMI than in height. Genome-wide significance was determined by using Bonferroni-corrected $p$-value thresholds based on the number of pathways tested in each database-phenotype-subgroup combination with significance value $\alpha = 0.05$ (see Supplementary Table 1). The second column records the number of enriched pathways that each gene belongs to, according to the KEGG database.

| | | | $\alpha$ = Bonferroni Correction | | | $\alpha = 0.001$ Thresh. | | $\alpha = 0.01$ Thresh. | |
|---|---|---|---|---|---|---|---|---|---|
| **Database (Trait)** | **Subgroup** | **# Pathways** | **Thresh.** | **# FPs** | **FDR (%)** | **# FPs** | **FDR (%)** | **# FPs** | **FDR (%)** |
| **KEGG (Height)** | British (4k, Rep. 1) | 1730 | $2.89\times10^{-4}$ | 0 | 0.000 | 0 | 0.000 | 13 | 0.751 |
| | British (4k, Rep. 2) | 1730 | $2.89\times10^{-4}$ | 0 | 0.000 | 1 | 0.058 | 18 | 1.040 |
| | British (4k, Rep. 3) | 1730 | $2.89\times10^{-4}$ | 1 | 0.058 | 1 | 0.058 | 20 | 1.156 |
| | British (4k, Rep. 4) | 1730 | $2.89\times10^{-4}$ | 0 | 0.000 | 4 | 0.231 | 23 | 1.329 |
| | British (4k, Rep. 5) | 1730 | $2.89\times10^{-4}$ | 0 | 0.000 | 2 | 0.116 | 22 | 1.272 |
| | British (10k, Rep. 1) | 1860 | $2.69\times10^{-4}$ | 0 | 0.000 | 1 | 0.054 | 26 | 1.398 |
| | British (10k, Rep. 2) | 1860 | $2.69\times10^{-4}$ | 0 | 0.000 | 1 | 0.054 | 12 | 0.645 |
| | British (10k, Rep. 3) | 1860 | $2.69\times10^{-4}$ | 2 | 0.108 | 2 | 0.108 | 21 | 1.129 |
| | British (10k, Rep. 4) | 1860 | $2.69\times10^{-4}$ | 1 | 0.054 | 2 | 0.108 | 16 | 0.860 |
| | British (10k, Rep. 5) | 1860 | $2.69\times10^{-4}$ | 0 | 0.000 | 0 | 0.000 | 12 | 0.645 |
| **KEGG (BMI)** | British (4k, Rep. 1) | 1730 | $2.89\times10^{-4}$ | 1 | 0.058 | 2 | 0.116 | 14 | 0.809 |
| | British (4k, Rep. 2) | 1730 | $2.89\times10^{-4}$ | 0 | 0.000 | 1 | 0.058 | 23 | 1.329 |
| | British (4k, Rep. 3) | 1730 | $2.89\times10^{-4}$ | 0 | 0.000 | 3 | 0.173 | 21 | 1.214 |
| | British (4k, Rep. 4) | 1730 | $2.89\times10^{-4}$ | 1 | 0.058 | 5 | 0.289 | 21 | 1.214 |
| | British (4k, Rep. 5) | 1730 | $2.89\times10^{-4}$ | 2 | 0.116 | 7 | 0.405 | 23 | 1.329 |
| | British (10k, Rep. 1) | 1860 | $2.69\times10^{-4}$ | 0 | 0.000 | 1 | 0.054 | 25 | 1.344 |
| | British (10k, Rep. 2) | 1860 | $2.69\times10^{-4}$ | 2 | 0.108 | 4 | 0.215 | 22 | 1.183 |
| | British (10k, Rep. 3) | 1860 | $2.69\times10^{-4}$ | 0 | 0.000 | 2 | 0.108 | 20 | 1.075 |
| | British (10k, Rep. 4) | 1860 | $2.69\times10^{-4}$ | 1 | 0.054 | 2 | 0.108 | 23 | 1.237 |
| | British (10k, Rep. 5) | 1860 | $2.69\times10^{-4}$ | 0 | 0.000 | 0 | 0.000 | 20 | 1.075 |

**Supplementary Table 8. MAPIT-R false discovery rates at different significance thresholds using KEGG pathway annotations and permuted phenotypes across the British replicate subgroups in the UK Biobank.** Here, we run MAPIT-R after conducting a single random permutation of either height or body mass index (BMI) measurements. Note that traits were independently permuted within each population subgroup ten different times. The random subgroups were created by subsampling either $N$ = 4,000 or 10,000 individuals from the full British cohort. Genome-wide significance at a Bonferroni-corrected $p$-value threshold was based on the number of pathways tested in each database-phenotype-subgroup combination with significance value set to $\alpha = 0.05$. The second column lists the total number of pathways that were tested across each of the ten phenotype permutations. # FPs denotes the number of false positives at that threshold and FDR reports the corresponding false discovery rate (listed as percentages). The remaining columns list the same information but at significance levels $\alpha$ = 0.001 and 0.01, respectively. See Supplementary Figures 7-12 and Supplementary Table 9 for other analyses using the 4,000 and 10,000 British replicate subgroups.

| | | | $\alpha$ = Bonferroni Correction | | | $\alpha = 0.001$ Thresh. | | $\alpha = 0.01$ Thresh. | |
|---|---|---|---|---|---|---|---|---|---|
| **Database (Trait)** | **Subgroup** | **# Pathways** | **Thresh.** | **# FPs** | **FDR (%)** | **# FPs** | **FDR (%)** | **# FPs** | **FDR (%)** |
| **REACTOME (Height)** | British (4k, Rep. 1) | 6500 | $7.692{\times}10^{-5}$ | 0 | 0.000 | 9 | 0.138 | 69 | 1.062 |
| | British (4k, Rep. 2) | 6500 | $7.692{\times}10^{-5}$ | 0 | 0.000 | 4 | 0.062 | 61 | 0.938 |
| | British (4k, Rep. 3) | 6490 | $7.704{\times}10^{-5}$ | 1 | 0.015 | 8 | 0.123 | 59 | 0.909 |
| | British (4k, Rep. 4) | 6500 | $7.692{\times}10^{-5}$ | 0 | 0.000 | 5 | 0.077 | 69 | 1.062 |
| | British (4k, Rep. 5) | 6500 | $7.692{\times}10^{-5}$ | 0 | 0.000 | 6 | 0.092 | 53 | 0.815 |
| | British (10k, Rep. 1) | 6690 | $7.474{\times}10^{-5}$ | 0 | 0.000 | 4 | 0.060 | 55 | 0.822 |
| | British (10k, Rep. 2) | 6690 | $7.474{\times}10^{-5}$ | 0 | 0.000 | 4 | 0.060 | 52 | 0.777 |
| | British (10k, Rep. 3) | 6690 | $7.474{\times}10^{-5}$ | 2 | 0.030 | 11 | 0.164 | 60 | 0.897 |
| | British (10k, Rep. 4) | 6690 | $7.474{\times}10^{-5}$ | 1 | 0.015 | 4 | 0.060 | 69 | 1.031 |
| | British (10k, Rep. 5) | 6690 | $7.474{\times}10^{-5}$ | 0 | 0.000 | 4 | 0.060 | 60 | 0.897 |
| **REACTOME (BMI)** | British (4k, Rep. 1) | 6490 | $7.704{\times}10^{-5}$ | 1 | 0.015 | 4 | 0.062 | 52 | 0.801 |
| | British (4k, Rep. 2) | 6500 | $7.692{\times}10^{-5}$ | 1 | 0.015 | 14 | 0.215 | 63 | 0.969 |
| | British (4k, Rep. 3) | 6490 | $7.704{\times}10^{-5}$ | 0 | 0.000 | 5 | 0.077 | 79 | 1.217 |
| | British (4k, Rep. 4) | 6500 | $7.692{\times}10^{-5}$ | 2 | 0.031 | 6 | 0.092 | 61 | 0.938 |
| | British (4k, Rep. 5) | 6500 | $7.692{\times}10^{-5}$ | 0 | 0.000 | 7 | 0.108 | 73 | 1.123 |
| | British (10k, Rep. 1) | 6690 | $7.474{\times}10^{-5}$ | 2 | 0.030 | 10 | 0.149 | 73 | 1.091 |
| | British (10k, Rep. 2) | 6690 | $7.474{\times}10^{-5}$ | 0 | 0.000 | 3 | 0.045 | 42 | 0.628 |
| | British (10k, Rep. 3) | 6690 | $7.474{\times}10^{-5}$ | 1 | 0.015 | 10 | 0.149 | 74 | 1.106 |
| | British (10k, Rep. 4) | 6690 | $7.474{\times}10^{-5}$ | 0 | 0.000 | 7 | 0.105 | 65 | 0.972 |
| | British (10k, Rep. 4) | 6690 | $7.474{\times}10^{-5}$ | 0 | 0.000 | 5 | 0.075 | 57 | 0.852 |

**Supplementary Table 9. MAPIT-R false discovery rates at different significance thresholds using REACTOME pathway annotations and permuted phenotypes across the British replicate subgroups in the UK Biobank.** Here, we run MAPIT-R after conducting a single random permutation of either height or body mass index (BMI) measurements. Note that traits were independently permuted within each population subgroup ten different times. The random subgroups were created by subsampling either $N$ = 4,000 or 10,000 individuals from the full British cohort. Genome-wide significance at a Bonferroni-corrected $p$-value threshold was based on the number of pathways tested in each database-phenotype-subgroup combination with significance value set to $\alpha = 0.05$. The second column lists the total number of pathways that were tested across each of the ten phenotype permutations. # FPs denotes the number of false positives at that threshold and FDR reports the corresponding false discovery rate (listed as percentages). The remaining columns list the same information but at significance levels $\alpha$ = 0.001 and 0.01, respectively. See Supplementary Figures 7-12 and Supplementary Table 8 for other analyses using the 4,000 and 10,000 British replicate subgroups.

**Supplementary Table 10. Results from applying a "leave-one-out" approach to MAPIT-R with proteasome gene families in body mass index (BMI) across the different ancestry-specific subgroups in the UK Biobank.** Here, pathway annotations were determined according to the REACTOME database. Subgroups in the UK Biobank included individuals based on their self-identified ancestries: "African", "British", "Caribbean", "Chinese", "Indian", and "Pakistani" (ordered here from a-f). The proteasome gene families analyzed included: *PSMA*, *PSMB*, *PSMC*, *PSMD*, *PSME*, and *PSMF*. These are the raw MAPIT-R $p$-values underlying Figure 5 in the main text and Supplementary Figures 15-20. Note that for some subgroups not all of the original pathways from Figure 5 were analyzable (generally due to number of SNPs being far greater than the number of individuals). (XLSX)

**Supplementary Table 11. Results from hypergeometric tests performed for body mass index (BMI) using both the original and size-restricted pathway annotations within the African subgroup in the UK Biobank.** Here, pathway annotations were determined according to both the KEGG and REACTOME databases. Listed are the -$\log_{10}$ transformed $p$-values from the hypergeometric tests. Differences in the -$\log_{10}$ $p$-values between the original and size-restricted hypergeometric analyses are also shown per gene. Attached Excel sheets contain lists of the hypergeometric -$\log_{10}$ $p$-values for both the original and size-restricted gene count analyses per gene in both the KEGG and REACTOME-BMI-African combinations. Differences in the -$\log_{10}$ $p$-values between the original and size-restricted hypergeometric analyses are also shown per gene. Plots of these differences in can be seen in Figure 4 in the main text. (XLSX)

**Supplementary Table 12. Genes that are annotated for multiple MAPIT-R enriched pathways in standing height and body mass index (BMI) across the different ancestry-specific subgroups in the UK Biobank.** Subgroups in the UK Biobank included individuals based on their self-identified ancestries: "African", "British", "Caribbean", "Chinese", "Indian", and "Pakistani" (ordered here from top-to-bottom). Phenotype and pathway database combinations include: (*i*) KEGG+Height, (*ii*) KEGG+BMI, (*iii*) REACTOME+Height, and (*iv*) REACTOME+BMI. Genome-wide significance at a Bonferroni-corrected $p$-value threshold was based on the number of pathways tested in each database-phenotype-subgroup combination with significance value set to $\alpha = 0.05$ (see Supplementary Table 1). (XLSX)

**Supplementary Table 13. Genes that are annotated for multiple MAPIT-R enriched pathways (after imposing a size restriction) in standing height and body mass index (BMI) across the different ancestry-specific subgroups in the UK Biobank.** Here, we only consider pathways that contain less than or equal to 1,000 SNPs. Subgroups in the UK Biobank included individuals based on their self-identified ancestries: "African", "British", "Caribbean", "Chinese", "Indian", and "Pakistani" (ordered here from top-to-bottom). Phenotype and pathway database combinations include: (*i*) KEGG+Height, (*ii*) KEGG+BMI, (*iii*) REACTOME+Height, and (*iv*) REACTOME+BMI. Genome-wide significance at a Bonferroni-corrected $p$-value threshold was based on the number of pathways tested in each database-phenotype-subgroup combination with significance value set to $\alpha = 0.05$ (see Supplementary Table 1). (XLSX)

### References


1. Sudlow C, Gallacher J, Allen N, Beral V, Burton P, Danesh J, et al. UK biobank: an open access resource for identifying the causes of a wide range of complex diseases of middle and old age [Journal Article]. PLoS Med. 2015;12(3):e1001779. Available from: https://www.ncbi.nlm.nih.gov/pubmed/25826379.

2. Abraham G, Qiu Y, Inouye M. FlashPCA2: principal component analysis of Biobank-scale genotype datasets [Journal Article]. Bioinformatics. 2017;33(17):2776–2778. Available from: https://www.ncbi.nlm.nih.gov/pubmed/28475694.

3. Das S, Forer L, Schonherr S, Sidore C, Locke AE, Kwong A, et al. Next-generation genotype imputation service and methods [Journal Article]. Nat Genet. 2016;48(10):1284–1287. Available from: https://www.ncbi.nlm.nih.gov/pubmed/27571263.

4. Crawford L, Zeng P, Mukherjee S, Zhou X. Detecting epistasis with the marginal epistasis test in genetic mapping studies of quantitative traits [Journal Article]. PLoS Genet. 2017a;13(7):e1006869. Available from: https://www.ncbi.nlm.nih.gov/pubmed/28746338.
